# Cell-free systems enable mechanistic characterization of genetically encoded RNA strand exchange circuits for programmable protein expression, logic, and biosensing

**DOI:** 10.1101/2025.09.25.678518

**Authors:** Fernanda Piorino, Eugenia Romantseva, Samuel W. Schaffter

## Abstract

Nucleic acid circuits are powerful tools for programming biology, but the design principles for operating these circuits in complex cellular environments remain poorly understood compared to simple *in vitro* settings. Cell-free expression systems (CFES) are uniquely suited to address this challenge, as they can integrate measurements ideal for either test tube or cellular settings. Further, because CFES are open systems, they enable a level of control over component concentration unattainable in cells. Here, we use CFES to characterize genetically encoded RNA circuits that operate *via* toehold-mediated strand exchange (TMSE). These circuits have been extensively characterized in *in vitro* transcription (IVT) reactions and were recently deployed as translational riboregulators in *Escherichia coli*, revealing key differences between the two environments. By systematically modulating the design parameters of these RNA circuits in both purified protein and lysate-based CFES, we elucidate the mechanisms linking TMSE to protein expression. Further, we combine measurements and alterations in CFES composition that are infeasible in cells to investigate interactions between cellular components and RNA circuit components that inform circuit design. Leveraging our mechanistic insights, we demonstrate how a single gate design can achieve multi-level and bidirectional gene regulation, logical operations, and small molecule biosensing. Our results establish a unified set of principles for designing and operating genetically encoded TMSE circuits across IVT, CFES, and bacterial environments, which should catalyze the widespread adoption of this platform for new applications in molecular programming, synthetic biology, and biotechnology.

## INTRODUCTION

A broad goal of molecular programming is to develop interoperable biochemical circuitry that can interface with and control myriad biological systems. Nucleic acid-based molecular circuits have made substantial progress in this direction, demonstrating applications spanning *in vitro* diagnostics to cellular computation. The programmable nature of nucleic acid base pairing interactions lends itself to rational design of biosensors and diagnostics^1–6^,, therapeutics^7^, riboregulators^8–11^, and molecular computing networks^10–12.^. However, achieving this breadth of functionality typically requires a suite of application- and environment-specific designs.

Nucleic acid circuits that operate *via* toehold-mediated strand exchange^15^ (TMSE) have the potential to enable a unifying device design that can achieve broad applicability across environments. TMSE circuits consist of an input strand that interacts with a complementary sequence in a partially double-stranded gate *via* a single-stranded toehold, prompting the release of an output strand that can facilitate new strand exchange reactions (Figure 1A). The modular and composable design of the double-stranded gates, combined with the precise thermodynamic and kinetic control available by modulating toehold lengths, has enabled construction of networks of hundreds of interacting components, each with the same underlying design, that can execute sophisticated information processing tasks^16,17^. Further, because only base pairing interactions are required for information processing, TMSE circuits have the potential to be deployed across broad application environments and organisms^18,19^. These programmable TMSE networks have been developed into *in vitro* diagnostics that classify complex patterns of infection and disease^20,21^ or precisely discriminate nucleic acid biomarkers^22^. These *in vitro* successes were facilitated by mechanistic understanding of the thermodynamics and kinetics of TMSE reactions in well-defined reaction conditions^15,23^.

**FIGURE 1:**
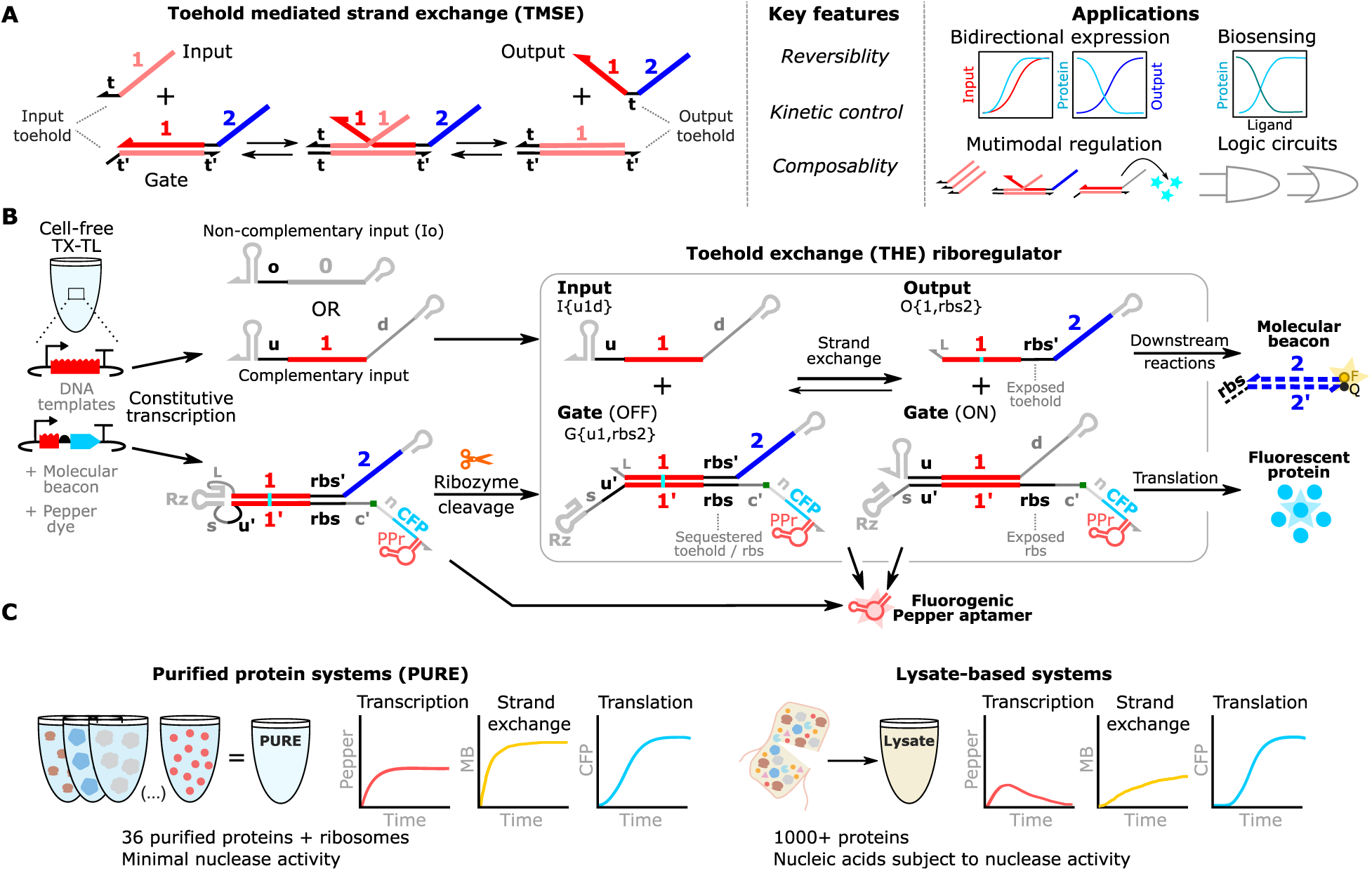
Overview of toehold exchange (THE) riboregulators and associated measurements in cell-free expression systems. (**A**) Schematic of a TMSE reaction along with its key features and applications. (**B**) Schematic of THE riboregulator production, operation, and measurement. The THE riboregulator (gate) and a complementary input are encoded on different plasmids added to a cell-free reaction that supports transcription and translation (TX-TL). Upon transcription, a ribozyme encoded on the gate cleaves the transcript, resulting in a single-stranded region (*u′*) complementary to the toehold on the input strand (*u*), and a double-stranded region including a branch migration domain (*1*) and a sequestered ribosome binding site (*rbs*), thus preventing translation of a downstream cyan fluorescent protein (CFP). The input toehold binds to the gate to initiate strand exchange. The output strand, once dissociated from the gate, can release a fluorophore strand (F) from a previously quenched (Q) molecular beacon complex. Strand exchange also exposes the *rbs*, allowing translation of CFP. The gate also encodes a fluorogenic Pepper RNA aptamer (PPr), which generates a fluorescence signal upon binding to an exogenously added dye. Gate transcripts in the uncleaved, OFF, and ON states contribute to the fluorogenic aptamer signal. A template encoding a non-complementary input, Io, can be added to the reaction to measure leak in the absence of a complementary input, and to maintain the same global transcription levels in reactions containing different concentrations of DNA templates. Additional details on circuit domains and nomenclature are shown in Supplementary Figures 1 and 2. Plasmid schematics are in Supplementary Figure 3. (**C**) Overview of reconstituted (PURE) versus lysate-based cell-free expression systems with expected kinetic trajectories for each measurement.

Without a similar mechanistic understanding in more complex biological systems, realizing the full potential of TMSE circuits in cells remains challenging. This bottleneck stems in part from the difficulty in conducting kinetic measurements and the lack of suitable tools for directly measuring strand exchange in cells. Further, the precise control of relative ratios of nucleic acid components necessary for mechanistic characterization of TMSE reactions is difficult to achieve in cells, particularly for genetically encoded systems. Cotranscriptionally encoded RNA strand displacement (ctRSD) systems, which operate analogously to TMSE circuits but can be genetically encoded for *in situ* circuit production, hold promise for expanding the application space of TMSE circuits. ctRSD circuits have been extensively characterized in *in vitro* transcription (IVT) environments^24–26^ and were recently deployed in bacteria using a toehold exchange (THE) riboregulator that couples RNA strand exchange to fluorescent protein expression^27^ (Figure 1B). However, ctRSD circuits in bacteria exhibited some behaviors that deviated from the expectations from IVT studies, raising questions about the role of the cellular environment in TMSE and the extent to which design principles elucidated in IVT translate to cells.

Cell-free expression systems^28,29^ (CFES) can bridge the gap between IVT systems and cells because they combine features of both environments (Figure 1C). Like simple, buffered *in vitro* systems, CFES are easy to operate and customize, enabling direct manipulation of reaction composition. For example, users can add any number of DNA templates encoding genetic circuit components, each at a defined concentration, a flexibility infeasible in cellular systems. Further, because they are non-living, non-replicating systems, CFES support kinetic measurements with tools unsuitable for cells^3,30^. In contrast to simple *in vitro* environments used to characterize TMSE reactions, such as buffered salt solutions^15^ or transcription reactions^24^, CFES can execute both transcription and translation. Lysate-based CFES can closely mimic the cellular environment, containing residual nucleic acids, small molecules, and proteins native to the host organism. Together, these characteristics make CFES an appealing platform for rapid prototyping of genetic circuits, both as a testbed for circuits designed for implementation in cells and as the main expression platform^31–34^. In addition, several studies have leveraged CFES to prototype RNA circuits and identified potential characteristics of CFES confounding prototyping efforts and responsible for diverging behaviors across CFES and cellular systems^32,33,35^.

Here, we use CFES to mechanistically characterize the properties of genetically encoded RNA strand exchange circuits that regulate protein expression. By combining measurement tools previously used in either IVT systems or cells — but incompatible with both systems —, we simultaneously measure transcription, RNA strand exchange, and translation dynamics of more than 30 different circuit designs using sequences from the ctRSD toolkit^24,25^. Using these measurements, we unambiguously characterize how changes in input and gate design influence strand exchange and protein expression. We conduct our measurements in *E. coli-*based CFES in both purified protein and lysate formats, which allows us to elucidate phenomena that appear to be related to cellular environment rather than component design. Additionally, we leverage the open environment of CFES to change their composition and identify a potential interaction between non-coding RNAs and cellular components that interferes with strand exchange and informs circuit design. Leveraging the unique properties of TMSE and the mechanistic insights we identified, we demonstrate how THE riboregulators can be applied for bidirectional gene regulation, multi-input logic, and small molecule biosensing, achieving with a single device the different capabilities that would require five different designs with existing nucleic acid biotechnologies. By bridging the gap between *in vitro*^24^ and cellular environments^27^, these results lay the foundation for new applications of TMSE and for future studies exploring interactions between cellular components and genetic circuits.

## RESULTS AND DISCUSSION

### Study and measurement overview

Cotranscriptionally encoded RNA strand displacement (ctRSD) circuits are composed of single-stranded input RNAs and partially double-stranded gate RNAs that are transcribed from DNA templates (Figure 1B). Gates are initially transcribed as a single RNA strand that folds into a hairpin. An internal self-cleaving ribozyme induces the hairpin transcript to cut, producing a partially doubled-stranded gate suitable for TMSE. After transcription, folding, and ribozyme cleavage, a gate presents a single-stranded input toehold (*u*′) that can recruit a complementary input RNA to bind and initiate strand exchange. Successful strand exchange releases a single-stranded output strand from the gate, which can initiate other strand exchange reactions. A toehold exchange (THE) riboregulator represents a slight modification of a ctRSD gate that couples TMSE to protein expression^27^.

The modularity and composability of TMSE gates is enabled by a process of toehold exchange^15^, whereby binding of the input toehold on the gate is exchanged for dissociation of the output toehold on the gate (Figure 1A). The desired measurement of a TMSE reaction is the rate of, or extent of, output toehold dissociation from a gate when different input sequences or concentrations are supplied. In *in vitro* environments, such as buffered solutions or transcription reactions^15,24^, a fluorescently labeled molecular beacon (MB) complex is added to measure TMSE; the dissociation of the output toehold from a gate initiates the displacement of a fluorescent strand on the MB (Figure 1B). In bacteria, THE riboregulators enable measurement of TMSE reactions by coupling dissociation of the output toehold to expression of a fluorescent protein^27^; in this scheme, the output toehold of a gate is designed to be a sequestered ribosome binding sequence (*rbs*) such that TMSE initiates translation (Figure 1B). However, it remains untested how well the MB and fluorescent protein measurements correlate for elucidating differences in TMSE reactions.

To resolve this measurement gap, CFES allows us to conduct both MB and fluorescent measurements simultaneously on the same system. We can easily add the MB to the reactions, which also execute transcription and translation necessary for measuring TMSE with THE riboregulators. Additionally, we can include a fluorogenic Pepper RNA aptamer construct downstream of the fluorescent protein coding sequence on THE riboregulators^36–38^ to explore the role of transcription kinetics across designs and environments.

Here, we characterized THE riboregulators with the above measurements in both a reconstituted system (PURExpress) and a lysate-based system (extract from BL21 Star (DE3) *E. coli*) (Figure 1C). PURExpress, subsequently termed PURE, is a commercially available mixture of purified proteins that enable transcription and translation, and provides an accessible, well-defined platform for rapid genetic circuit prototyping^39,40^. PURE has minimal nuclease activity, which limits the role of RNA degradation and enables use of linear DNA templates for expression. Lysates from BL21 Star (DE3), a common host strain for protein expression^41^ and CFE^42^, retain nucleases and other cellular components after cell lysis, and thus presumably resemble the cellular environment more closely. This strain was also used to test THE riboregulators in *E. coli*^27^. We selected plasmids rather than linear DNA to enable comparison between the two CFES and between CFES and cells^27^, with input and gate sequences encoded on separate plasmids for independent control of concentrations. We encoded RNA strand exchange components on high-copy plasmid DNA under constitutive expression from a T7 RNAP promoter, and optimized reaction conditions for our measurements in both CFES (Supplementary Section 2).

### Validating individual measurements in CFES

Before measuring TMSE, we first conducted control experiments to ensure that each of our intended measurements worked independently in both CFES. To validate MB measurements, we titrated a plasmid expressing the output strand of a THE riboregulator, which should react directly with the MB (Figure 2A). In these and subsequent control experiments, we kept the total DNA concentration constant at 20 nmol/L across samples by adding a template expressing a non-complementary input RNA (Io). A constant transcription load allows for comparison across samples as the overall transcription rate depends on the total DNA template concentration (Supplementary Figures 8-10). As expected, the MB signal increased faster at increasing output plasmid concentrations in both CFES (Figure 2B). In lysate, the rate of MB signal increase was over 2-fold slower than in PURE and output plasmid concentrations below 5 nmol/L were unable to saturate the MB, which could be due to degradation of the output RNA.

**FIGURE 2:**
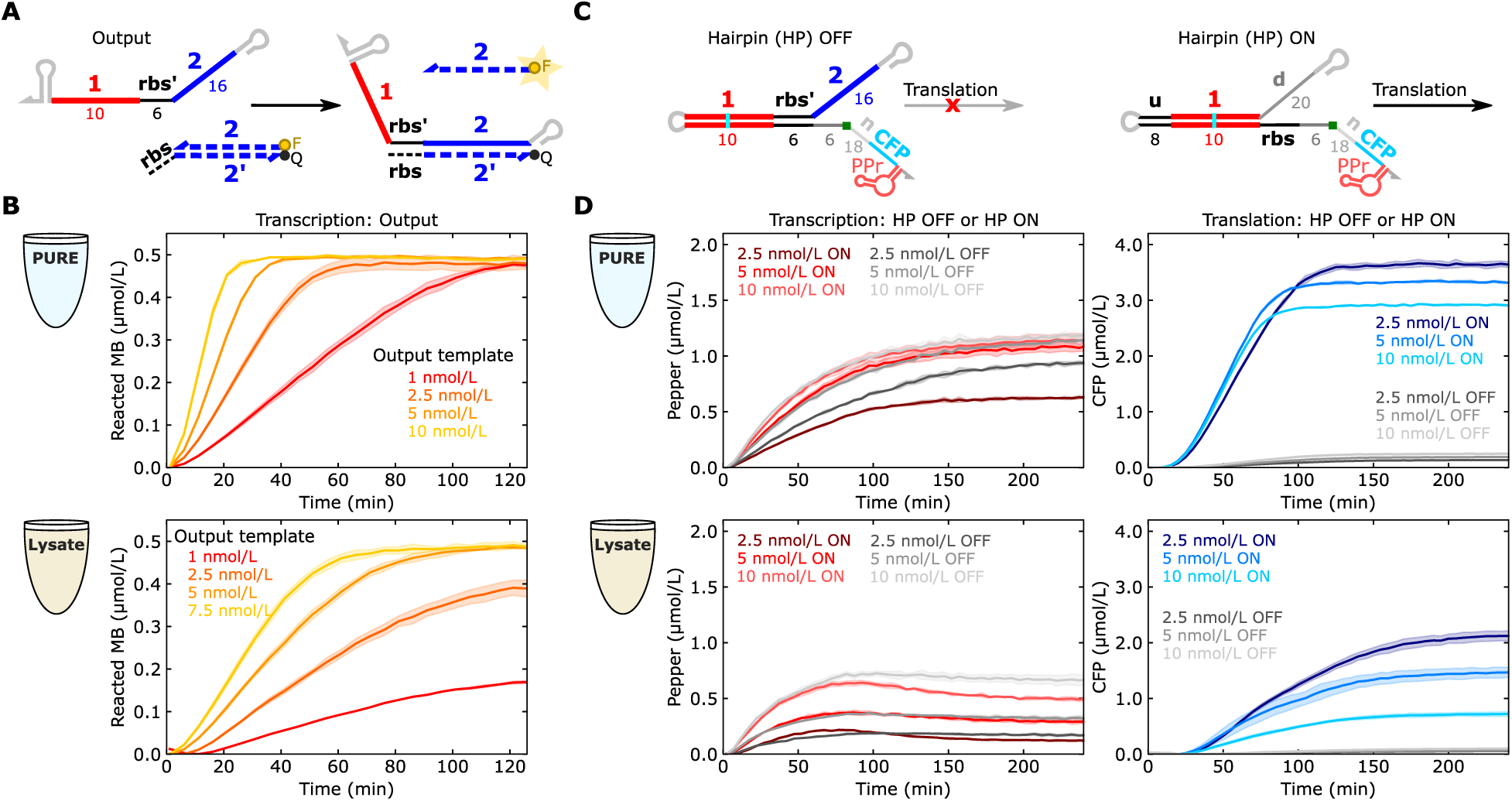
Characterizing measurements with DNA templates expressing control constructs. (**A**) Schematic of a control for molecular beacon (MB) measurements in CFE systems. An output RNA that reacts with the molecular beacon was constitutively transcribed alongside 500 nmol/L of molecular beacon complex. (**B**) Reacted molecular beacon kinetics for varying concentrations of output template DNA in PURE (top) or lysate (bottom). (**C**) Schematics of translation controls for fluorogenic aptamer and fluorescent protein measurements. HP OFF and HP ON were transcribed separately to determine minimum and maximum translation levels for a THE riboregulator. Domain lengths (non-bold numbers below schematics) were chosen to be consistent with previous ctRSD gate designs^24,25^, while also maximizing the predicted translation rate for the ON state^27^. (**D**) Pepper and CFP signals at different template concentrations of HP OFF and HP ON in PURE (top) or lysate (bottom). In (C) and (E), the total DNA template concentration was 20 nmol/L, with Io making up the difference from the concentrations shown in the plots, and the shaded regions indicate the standard deviation of three technical replicates. Additional experiments at lower total template concentrations are shown in Supplementary Figures 9 and 10.

We next tested the Pepper RNA aptamer and cyan fluorescent protein (CFP) measurements using plasmids encoding hairpin (HP) equivalents of an OFF and ON THE riboregulator (Figure 2C). These constructs serve as best case scenarios for OFF and ON protein expression for a THE riboregulator as HP OFF strongly sequesters the *rbs* and HP ON translates directly after folding. As expected, CFP production was low for HP OFF plasmids and high for HP ON plasmids in both CFES (Figure 2D). Interestingly, the final protein concentration was inversely proportional to HP ON plasmid concentration, perhaps due to faster depletion of energy as the rate of mRNA production increases. In both CFES, the Pepper signal increased with increasing plasmid concentration for both HP OFF and HP ON, although in PURE the differences between the 5 nmol/L and 10 nmol/L HP plasmid concentrations were minimal. In lysate, the Pepper signal decreased steadily after reaching a maximum value, reflecting ribonuclease-mediated degradation of transcripts. In these control experiments, both MB and Pepper are indicators of transcription rates, but Pepper took ∼2 times longer than MB to reach 0.5 µmol/L. This lag could be due to Pepper’s location at the end of the transcript, slow aptamer folding, or slow dye binding (Figure 2B vs. 2D).

### Tuning protein expression by modulating transcription or translation strength

Having validated each individual measurement in both CFES, we next applied these measurements to characterize the full RNA strand exchange process. Specifically, we tested different equimolar concentrations of input and gate templates (Figure 3A) using either a complementary input (I{u1d}, ON state) or a non-complementary input (Io, OFF state). As expected, we observed that increasing concentrations of input and gate templates increased the rate of production across all measurements (Figure 3B). In PURE, MB and CFP measurements showed similar kinetics and CFP levels followed a similar trend to the HP ON control, with increasing gate concentrations resulting in faster CFP production but lower final CFP concentration. However, in lysate the MB signal was indistinguishable between ON and OFF states, and the CFP trends contrasted the HP ON control trends, with increasing gate concentrations resulting in higher CFP production levels. These results suggest that RNA strand exchange may be inefficient in lysate, resulting in a low concentration of gate in the ON state and thus weak protein production. Notably, CFP production in lysate was greater for a THE riboregulator in the OFF state than HP OFF, which could indicate incorrect folding of the gate that increases leak and inhibits RNA strand exchange. We did not measure ribozyme cleavage efficiency directly, but gates with a mutated ribozyme incapable of cleavage produced low protein concentration in the ON and OFF states in both CFES (Supplementary Figure 11). Additionally, we observed similar performance in both CFES for THE riboregulators with an alternative input toehold sequence (*v*), an alternative ribozyme sequence (*Rh*), and an alternative input branch migration sequence (domain *4*) (Supplementary Figure 12). Together, these results demonstrate tuning protein expression levels with RNA strand exchange by modulating the transcription rates of components (Figure 3B, Supplementary Figure 13C).

**FIGURE 3:**
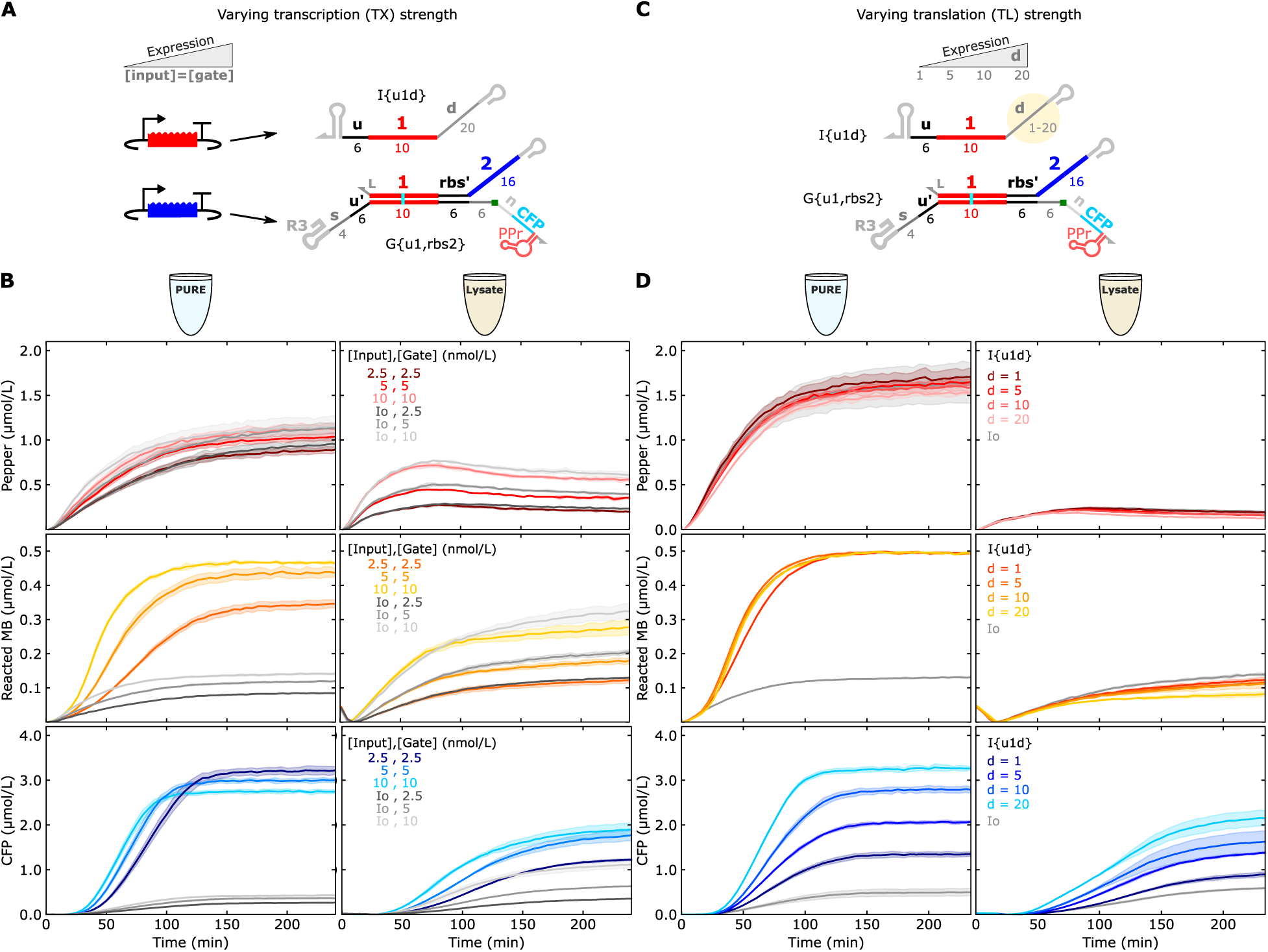
Tuning protein expression by modulating transcription or translation strength. (**A**) Transcription strength was modulated by changing the concentrations of input and gate templates. (**B**) Kinetics of the system in (A) in PURE (left) and lysate (right). Input and gate DNA template concentrations are labeled in each plot with Io added to keep a constant 20 nmol/L total DNA template across samples. Additional conditions with unequal template concentrations are shown in Supplementary Figure 13. (**C**) Translation strength was varied by changing the length of the single stranded *d* domain on the input. This domain serves as an unstructured ribosome standby site that enhances translation initiation^46^, allowing translation strength to be controlled without changing the *rbs* sequence. (**D**) Kinetics of the system in (C) in PURE (left) and lysate (right). In PURE, the input and riboregulator template concentrations were both 5 nmol/L. In lysate, the input template concentration was 5 nmol/L and the riboregulator template concentration was 2.5 nmol/L. In (B) and (D), the shaded regions indicate the standard deviation of three technical replicates.

We next explored tuning protein expression by modulating the translation strength of a THE riboregulator at a constant transcription level. Translation strength is typically tuned by changing the complementarity of the *rbs* to *E. coli* 16S rRNA^43^. However, changing the sequence of an mRNA has disadvantages, as minor changes can alter RNA folding^44^ to influence protein expression in difficult-to-predict ways. THE riboregulators offer a method to modulate translation strength without changing the sequence of the protein-encoding mRNA by instead changing the length of a single-stranded domain, termed *d,* upstream of the branch migration sequence in the input (Figure 3C). The *d* domain is hypothesized to serve as a standby site for ribosomes, thereby influencing ribosome binding and translation initiation^45–47^. In both PURE and lysate, decreasing the *d* domain length from 20 bases to 1 base decreased the rate of protein production (Figure 3D), consistent with results in *E. coli*^27^. Pepper measurements indicated global transcription rates were similar across samples and MB measurements in PURE indicated RNA strand exchange was unchanged for inputs with different *d* domain lengths (Figure 3D), suggesting protein expression differences were primarily due to changes in translation strength of the ON gate.

### Tuning protein expression by modulating RNA strand exchange rates

TMSE offers kinetic control of reactions by altering the lengths of input and output toeholds. This capability is crucial for engineering circuits that execute logic, subtraction, and timing^16,17,48,49^. TMSE can be viewed as a three-step process whereby an input strand can reversibly bind to a gate and undergo a random walk of branch migration. Once the input strand nears the end of the branch migration domain, the output strand can reversibly unbind from the gate^23^ (Figure 4A). By modulating the relative lengths of the input and output toeholds of a strand exchange reaction, the rate of output production can be controlled^15,23^.

**FIGURE 4:**
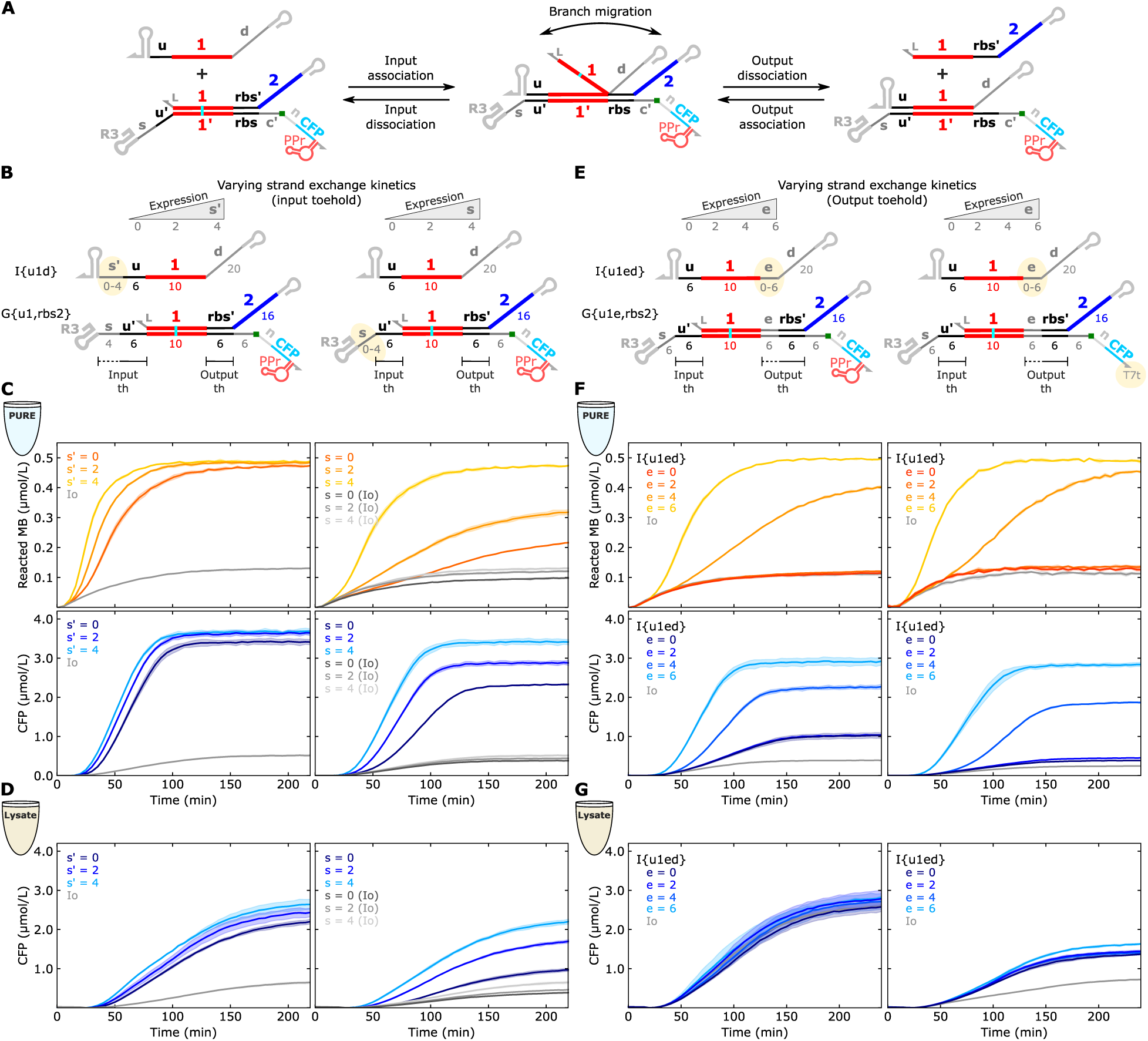
Tuning protein expression by modulating RNA strand exchange parameters. (**A**) An expanded schematic of toehold mediated strand exchange (TMSE). With respect to base pairing thermodynamics, TMSE is a reversible process composed of three steps: input/output association, branch migration, and input/output dissociation^23^. In the canonical THE riboregulator design, the *u* and *rbs* toeholds are both six bases so TMSE is not heavily biased in either the forward or reverse direction. (**B**) TMSE can be biased in a forward direction by increasing the length of the toehold on the input strand to disfavor input dissociation (left) or by changing the length of a single-stranded spacer (*s*) between the input toehold and the ribozyme on the THE riboregulator (right). Longer spacers disfavor input dissociation. (**C**,**D**) Kinetics of the systems in (B) in PURE (C) or lysate (D). Pepper measurements and results for additional combinations of *s:s′* lengths are in Supplementary Figure 14. (**E**) TMSE can be biased in the reverse direction by increasing the length of the output toehold that has to dissociate after branch migration, designated by *e* on the riboregulator (left). Longer output toeholds can be partially overcome by increasing the length of the input toehold (right). (**F**,**G**) Kinetics of the systems in (E) in PURE (F) or lysate (G). In PURE, the input and riboregulator template concentrations were both 5 nmol/L. In lysate, the input template concentration was 5 nmol/L and the riboregulator template concentration was 2.5 nmol/L. Pepper measurements and results for a gate with a 4-base *e* domain are in Supplementary Figure 15. In (C), (D), (F) and (G), the shaded regions indicate the standard deviation of three technical replicates.

We reasoned these properties of TMSE could be leveraged to tune protein expression with THE riboregulators. We first increased the degree of complementarity (*via* the *s′* domain) between the input strand toehold and the gate, which should decrease the input strand dissociation rate and result in faster strand exchange (Figure 4B, left). In PURE, output RNA production, as measured by the MB, increased with increasing input toehold complementarity (Figure 4C, left). As expected, the rates of protein production increased with increasing input toehold complementarity in both CFES (Figure 4C,D, left). In addition to changing the length of complementarity with the input toehold, the length of the spacer (*s*) domain on the gates should modulate strand exchange kinetics (Figure 4B, right). In IVT experiments, decreasing the length of the spacer from 4 bases to 0 bases decreased the strand exchange rate constant by nearly two orders of magnitude^24,25^, likely due to steric hindrance^50,51^. Decreasing spacer length on a THE riboregulator greatly reduced the rate of protein production in both CFES (Figure 4C,D, right).

We next explored how increasing the length of the output toehold influenced strand exchange and protein expression kinetics. We designed a gate with a double-stranded branch migration domain extended by 6 base pairs (*e:e′* domains). If used with an input strand without an *e* domain, a total of 12 bases (*e*+*rbs′*) would have to dissociate after branch migration for the output to be released and protein translation to initiate. By designing input strands with increasing *e* lengths, the length of the gate output toehold can be modulated between 6 bases and 12 bases (Figure 4E, left).

Modulating the length of the output toehold had different effects on protein expression kinetics in PURE and lysate. In PURE, a gate with a 12-base output toehold showed minimal output release and protein expression, and decreasing the length of the output toehold increased the rates of strand exchange and protein expression (Figure 4F, left). In lysate, however, the extended gate had high protein expression even with a non-complementary input (Io), which masked any trend in protein expression with increasing output toehold length (Figure 4G, left). This high level of leak was not observed in *E. coli*^27^, which used an extended gate with a different 3′ untranslated region (UTR) that lacked the Pepper aptamer and used a less efficient terminator^52^. Testing the extended gate with this alternative 3′ UTR (Figure 4F, right) reduced leak in lysate, but, in contrast to PURE, protein expression in the ON state did not vary with output toehold length (Figure 4G, right). This phenomenon was not specific to this gate sequence as variants with a shorter *e* domain, an alternative *e* sequence, or an alternative input domain had the same trend (Supplementary Figures 15 and 16). With the alternative 3′ UTR, this unexpected trend now matched the trend observed for THE riboregulators in *E. coli*^27^, suggesting cellular components not involved in transcription or translation may play a role.

### Effects of cellular components on RNA strand exchange cascades

To further investigate how the biological context affects RNA circuit behavior, we explored a previous finding that multi-layer strand exchange cascades required that gates upstream of the THE riboregulator include a hairpin structure on the 3′ end of their output strand to transduce a measurable signal in *E. coli*^27^ but not in IVT^24,25^. We hypothesized that, in *E. coli*, the 3′ hairpins conferred stability to the output strands of each layer of the cascade, thereby preventing ribonuclease-mediated degradation before the strands were able to react downstream. If this were the case, we would expect PURE – which manufacturers state to have minimal ribonuclease activity – to behave like IVT. Surprisingly, multi-layer cascades in PURE required upstream gates with a 3′ hairpin on their output strands to produce signal above the OFF state (Supplementary Figure 17). We confirmed this phenomenon was related to the free output strand rather than the dsRNA gate, and that it was not sequence specific, as the same trend held for upstream gates with alternative input domains, ribozymes, output domains, and 3′ hairpins (Supplementary Figures 18 and 19). This phenomenon likely affects all input RNAs and was previously unidentified because all input RNAs used in this study had terminators with 3′ hairpins by design. In fact, multi-layer strand exchange cascades composed of gates with an inverted transcription order^25^ – which results in a terminator hairpin at the 3’ end of the output strands – worked well in PURE (Supplementary Figure 20).

We hypothesized that ribosomes – which can bind to unstructured regions of RNA promiscuously^47,53,54^ – could be interacting with ssRNA components and interfering with strand exchange, with the structure of a 3′ hairpin disfavoring such interactions. To explore this possibility, we used a THE riboregulator triggered by an output strand (O{5,u1d}) with and without a 3′ hairpin in a PURExpress reaction lacking ribosomes (Δribosomes kit) (Figure 5A). In the absence of ribosomes, strand exchange proceeded similarly with and without a 3′ hairpin on O{5,u1d} (Figure 5B, left). Using a ribosome solution supplied by the manufacturer, we added ribosomes back to the Δribosomes kit to a concentration equivalent to previous PURE experiments (Supplementary Figure 21). In this reaction, the MB and CFP signals in the reaction using O{5,u1d} without a 3′ hairpin were substantially reduced (Figure 5B, middle). Similar results were obtained for a 2-layer cascade (Supplementary Figure 22). We confirmed these results were not due to differences in ionic conditions in the reactions with and without ribosomes (Supplementary Figure 23). Suspecting a specific aspect of ribosomal activity might be at play, we included antibiotics that inhibit translation initiation, elongation, or termination^55^ in our reactions, but saw no recovery of activity for O{5,u1d} without a 3′ hairpin (Supplementary Figures 24-25). These results suggest that, while something in the ribosome solution underlies the observed hairpin dependence, ribosomal reactions associated with active translation do not interfere with RNA strand exchange.

**FIGURE 5:**
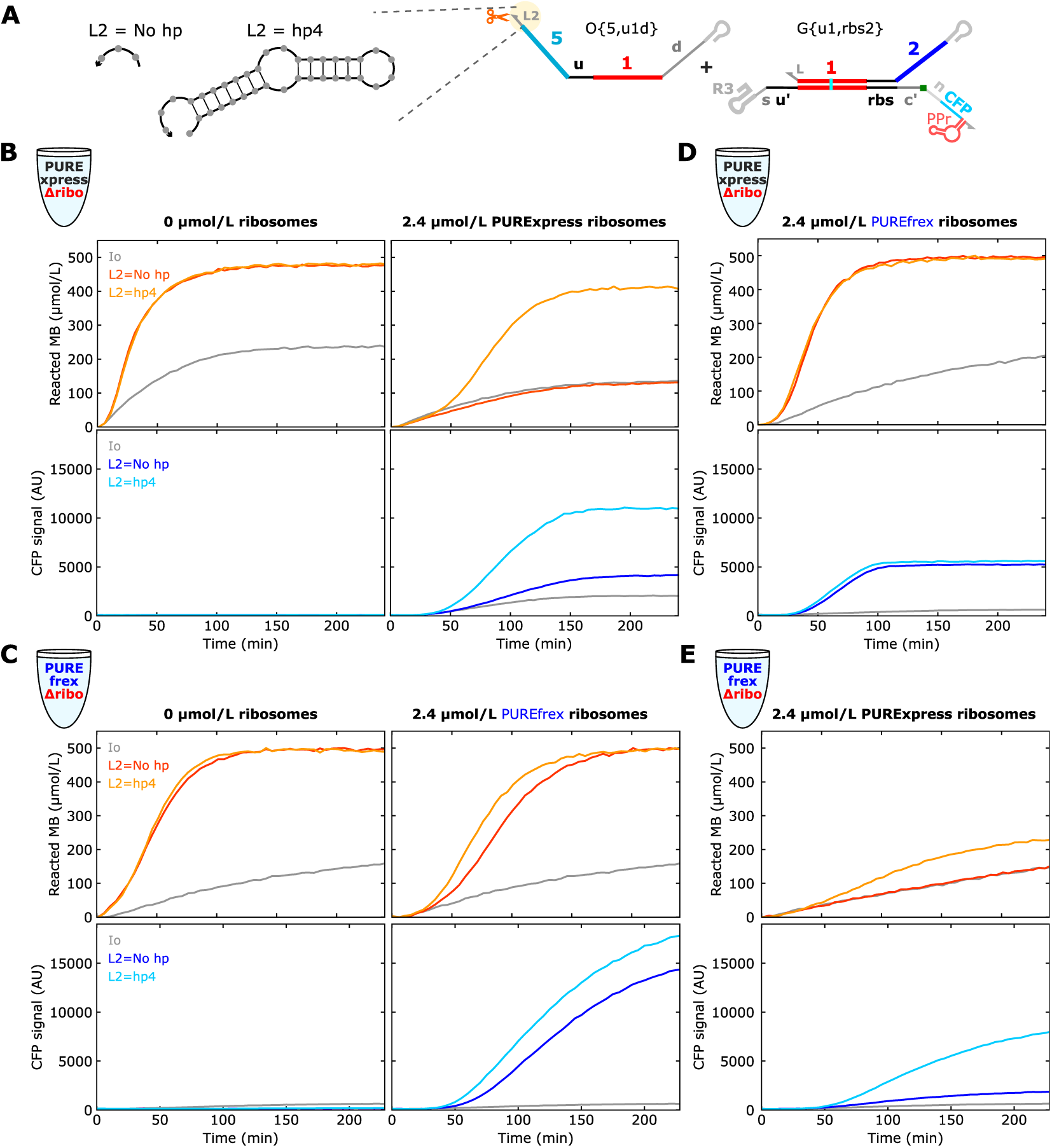
Characterizing different ribosome solutions with RNA strand exchange components. (**A**) Schematic of O{5,u1d}, which mimics an upstream output of a multi-layer cascade, as the input to a THE riboregulator. O{5,u1d} without (L = No hp) and with a 3ʹ hairpin (L = hp4) were tested. Kinetics of the system in (A) in **(B)** PURExpress and **(C)** PUREfrex without (left) and with (right) ribosomes, in (**D**) PURExpress with PUREfrex ribosomes, and in (**E**) PUREfrex with PURExpress ribosomes. Output and gate DNA template concentrations were each 4 nmol/L with Io added to keep a constant 20 nmol/L total DNA template across samples. All reactions had a single replicate. Results with Pepper aptamer kinetics are shown in Supplementary Figure 29.

To determine whether our results were unique to the PURExpress kit we were using, we tested another lot of PURExpress and a PURE system from a different manufacturer, called PUREfrex. Both lots of PURExpress exhibited the hairpin dependence, but to differing degrees (Supplementary Figure 26). In PUREfrex with ribosomes, removing the 3′ hairpin on O{5,u1d} only slightly decreased MB and CFP signals (Figure 5C). Further, adding PUREfrex ribosomes to a PURExpress reaction led to identical MB and CFP signals for O{5,u1d} with and without a 3′ hairpin (Figure 5D), while adding PURExpress ribosomes to a PUREfrex reaction gave rise to the hairpin dependence (Figure 5E). Similarly, addition of PURExpress ribosomes to an IVT reaction exacerbated the 3′ hairpin dependence, while addition of PUREfrex ribosomes did not (Supplementary Figure 27). Together, these results suggest that the requirement for a 3′ hairpin is likely unrelated to an interaction between ribosomes and RNA strand exchange components, but may be related to impurities in the ribosome solution that arise during the ribosome purification process, such as 3′-specific RNA exonucleases (Supplementary Figure 28).

### Using mechanistic insights to realize THE riboregulator applications

The unique properties of TMSE — reversibility, composability and kinetic tunability — should make THE riboregulators a versatile platform for many applications, such as precise control of RNA and protein expression, biosensing, and programmable molecular computing. To showcase this versatility, we developed representative applications using the same general THE riboregulator design for many different functions. We drew on the mechanistic insights from Figures 2-5, in particular the observation that RNA input and output strands need a 3′ hairpin to work in PURE.

Based on our experiments with the extended branch migration domain (Figure 4F), we expected the THE riboregulator reaction to be reversible, allowing a single design to both activate or repress protein expression depending on whether the RNA input or RNA output is produced (Figure 6A). Bidirectional gene regulation is useful for feedback loops and dual-mode biosensing, but typically involves separate modules for activation and repression^8,13^. We demonstrated this bidirectional gene regulation in two different kinetic regimes by modulating the length of the spacer (*s*) domain on the gate (Figure 6B). Importantly, we designed both input and output strands with a 3′ hairpin terminator rather than using commonly employed runoff transcription; inputs and outputs without a 3′ hairpin had minimal activity in both regulation directions (Supplementary Figures 30, 31).

**FIGURE 6:**
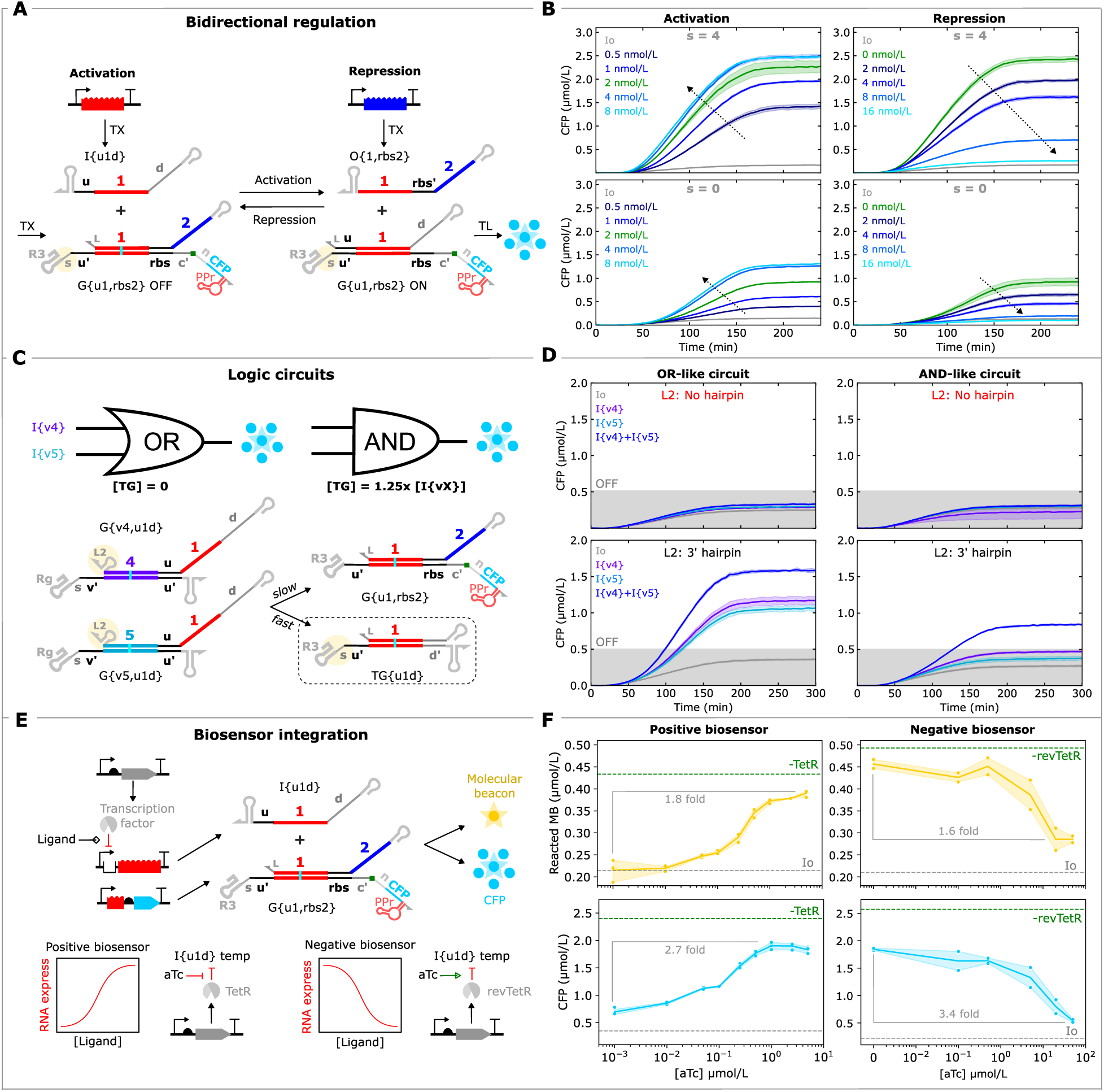
Versatile applications of THE riboregulators in PURExpress. **(A)** Schematic of bidirectional regulation of gene expression by THE riboregulators. **(B)** Activation or repression of CFP expression by modulating input (left) or output (right) template concentrations, respectively. **(C)** Thresholding-based OR-like (left) and AND-like (right) logic elements. Gates G{v4,u1d} and G{v5,u1d} interact with the two inputs, I{v4} and I{v5}, producing outputs that trigger G{u1,rbs2}. The AND element also includes a threshold gate (TG) that reacts faster than G{u1,rbs2} with either of the outputs from the upstream layer (due to TG having a 4-base *s* domain that the riboregulator lacks), such that CFP is produced only when both inputs are present and exceed the concentration of the threshold. **(D)** CFP production from the logic elements from inputs without (top) and with (bottom) a 3′ hairpin. CFP signal only exceeds a threshold concentration of 5 µmol/L when the inputs have a hairpin. MB and Pepper aptamer measurements for this circuit are in Supplementary Figure 32. **(E)** Integration of THE riboregulators with anhydrotetracycline (aTc) biosensors. **(F)** MB and CFP signals at 4 h for different concentrations of aTc for both biosensors. The Methods section (“THE riboregulator application experiments”) describes the experiments in detail. In (B), (D), and (F, positive biosensor), the shaded regions indicate the standard deviation of three technical replicates. In (F, negative biosensor), the shaded regions indicate the standard deviation of two technical replicates.

THE riboregulators can also be combined with other RNA circuit components to enable more complex information processing circuits. As a demonstration, we implemented a threshold-based method for executing multi-input OR and AND logic, a common element in many state-of-the-art TMSE circuits^17^. The circuit employs two ctRSD gates that, upon strand exchange, release RNA outputs complementary to a THE riboregulator (Figure 6C), thus enabling OR-like logic. To achieve AND-like logic, the circuit also includes a threshold gate designed to react with the O{X,u1d} strands much faster than the THE riboregulator. If the concentration of the threshold gate template is set slightly above the concentration of either of the input templates, both inputs will be required to produce an output. Our kinetic results from Figure 4 suggested that designing a threshold gate with a 4-base spacer domain and a THE riboregulator without a spacer domain should produce the necessary differences in reaction rates to achieve thresholding. As this circuit is a 2-layer cascade, we expected the ctRSD gates to need a 3′ hairpin on their output strands for the circuit to work. As expected, neither scheme transduced CFP signal above the OFF state when 3′ hairpins were left off the ctRSD gates, while the circuit with these 3′ hairpins exhibited the designed logic (Figure 6D). While there are many ways to build RNA logic circuits, this implementation highlights the composability of TMSE reactions, as the same circuit design is used for both logic elements. In contrast, other RNA biotechnologies typically require a different gate design for each logic operation, often imposing different constraints on the input sequences^9,12^.

An obvious application in which logic circuits could be important is “smart” biosensing. Thus, as our last demonstration we coupled THE riboregulators with transcription factor-based biosensors. IVT biosensors that use purified transcription factors that respond to small molecules have been developed and integrated with DNA-based TMSE information processing layers^3,56^. Inspired by this scheme, we integrated these biosensors with a THE riboregulator in PURE with three modifications: (1) we programmed the transcription factors to produce an RNA input to a THE riboregulator, (2) we added a 3ʹ hairpin terminator to each DNA template — instead of using runoff transcription templates — to mitigate potential issues with the ribosome solution, and (3) we coexpressed the transcription factors in the PURE reaction instead of purifying them (Figure 6E). This integrated system offers a few potential benefits over previously described IVT systems^3,56^. For example, the incorporation of translation provides a layer of signal amplification over the MB reporter, which could increase sensitivity. Additionally, *in situ* production of both circuit components and transcription factors enables fast prototyping and reduced labor and cost, particularly with respect to the TMSE components.

Using template concentrations based on previous transcription titrations (Figure 2), we tested two different biosensors for anhydrotetracycline (aTc), one that is induced by aTc to derepress transcription (positive biosensor, TetR) and one that is induced by aTc to repress transcription (negative biosensor, revTetR^57^). Both biosensors showed the expected response to titrations of aTc (Figure 6F). Further, we observed a higher fold change for the CFP signal than the MB signal for both biosensors, indicating that incorporation of translation provided signal amplification. This signal could potentially be amplified further by replacing a fluorescent protein output with an enzyme, such as LacZ^58^.

Together, these results highlight the versatility of THE riboregulators to interface with a wide range of cell-free biotechnologies and how the mechanistic insights we uncovered are crucial for successful implementation.

## CONCLUSIONS

Here we leveraged CFES to characterize the mechanisms underlying genetically encoded RNA strand exchange circuits. Our characterization was enabled by simultaneous measurements of transcription, translation, and strand exchange. Using these measurements, we demonstrated how different properties of TMSE reactions can be modulated to tune protein expression at the transcriptional (Figure 3B), post-transcriptional (Figure 4), and translational levels (Figure 3D), illustrating the versatility of THE riboregulators for programming biology. In principle, expression from a single THE riboregulator design could be tuned at each of these levels by changing only the sequence of a small non-coding RNA input (Supplementary Figure 35), a level of fine-tuned control not readily available with existing riboregulator technologies. The ability to manipulate the composition of CFES allowed us to identify potential interactions between RNA strand exchange components and species present in ribosome solutions, bridging our gap in understanding between previous IVT studies and recent studies in *E. coli*^27^. We expect the framework for prototyping in CFES presented here will be applicable for characterizing, understanding, and successfully deploying biotechnologies beyond the specific RNA circuits we studied^2,7–9^.

A central motivation of this study was to use CFES to complement and supplement previous TMSE characterization in IVT and in cells. Reassuringly, we found the same toehold, input/output, and ribozyme sequences generally functioned well across environments, highlighting interoperability. However, different CFES may serve different purposes when working with RNA circuits. For evaluating initial designs and comparing to IVT, a well-defined system like PURE is ideal. This system minimizes confounding variables such as RNA degradation, simplifying the interpretation of results. However, many trends that agreed in IVT and PURE did not transfer to lysate and cells^27^ (Figure 4). Thus, if the primary research question is to explore phenomena observed in cells, then characterization in lysate produced more comparable results. Notably, we found that behavior in lysate, like in cells, was more sensitive to changes in 3′ UTRs than in PURE (Supplementary Section 5). In lysate, however, we observed very different trends in protein expression when the 3′ UTR of the THE riboregulator was changed (Figure 4, Supplementary Figure 16) compared to the trends observed in *E. coli*^27^. Therefore, it is important to keep in mind that adding fluorescent aptamer sequences or changing terminator sequences may change results substantially.

CFES allowed us to study three phenomena that had markedly different results in IVT compared to *E. coli*: the trends of increasing (1) input and (2) output toehold lengths on strand exchange and (3) the need for a 3′ hairpin on input/output strands. The trend of increasing strand exchange rate when increasing the input toehold length from 6 bases to 10 bases held across IVT^24^, PURE, and lysate, suggesting the opposite trend observed in *E. coli*^27^ is unique to the cellular environment. However, we can tune reaction kinetics across all environments by increasing the length of the *s* domain on the gate, which had the same effect in IVT, PURE, lysate, and *E. coli*. The relationship between output toehold length and strand exchange was split between reconstituted systems (IVT and PURE) and cell-like systems (lysate and cells). The IVT and PURE results matched expectations^15^ from a base-pairing perspective, with increasing output toehold length showing decreasing strand exchange (Figure 4). In contrast, lysate and cells showed little dependence on output toehold length, which could implicate RNA degradation pathways^59–61^. The requirement of 3′ hairpins on RNA inputs for strand exchange, observed in *E. coli*^27^ and in PURE, might also be related to RNA stability (Figure 5). Different PURE lots and kits showed varying levels of hairpin dependence, likely due to differences in the purity of ribosome preparations. Nonetheless, hairpins enabled effective strand exchange in all environments and was necessary for realizing logic and biosensing circuits (Figure 6), illustrating the importance of this design feature.

Our results only begin to characterize the myriad possible interactions with cellular components and synthetic RNA systems. A class of enzymes to explore further are RNases^59–62^, which could play a role in some of the differences between PURE and lysate. Endogenous RNAs absent in reconstituted systems or at low concentrations in lysate systems could also interfere with RNA strand exchange^63,64^. Additional experiments should elucidate the effect of ribonucleases on RNA strand exchange, even at the low concentrations that seem to be present in PURExpress. Beyond ribonucleases, some RNA-binding proteins may enhance RNA:RNA interactions. Many of these proteins have low sequence specificity but preferentially interact with 3′ secondary structures^62,65,66^. Studying the above interactions will likely require new measurements or improvement of the measurements presented here. For example, the molecular beacon measurement was not sensitive enough to characterize strand exchange in lysate (Figure 3), preventing us from assessing strand exchange without translation. Other molecular beacon sequences or designs could resolve this issue, but we found molecular beacons with different toeholds had markedly different ON/OFF ratios across IVT, PURE, and lysate (Supplementary Section 6). Therefore, other methods to measure strand exchange more directly may be necessary. That being said, our assay using RNA inputs with and without 3ʹ hairpins serves as a sensitive method for assessing impurities across different batches of PURE, although more work needs to be done to identify the specific species behind this phenomenon.

Despite the differences observed across environments and other yet untested interactions, the work described here lays the foundation for programming biology with RNA circuits across environments and applications, with THE riboregulators showing promise for broad and versatile implementation. Specifically, our results showcase the ease of implementing THE riboregulators for processing logic and biosensing. Beyond small molecule biosensors, THE riboregulators could provide the basis for cell-free biosensors that detect small RNAs^67^, such as miRNAs, because THE riboregulators require only short (16 base) inputs. Given that TMSE reactions are reversible, THE riboregulators are also well suited for discriminating single-nucleotide polymorphisms^6,22^. Further, THE riboregulators could be connected to protein biosensors that regulate transcription^68^. Applications involving the expression of multiple proteins could benefit from cascades of RNA strand exchange reactions. With cascades, users could control the timing and order of production of different proteins^48^. Further, different *d* domain lengths could be used to independently control translation strength at each layer while maintaining a constant flux of strand exchange. Such autonomous temporal control could be useful for coordinating expression of multi-protein systems^69^ or for operating biosensors that need to coordinate expression of sensing and reporting proteins^70^, especially in closed systems like synthetic cells^71^. Predictive kinetic models relating protein expression levels to RNA strand exchange^25,72–74^ would further facilitate the above applications. Our results identifying a unified set of design principles for using genetically encoded RNA strand exchange circuits across common synthetic biology environments positions this platform to catalyze development of a wide range of new biotechnologies.

## MATERIALS AND METHODS

### DNA, bacterial strains, and chemicals

#### DNA

The majority of the DNA for experiments in the main text was ordered as clonal gene fragments from Twist Biosciences in plasmid backbones provided by the company — either “pTwist Amp High Copy” (high-copy backbone encoding resistance to ampicillin) or “pTwist Kan Medium Copy” (medium-copy backbone encoding resistance to kanamycin). Some DNA sequences encoding inputs and gates were ordered as eBlock gene fragments from Integrated DNA Technologies (IDT), PCR amplified, and subsequently cloned into pTwist backbones or used as linear templates in PURE. See Supplementary Figure 3 for additional details. DNA oligonucleotide primers for PCR were ordered from IDT with standard desalting. DNA sequences, including their origin, are provided in Supplementary File S1. Annotated GenBank files of DNA templates are provided in Supplementary File S2.

#### RNA

The oligonucleotides of the molecular beacon complex were ordered as 2′-O-methyl RNA (2omR) with standard desalting for the fluorophore-modified (HEX) strand and HPLC purification for the quencher-modified (Iowa Black RQ) strand. 2omR bases were used to prevent degradation of the molecular beacon in lysate, but we also found that DNA-based molecular beacons produced spurious fluorescence in PURE (Supplementary Figure 4). Molecular beacon complexes were prepared by mixing 20 µmol/L of each strand in transcription buffer (Thermo Fisher, cat. no. EP0113) and heating to 90 °C for 5 min followed by cooling to 20 °C at a rate of 1 °C/min. Schematics and nomenclature for RNA species are described in Supplementary Figures 1 and 2.

#### Protein sequences

The sCFP3A amino acid sequence, referred to throughout as ‘CFP’ for simplicity, was taken from GenBank (AAZ65848.1). This sequence was then codon optimized for expression in bacteria with IDT’s codon optimization tool.

#### Pepper RNA aptamer

Unless otherwise specified, THE riboregulators used in this work used a dimeric Pepper aptamer grafted onto F30 and tRNA scaffolds^37^. This Pepper aptamer is termed tDF30ppr. A version of this aptamer lacking the tRNA scaffold, termed DF30ppr, was used in control experiments included in the Supplementary Section 5. In both tDF30ppr and DF30ppr, two point mutations were made to the Pepper sequence to reduce sequence complexity and increase the likelihood of successful synthesis.

#### Strains

*E. coli* DH10B (Thermo Fisher, cat. no. EC0113) was used for plasmid cloning and extraction. BL21 Star (DE3) (Thermo Fisher, cat. no. C601003) was used to prepare lysates for cell-free gene expression and to purify proteins.

#### PURE

PURExpress® *In Vitro* Protein Synthesis Kit (cat. no. E6800L), PURExpress® ΔRibosome Kit (cat. no. E3313S), and *E. coli* ribosomes (cat. no. P0763S) were purchased from New England Biolabs. Unless otherwise stated, PURExpress experiments were run using lot #10272724 (lot ‘M’ in Supplementary Figure 26). A different lot (lot ‘C’) was also tested in Supplementary Figure 26. PUREfrex 2.1 (cat no. PF213-2ML-EX) was purchased from GeneFrontier.

#### Chemicals

Supplementary File S4 lists all chemical reagents used in cell-free reactions and all antibiotics used for plasmid selection. Enzyme and kit product numbers are referenced when first introduced below.

### Cloning and PCR

#### PCR

PCRs were conducted with Phusion Flash Master Mix (Thermo Fisher, cat. no. F-548) and 0.5 µmol/L of each DNA primer. Plasmid backbones for cloning were prepared *via* PCR of 0.2 ng/µL plasmid DNA and the following thermocycler protocol: 30 cycles of a denaturing step (30 s, 98 °C), a primer annealing step (30 s, 60 °C), and an extension step (120 s, 72 °C). A 5 min extension at 72 °C was executed at the end of the program. Backbone PCR products were digested by DpnI (New England Biolabs, cat. no. R0176S) to remove methylated plasmid DNA. PCR products were purified using the PureLink PCR purification kit (Invitrogen, cat. no. K310001). To remove non-specific products shorter than 300 bp from plasmid backbone PCRs, a wash step with buffer B3 was included after DNA binding with buffer B2. Purified PCRs were quantified using a Nanodrop spectrophotometer (DeNovix D-11 Series). Linear DNA inserts or transcription templates were prepared similarly except the extension step in the PCR was shortened to 30 s, the DpnI digestion was not included, and the B3 wash step was omitted during purification.

#### Cloning

Plasmids assembled in house were cloned *via* Gibson assembly or inverse PCR. Gibson reactions were prepared using Gibson assembly master mix (New England Biolabs, cat. no. E2611L) with a 3:1 molar ratio of insert to backbone DNA, and incubated at 50 °C for 1 h prior to transformation. For inverse PCRs, DNA primers were first phosphorylated by T4 Polynucleotide Kinase (New England Biolabs, cat. no. M0201S) at 37 °C for 30 min. Inverse PCR products were ligated by T4 DNA Ligase (New England Biolabs, cat. no. M0202S) at room temperature for 2 h.

### DNA sequence verification

Plasmids underwent whole-plasmid or ZeroPrep sequencing by Plasmidsaurus using Oxford Nanopore Technology with custom analysis and annotation.

### DNA plasmid extraction and quantification

Plasmids were transformed into chemically competent DH10B *E. coli*, and transformants were plated on LB agar with appropriate antibiotic and grown overnight at 37 °C. 100 mL of LB with appropriate antibiotic was inoculated with one colony from the transformation and allowed to shake at 37 °C, 250 rpm, for 16 h in a baffled 250 mL flask. After growth, a glycerol stock was prepared using 200 µL of the culture to enable long-term storage at −80 °C and to bypass the transformation step in future DNA extractions.

Cells were harvested by centrifugation at 4 °C, 2700 x g, for 20 min. Plasmid DNA was extracted using the E.Z.N.A Plasmid DNA Midi kit (Omega Biotek, cat. no. D6904-04) and the protocol provided by the manufacturer. Upon being eluted, DNA was precipitated in 100 % isopropanol (Millipore Sigma, cat. no. 34863) and washed in 70 % ethanol (Millipore Sigma, cat. no. E7023) following the kit’s protocol. The DNA pellet was allowed to air dry for 30 min, then resuspended in 30 µL of elution buffer. For ‘High Copy’ plasmids, at this stage the DNA solution typically had a concentration between (2500 and 4000) ng/µL based on spectrophotometric measurements on a Nanodrop instrument.

Each plasmid underwent an additional purification using the PureLink PCR Purification kit (Invitrogen, cat. no. K310002). The 30 µL DNA solution was mixed with 70 µL of water and 400 µL of binding buffer B2. When the mass of DNA exceeded 40 µg — the maximum binding capacity of the column —, the mixture of DNA and B2 was divided across more than one column. The rest of the purification was performed following the protocol provided by the manufacturer. DNA was eluted in 35 µL of elution buffer per column; samples divided into multiple columns were eluted into the same container.

Following purification, plasmid DNA was quantified using the Qubit™ 1x dsDNA high-sensitivity assay (Invitrogen, cat. no. Q33231) on a Qubit 4 fluorometer (Invitrogen, cat. no. Q33226). Samples were prepared by mixing 2 µL of a 100-fold dilution of plasmid DNA with 198 µL of Qubit working solution. Concentrations of plasmids measured using the Qubit assay were generally (40 to 50) % lower than those measured using a Nanodrop instrument. All cell-free reactions were prepared based on Qubit-measured concentrations.

Linear DNA templates were prepared *via* PCR as described above, purified using the PureLink PCR Purification kit, and quantified using the Qubit™ 1x dsDNA high-sensitivity assay for comparability to plasmid DNA concentrations. We found Qubit and Nanodrop concentrations were in much closer agreement for linear templates than for plasmids.

The few Medium Copy plasmids tested required two 100 mL LB cultures to be grown and combined onto a single column for plasmid extraction to obtain high enough DNA concentrations for use in multiple experiments. Because of that, High Copy plasmids were used for most constructs tested. Additionally, we found that MB and CFP measurements of Medium Copy plasmids encoding a riboregulator deviated more from linear templates in PURE than a High Copy counterpart (Supplementary Figures 5 and 13), even though concentrations were putatively the same across template types. These results could be due to increased carryover of inhibitors from Medium Copy plasmid extraction.

### Lysate preparation

The extract preparation protocol was adapted from published protocols^42,75^ and divided into a four-day protocol.

*Day 1*: An *E. coli* BL21 Star (DE3) glycerol stock was streaked onto an LB (10 g/L tryptone, 10 g/L sodium chloride, 5 g/L yeast extract) plate without antibiotic and incubated at 37 °C for 16 h.

*Day 2*: 50 mL of LB without antibiotic was inoculated with one colony from the *E. coli* BL21 Star (DE3) plate at 37 °C and 250 rpm for 16 h in a baffled 250 mL flask.

*Day 3:* 400 mL of 2xYTP (16 g/L tryptone, 5 g/L sodium chloride, 10 g/L yeast extract, 40 mmol/L potassium phosphate dibasic, 22 mmol/L potassium phosphate monobasic) was inoculated with 25 mL of the overnight culture at 37 °C, 250 rpm. About 1.5 h into growth, 0.4 mmol/L isopropyl β-d-1-thiogalactopyranoside (IPTG) was added to the culture to induce expression of T7 RNA polymerase. Cells continued to grow until the OD600 reached (1.6 to 1.7), about (3.5 to 4) h into growth. Upon reaching the target OD600, cells were harvested by centrifugation at 4 °C, 2700 x g and washed three times with S30 buffer (10 mmol/L Tris acetate, 14 mmol/L magnesium acetate, 60 mmol/L potassium acetate, pH adjusted to 8.2 with 5 mol/L potassium hydroxide, and 2 mmol/L dithiothreitol), with a centrifugation step between washes. After the last wash step, the mass of the cell pellet was determined, and cells were stored at −80 °C. Per 400 mL of culture, a cell mass of 2 g to 2.5 g is expected.

*Day 4:* Cell pellets were thawed on ice, then resuspended in 1 mL of S30 buffer per g of cells. The cellular resuspension was divided into 1 mL aliquots in sterile 1.5 mL microcentrifuge tubes. Cellular resuspensions were then lysed on ice using a Q125 sonicator (Qsonica) with a 3.175 mm diameter probe, 20 kHz frequency, 50 % amplitude, and cycles of 10 s on and 10 s off, delivering (260 ± 10) J after 5 cycles. At the start of each cycle, the tip of the probe was positioned close to the bottom of the tube to prevent bubble formation; approximately 5 times per cycle, the tube was moved down slowly such that the tip reached the 0.5 mL mark of the tube and then reverted to its original position. Immediately after lysis, 3 mmol/L DTT was added to each tube. Lysed cells were centrifuged at 4 °C, 12000 x g, for 15 min. The supernatant was consolidated into a single tube, mixed by inversion, then divided into 2 mL aliquots in 14 mL culture tubes and allowed to shake at 37 °C, 250 rpm, for 80 min—this is the runoff reaction. The sample was then centrifuged at 4 °C, 12000 x g, for 15 min. The supernatant (the final extract to be used in CFE reactions) was consolidated into one tube, mixed by inversion, divided into 200 µL aliquots, and stored at −80 °C for future use.

### Plate reader experiments

Unless otherwise specified, all cell-free reactions were run in three 10 µL technical replicates in clear, flat-bottom 384-well plates (Greiner Bio-one, cat. no. 784101) in a BioTek Synergy Neo2 plate reader at 37 °C. Measurements of sCFP3A (excitation: 430 nm, emission: 475 nm, bandwidth: 10 nm, gain: 85), MB (excitation: 524 nm, emission: 565 nm, bandwidth: 20 nm, gain: 80), and Pepper-HBC620 complex (excitation: 577 nm, emission: 620 nm, bandwidth: 20, gain: 75) were taken every 5 min from the bottom of the plate. All reactions included 5 µmol/L HBC620 Pepper dye and 500 µmol/L of the molecular beacon complex. DNA templates concentrations for each experiment are specified in the relevant figure captions or figure legends. At the end of each experiment a synthetic trigger for the molecular beacon, composed of 2′-O-methyl RNA bases, was added to each sample (final concentration 2.3 µmol/L) to obtain a maximum MB fluorescence signal for concentration calibration. The compositions of the different reaction environments are described below.

#### Cell-free reactions in lysate

Reactions were prepared as previously described^76^. Unless otherwise specified, reactions contained 17.5 mmol/L magnesium glutamate, 10 mmol/L ammonium glutamate, 133 mmol/L potassium glutamate, 1.2 mmol/L ATP, 0.85 mmol/L GTP, 0.85 mmol/L CTP, 0.85 mmol/L UTP, 0.034 mg/mL folinic acid, 0.171 mg/mL tRNA from *E. coli* MRE 600, 0.33 mmol/L nicotinamide adenine dinucleotide, 0.267 mmol/L coenzyme A, 4 mmol/L sodium oxalate, 1 mmol/L putrescine, 1.5 mmol/L spermidine, 50 mmol/L HEPES, 2 mmol/L each of the twenty standard amino acids, 0.03 mol/L PEP, 27 % extract by volume. 17.5 mmol/L magnesium glutamate was used because it yielded the highest ON/OFF ratio for CFP of magnesium glutamate concentrations spanning (10 to 30) mmol/L (Supplementary Figure 6).

#### PURExpress reconstituted system

Reactions were prepared following the protocol provided by the manufacturer. Components other than Solution A and Solution B were mixed on ice, followed by addition of Solution A (40 % by volume) then Solution B (30 % by volume).

#### PUREfrex reconstituted system

PUREfrex 2.1 reactions for were prepared following the protocol provided by the manufacturer. Cysteine was added at a final concentration of 0.3 mmol/L.

#### In vitro transcription (IVT)

Unless otherwise specified, IVT reactions were conducted in transcription buffer (40 mmol/L tris-HCl (pH 7.9), 6 mmol/L MgCl2, 10 mmol/L dithiothreitol, 10 mmol/L NaCl, and 2 mmol/L spermidine) supplemented with 2 mmol/L of each ribonucleotide triphosphate (NTP) (Thermo Fisher, cat. no. R0481). The concentrations of T7 RNAP (Thermo Fisher, cat. no. EP0113) used are specified in the relevant figure captions.

### THE riboregulator application experiments

#### Bidirectional regulation

CFP expression was activated or repressed by modulating input or output template concentrations, respectively. CFP expression was further tuned by modulating the length of the spacer domain *s* using G_1 (*s* = 4) or G_3 (*s* = 0). In these experiments, the gate DNA template concentration was 2 nmol/L and input or output template concentration varied from 0 nmol/L to 16 nmol/L with Io added to keep a constant 20 nmol/L of total template across all samples (Figure 6B). For the output strand, the DNA template was linear.

#### Thresholding-based logic

Input and gate DNA template concentrations were 4 nmol/L with Io added to keep a constant 20 nmol/L total template across samples. The concentration of the threshold gate DNA template was 5 nmol/L in the AND-like circuit and 0 nmol/L in the OR-like circuit. With exception of G{u1,rbs2}(G_3) and Io, linear DNA templates were used. G_3 (G{u1,rbs2}) lacked a spacer domain to provide a kinetic difference between its strand exchange rate and the threshold strand exchange rate.

#### Biosensing

Different versions of the TetR-responsive operator region were tested. Figure 6 experiments used an input template with a single copy of the operator downstream of the promoter because this configuration exhibited the largest fold signal repression by TetR (Supplementary Figure 33). For both positive and negative biosensors, the input DNA template concentration was 8 nmol/L and the gate DNA template concentration was 2 nmol/L. Io was not added. For the positive biosensor, the TetR-encoding template concentration was 4 nmol/L. For the negative biosensor, the TetR-encoding template concentration was 2 nmol/L, and this template was expressed in the PURE reaction for 16 min before the other DNA templates were added. The negative biosensor used a TetR mutant with 3 mutations: A56P, I57L, E58V^57^. Other mutants were screened, but exhibited weaker MB signal repression in the presence of aTc (Supplementary Figure 34).

### Protein purification

To generate a calibration curve for sCFP3A measurements, sCFP3A was purified in NEBExpress® Ni Spin Columns (New England Biolabs, cat. no. S1427L). Using inverse PCR, 6 histidine amino acid residues (CATCATCACCACCACCAT) were inserted at the C-terminus of sCFP3A in a plasmid encoding T7 RNA polymerase-driven sCFP3A expression and resistance to kanamycin.

BL21 Star (DE3) *E. coli* cells were transformed with the sCFP3A-6xHis plasmid, and one colony of this transformation was used to inoculate a 5 mL LB culture supplemented with 50 μg/mL kanamycin. This culture was grown at 37 °C, 250 rpm for 16 h. A 200 mL LB culture containing 50 μg/mL kanamycin was inoculated with 2 mL of the smaller culture and grown at 37 °C and 250 rpm. After 2 h of growth, 0.4 mmol/L IPTG were added to the culture to induce expression of T7 RNA polymerase. The culture was allowed to grow for another 2 h, after which cells were harvested by centrifugation at 4 °C, 2700 x g, 15 min. The cell pellet was resuspended in 1 mL of lysis buffer (supplied by the nickel column’s manufacturer) per g of cells. The cell resuspension was divided into 1 mL aliquots in 1.5 mL microcentrifuge tubes. Cells were then lysed on ice using a Q125 sonicator (Qsonica) with a 3.175 mm diameter probe, 20 kHz frequency, 50 % amplitude, and cycles of 10 s on and 10 s off, delivering about 300 J after 6 cycles. Lysed cells were centrifuged at 4 °C, 12000 xg, for 15 min, and the lysate supernatant was consolidated into one tube. Purification proceeded following the manufacturer’s protocol. 1 mL of lysate was added to each of two nickel columns, with 500 µL added at a time followed by binding and centrifugation. Each time, the lysate was allowed to bind to the column for 10 min at 4 °C, with gentle mixing by inversion every 3 min. Protein was eluted in 200 µL elution buffer. The purified protein fractions from each column were combined, and purification was verified on an SDS-PAGE gel (Bio-rad, cat. no. 4568096). The histidine tag was not removed.

Purified sCFP3A was quantified with the LabChip® GXII Touch™ protein characterization system (Revvity, CLS138160) and the ProteinEXact™ assay and lab chip (Revvity, CLS150337).

#### *In vitro* transcription

To generate a calibration curve for the Pepper aptamer, Pepper RNA transcripts were prepared via *in vitro* transcription (IVT) using a plasmid encoding T7 RNA polymerase-driven Pepper expression. Prior to transcription, this plasmid was linearized via PCR to remove the origin of replication and the antibiotic resistance gene.

The 100 µL IVT reaction contained 1 µg of linear DNA template, 2 mmol/L each of ATP, GTP, CTP and UTP, 0.6 U/µL T7 RNA polymerase (Thermo Scientific, cat. no. EP0113), and the transcription buffer provided by the polymerase’s manufacturer. The reaction was run in 100 µL volumes at 37 °C for 2 h in a thermocycler. Each reaction was then treated with 0.1 U/µL DNase I for 15 min at room temperature prior to purification and elution in 30 µL nuclease-free water using the RNA Clean & Concentrator-25 kit (Zymo Research, cat. no. R1018).

RNA was quantified using the Qubit™ RNA high-sensitivity assay (Invitrogen, cat. no. Q32855) on a Qubit 4 fluorometer (Invitrogen, cat. no. Q33226).

### Calibration curves for Pepper, molecular beacon, and sCFP3A measurements

For each calibration curve, the relevant molecule was added to PURE or lysate reactions without any DNA templates at multiple concentrations capturing fluorescence intensity values above and below those obtained in our experiments. The cell-free reactions were incubated in a plate reader at 37 °C and measured for 4 h. Fluorescence values at 90 min were used to create calibration curves for each molecule. Linear regression was used to fit the calibration curves using the polyfit() function of the Numpy package in Python 3.8.18. The slope of these fit lines was used to convert fluorescence intensities to concentrations. To avoid physically unrealistic negative concentrations, the minimum fluorescence value of a given sample was used as the value corresponding to a concentration of 0 mol/L.

#### Pepper aptamer

Purified Pepper mRNA prepared via IVT was used to generate calibration curves. In PURE, the signal stabilized after 1 h. In lysate, the signal decreased sharply in the first 30 min followed by a slow decay over the remainder of the experiment (Supplementary Figure 41A). Ribonuclease-mediated degradation in lysate confounds concentration calibration; the signal associated with an RNA concentration at a given time point may be used for comparing across samples and conditions, but does not reflect a stable RNA concentration during the reaction.

#### Molecular Beacon

A synthetic molecular beacon trigger strand, composed of 2omR bases, was added at different concentrations to PURE or lysate reactions containing a fixed concentration of the molecular beacon complex. The resulting calibration curves showed a linear trend between molecular beacon fluorescence and increasing concentrations of trigger RNA added (Supplementary Figure 41B). Thus, to convert molecular beacon signal to concentration the signal in each sample was normalized to be between 0 and 1 based on the minimum and maximum fluorescence values in an experiment and then multiplied by the total concentration of MB added to the reaction:

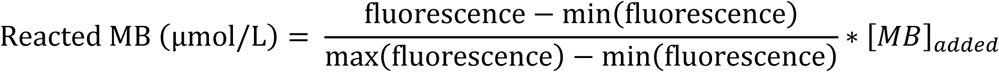

The synthetic molecular beacon trigger strand was spiked into every sample at the end of each experiment to a concentration in excess of the molecular beacon complex to ensure each sample had an internal measurement of fully reacted molecular beacon signal for normalization.

*sCFP3A:* Purified sCFP3A was used to generate calibration curves. In both PURE and lysate, the protein signal decreased slightly in the first 30 min, likely due to temperature equilibration of the samples in the plate reader, but then remained nearly constant (Supplementary Figure 41C).

## AUTHOR CONTRIBUTIONS (CRediT)

**FP**: Conceptualization (Template and experiment design), Methodology and Investigation (Plasmid cloning and preparation, lysate preparation, lysate experiments, PURE experiments), Project administration, Writing – original draft

**ER**: Methodology and Investigation (PURE experiments), Writing – review and editing

**SWS**: Conceptualization, Data curation, Formal analysis, Methodology and Investigation (Plasmid cloning, IVT experiments), Project administration, Supervision, Visualization, Writing – original draft

## DATA AND CODE AVAILABILITY

Fluorescence data for all experiments in this study will be available on Zenodo. DNA sequences, including their origin, are provided in Supplementary File S1. Annotated GenBank files of DNA templates are provided in Supplementary File S2. Annotated sequences are also available on <u>Benchling</u>.

Data analysis and visualization was conducted in Python (3.8.18) in the Spyder IDE (5.4.3). Example analysis and plotting code is available in Supplementary File S3. Additionally, the ctRSD simulator 2.1 package (https://ctrsd-simulator.readthedocs.io/en/latest/SeqCompiler.html) contains a sequence compiling function to stitch together sequences for any combination of sequence domains explored in this study. Scripts for generating the sequences in this study using the sequence compiler are available in Supplementary File S3. A Google CoLab for running the sequence compiler is available at: <u>CoLab link</u>

## ACKNOWLEGDMENTS

The authors thank Svetlana Ikonomova, Zoila Jurado, David Ross, Elizabeth Strychalski, Chad Sundberg, and Molly Wintenberg for insightful discussions throughout the project and feedback on the manuscript. In addition, the authors thank Katharina Yandrofski and Kayla Diaz at the Institute for Bioscience and Biotechnology Research (IBBR) for usage of and assistance with their LabChip® GXII Touch protein characterization instrument. The authors also thank Bright Eyes, whose album *I’m Wide Awake, It’s Morning* was inspiration throughout the study.

## Disclaimer

Certain commercial entities, equipment, or materials may be identified in this document to describe an experimental procedure or concept adequately. Such identification is not intended to imply recommendation or endorsement by the National Institute of Standards and Technology, nor is it intended to imply that the entities, materials, or equipment are necessarily the best available for the purpose. Official contribution of the National Institute of Standards and Technology; not subject to copyright in the United States.

