## Supplementary Figures for "Cell-free systems enable mechanistic characterization of genetically encoded RNA strand exchange circuits for programmable protein expression, logic, and biosensing"

### Supplementary Information

*Disclaimer:* Certain commercial entities, equipment, or materials may be identified in this document to describe an experimental procedure or concept adequately. Such identification is not intended to imply recommendation or endorsement by the National Institute of Standards and Technology, nor is it intended to imply that the entities, materials, or equipment are necessarily the best available for the purpose. Official contribution of the National Institute of Standards and Technology; not subject to copyright in the United States.

#### 1 Overview of components and component nomenclature

Supplementary Figs. 1 and 2 provide an overview of the full nomenclature for all components in this study. This nomenclature, albeit complicated, uniquely describes the sequence identity of every component. The sequences of each domain, separated in domain specific tabs are provided in `ctRSD_domains_list_210.xls` and at: https://github.com/usnistgov/ctRSD-simulator/blob/main/ctRSD-simulator-2.0/Sequence%20Compiler/ctRSD_domains_list_210.xls

The full nomenclature can also be used to generate component sequences for any combination of domains using sequence compiler software we developed in Python. The full documentation for the sequence compiler can be found at:
https://ctrsd-simulator.readthedocs.io/en/latest/SeqCompiler.html

A Google CoLab for running the sequence compiler is available at: CoLab link

All gene fragment sequences ordered for this manuscript are in Supplementary File S1.

GenBank files of all plasmids used in this study are in Supplementary File S2.

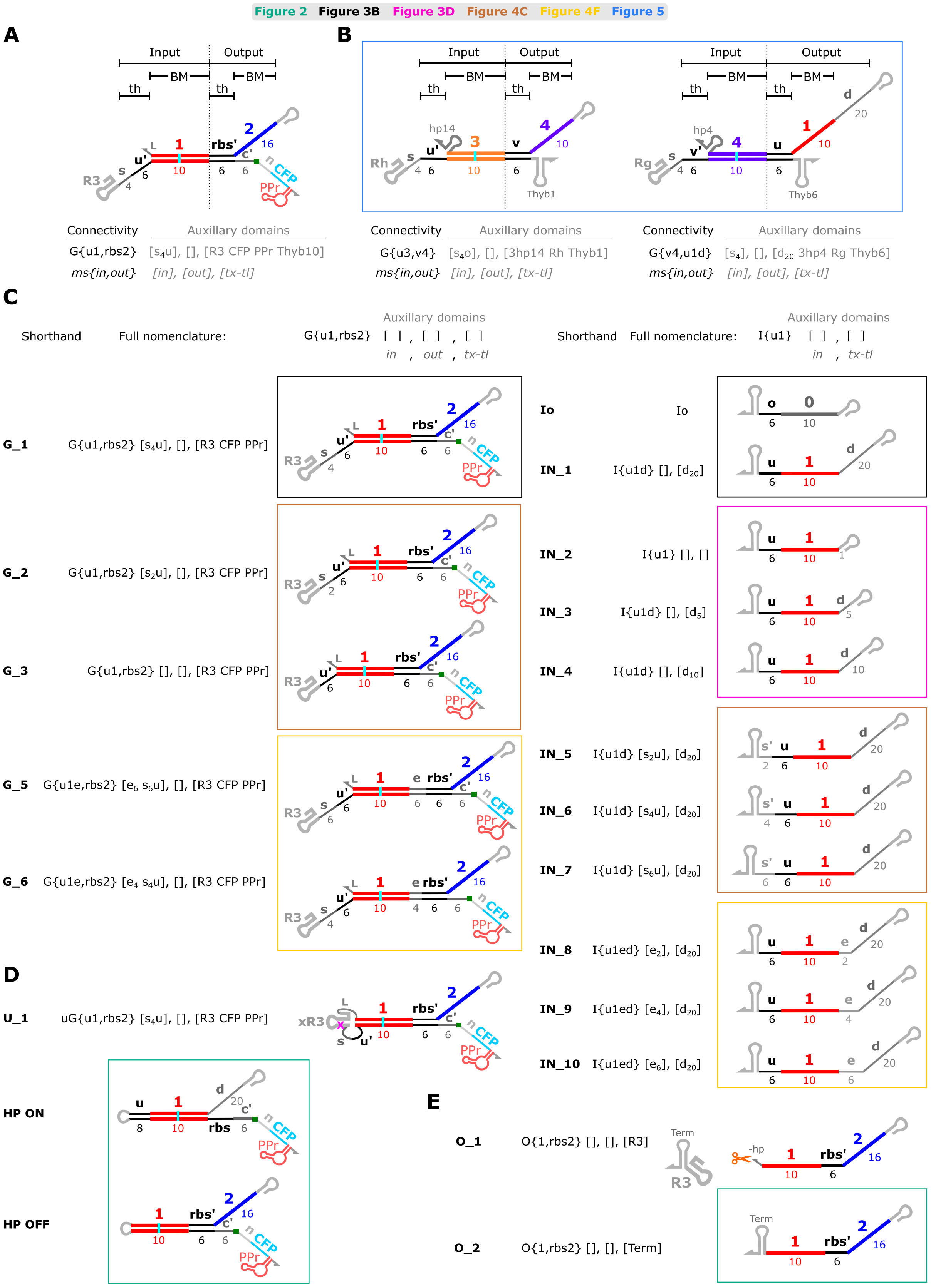
Figure 2 Figure 3B Figure 3D Figure 4C Figure 4F Figure 5
A
Input Output
BM BM
th th
Connectivity
Auxillary domains
G{u1,rbs2}
[s4u], [], [R3 CFP PPr Thyb10]
ms{in,out}
[in], [out], [tx-tl]
B
Connectivity
Auxillary domains
G{u3,v4}
[s4o], [], [3hp14 Rh Thyb1]
ms{in,out}
[in], [out], [tx-tl]
Connectivity
Auxillary domains
G{v4,u1d}
[s4], [], [d20 3hp4 Rg Thyb6]
ms{in,out}
[in], [out], [tx-tl]
C
Shorthand
Full nomenclature:
G{u1,rbs2} [ ] , [ ] , [ ]
in , out , tx-tl
Shorthand
Full nomenclature:
I{u1} [ ] , [ ]
in , tx-tl
G_1 G{u1,rbs2} [s4u], [], [R3 CFP PPr]
G_2 G{u1,rbs2} [s2u], [], [R3 CFP PPr]
G_3 G{u1,rbs2} [], [], [R3 CFP PPr]
G_5 G{u1e,rbs2} [e6 s6u], [], [R3 CFP PPr]
G_6 G{u1e,rbs2} [e4 s4u], [], [R3 CFP PPr]
Io Io
IN_1 I{u1d} [], [d20]
IN_2 I{u1} [], []
IN_3 I{u1d} [], [d5]
IN_4 I{u1d} [], [d10]
IN_5 I{u1d} [s2u], [d20]
IN_6 I{u1d} [s4u], [d20]
IN_7 I{u1d} [s6u], [d20]
IN_8 I{u1ed} [e2], [d20]
IN_9 I{u1ed} [e4], [d20]
IN_10 I{u1ed} [e6], [d20]
D
U_1 uG{u1,rbs2} [s4u], [], [R3 CFP PPr]
HP ON
HP OFF
E
O_1 O{1,rbs2} [], [], [R3]
O_2 O{1,rbs2} [], [], [Term]

**Supplementary Figure 1:** Nomenclature and schematics of toehold exchange (THE) riboregulators and cotranscriptionally encoded RNA strand displacement (ctRSD) species used in this study. Connectivity species names based on how species connect to one another are used in the figures. The shorthand and full names with auxiliary domains are used in Supporting File S1 to give each species a unique name in absence of accompanying schematics. Colored boxes around clusters of species highlight which main text figures the species are introduced in. (**A**,**B**) Schematics of a THE riboregulator gate (A) and upstream ctRSD gates (B). THE riboregulator and ctRSD gates are composed of input and output domains, which each possess two subdomains: a toehold (th) subdomain for initiating strand displacement reactions and a branch migration (BM) subdomain for strand displacement specificity. Bold numbers and letters above the gate represent domain sequence identity and nonbold numbers below the gate represent domain length in bases. Connectivity names specify the molecular species (ms: G{ } for gate, I{ } for input, O{ } for output, *etc*.) and how it is connected to other species with input and output domains specified within the first and second positions inside the curly brackets, separated by a comma. Auxiliary domains, which are specified after the connectivity name, are not related to how species are connected. The convention is to group different types of auxiliary domains together and separate these groupings by a comma. The first grouping is for domains related to the input portion of the species (input extensions (*e*) and spacers / extended toeholds (s)), the second grouping is for domains related to the output portion of the species (output extensions (*e*)), and the last grouping is for domains related to transcriptional encoding or translation properties of the species. Note the *d* domain is included in the connectivity name because it *implies* a connection to a THE riboregulator, but its identity is specified in the third auxiliary group because its role is related to protein translation and it is not involved in base pairing interactions between species or expected to influence the kinetics of reactions between RNA species. Within auxiliary groupings [ ], the convention is to list domains in the 5′ to 3′ order they appear within the species. For auxiliary domains that can be different lengths *e.g.*, *s*, *e*, *d*, *etc*., the convention is to specify the domain type first, the length in bases second as a subscript, and any sequence identifier last. For example, $s_4u$ is a spacer that is 4 bases in length and an extension of the *u* toehold. Across connectivity and auxiliary domain names, numbers in subscripts refer to the length of the domain in bases and all numbers not in subscripts are part of the sequence identifier. Specific subdomain sequences are part of the ctRSD sequence compiler and listed in the `ctRSD_domains_list_210.xlsx` file. (**C**) Schematics of prominent gates and inputs used in this study. (**D**,**E**) Additional control constructs used in the study. (**D**) U_1 contains a point mutation in the ribozyme sequence that abolishes cleavage activity, serving as a "no toehold exchange" control. HP OFF and HP ON are constructs that mimic the OFF and ON THE gate constructs but without the added complexity of the ribozyme or the need for strand exchange. HP OFF serves as a control for the lowest level of CFP expression for an OFF gate, as it should have a stable hairpin stem sequestering the *rbs*. HP ON serves as a control for the highest level of CFP expression from an ON gate, as it is immediately translated upon being transcribed. (**E**) O_1 and O_2 are two transcription controls used to calibrate molecular beacon measurements. Both mimic direct production of the output of a THE gate. O_1 has the self-cleaving ribozyme at the 3′ end to match the exact structure of the output of a THE gate. O_2 ends in the 3′ terminator structure.

For brevity a few domains that are not changed, or rarely changed, in this study are omitted in the full nomenclature. All the components in this study have the same 5′ hairpin (5hp) and, unless otherwise stated, the same terminator (Thyb10), so these are not explicitly listed in the above nomenclature for each component. The short linker (L) domain adjacent to the ribozyme is not explicitly defined for each component above as the default is to use the same 3-base linker. If the linker (L) is changed to something else, such as a hairpin-forming sequence *e.g.,* 3hp14, 3hp4, this will be specified in the third [tx-tl] bracket of the auxiliary domains.

**A**

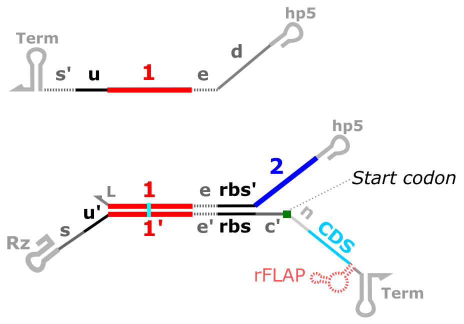

**B**

| Domain name | Length | Description |
|---|---|---|
| *hp5* | *Variable* | *5' hairpin:*<br>*Promote uniform transcription rates and stability* |
| *d* | *1-20* | *Distal domain:*<br>*Provide ssRNA standby site for ribosome binding* |
| * *e* | *0-6* | *Extended branch migration domain:*<br>*Extend dsRNA duplex or output toehold* |
| *1,2 (3,4,N)* | *10* | *Branch migration domain:*<br>*Promote strand exchange specificity* |
| *u (v,w,rbs')* | *6* | *Toehold:*<br>*Facilitate strand exchange* |
| *rbs* | *6* | *Ribosome binding sequence:*<br>*Interacts with 16S rRNA of ribosome to facilitate translation* |
| *rbs'* | *6* | *rbs complementary domain:*<br>*Sequester rbs to prevent translation* |
| *Term* | *Variable* | *Terminator:*<br>*Terminate transcription* |
| *s* | *0-10* | *Spacer:*<br>*Reduce steric hindrance for input binding* |
| * *s'* | *0-10* | *Spacer complementary domain:*<br>*Extend the input toehold length* |
| *Rz (R3,Rg,Rh,Rm)* | *60-100* | *Self-cleaving ribozyme:*<br>*Produce dsRNA gates* |
| *L* | *3* | *Ribozyme linker:*<br>*Facilitate efficient cleavage; or a hairpin for ouput stability* |
| *c'* | *6* | *rbs to AUG linker:*<br>*Optimal spacing for translation initiation* |
| *n* | *18* | *N-terminal linker:*<br>*Provide ssRNA footprint for ribosome* |
| * *rFLAP* | *Variable* | *RNA fluorescent aptamer:*<br>*Measure transcription and RNA stability* |
| *CDS* | *Variable* | *Protein coding sequence:*<br>*Encodes for protein expression after strand exchange* |

**Supplementary Figure 2:** Definitions and functions of the domains of toehold exchange riboregulators and their inputs used in this study. (**A**) Schematics of an input and THE riboregulator with the relevant domains from this study shown. The domains with dashed lines are optional and not necessary or not conventionally used in the canonical THE riboregulator design (Supplementary Figure 1A). (**B**) Domain definitions and desired functions, along with their typical lengths in bases. The domain names with * to their left correspond to the non-necessary domains illustrated with dashed lines in (A). The domain names here match those used in the ctRSD sequence compiler (https://ctrsd-simulator.readthedocs.io/en/latest/SeqCompiler.html).

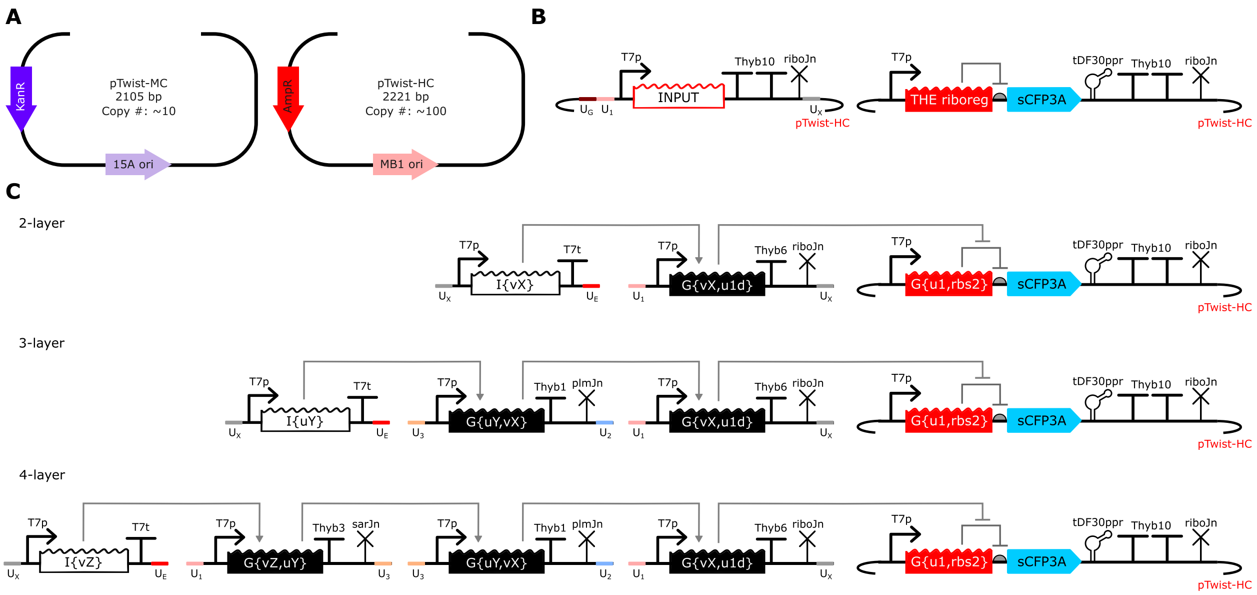

**Supplementary Figure 3**: Design of DNA templates encoding THE riboregulators, inputs, and other circuit domains. Most DNA templates used in this work were synthetized by Twist Biosciences to facilitate sharing and purchasing DNA. In these DNA templates, TMSE circuit components were encoded on plasmid backbones designed by the company. (**A**) Plasmid backbones from Twist Biosciences. Unless otherwise specified, a high-copy backbone (pTwist-HC) was selected to enable plasmid extraction at high yields. Additional control experiments included a medium-copy backbone (pTwist-MC). (**B**) Insert schematics for inputs and THE riboregulators placed on the Twist backbones in (A). (**C**) Schematics of templates for multi-layer cascades. For multi-layer cascades, the templates upstream of the THE riboregulator were encoded on linear templates. The colored $U_i$ domains indicate homology domains from[1] used for cloning these templates in other projects[2]. Thyb10 is an efficient T7 RNAP terminator characterized in[3].

#### 2 Identifying suitable experimental conditions

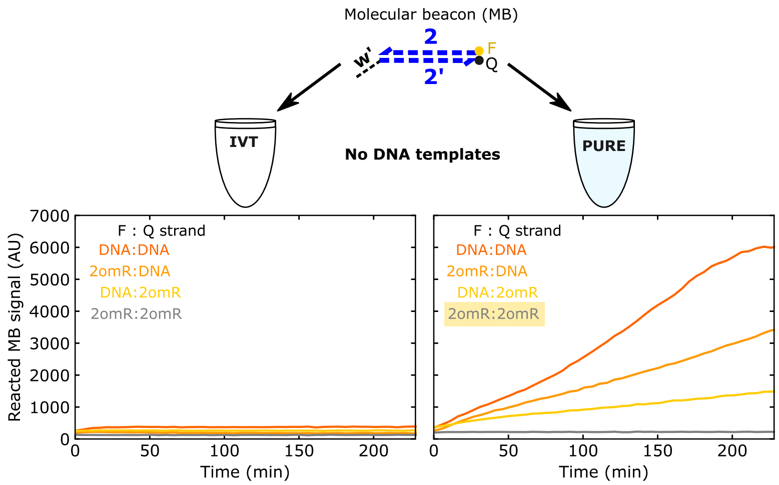

**Supplementary Figure 4**: DNA-based molecular beacons are not suitable for use in PURE. In *in vitro* transcription (IVT) reactions lacking DNA templates or trigger strands (left), a molecular beacon generated negligible signal regardless of whether the strands were DNA or 2′-O-methyl RNA (2omR). In PURE reactions (right), however, molecular beacons with at least one DNA strand generated measurable signals. Both strands of the molecular beacon must be made of 2omR – used in this study – to completely remove the spurious signal observed in PURE. In both experiments, 500 nmol/L of molecular beacon was used.

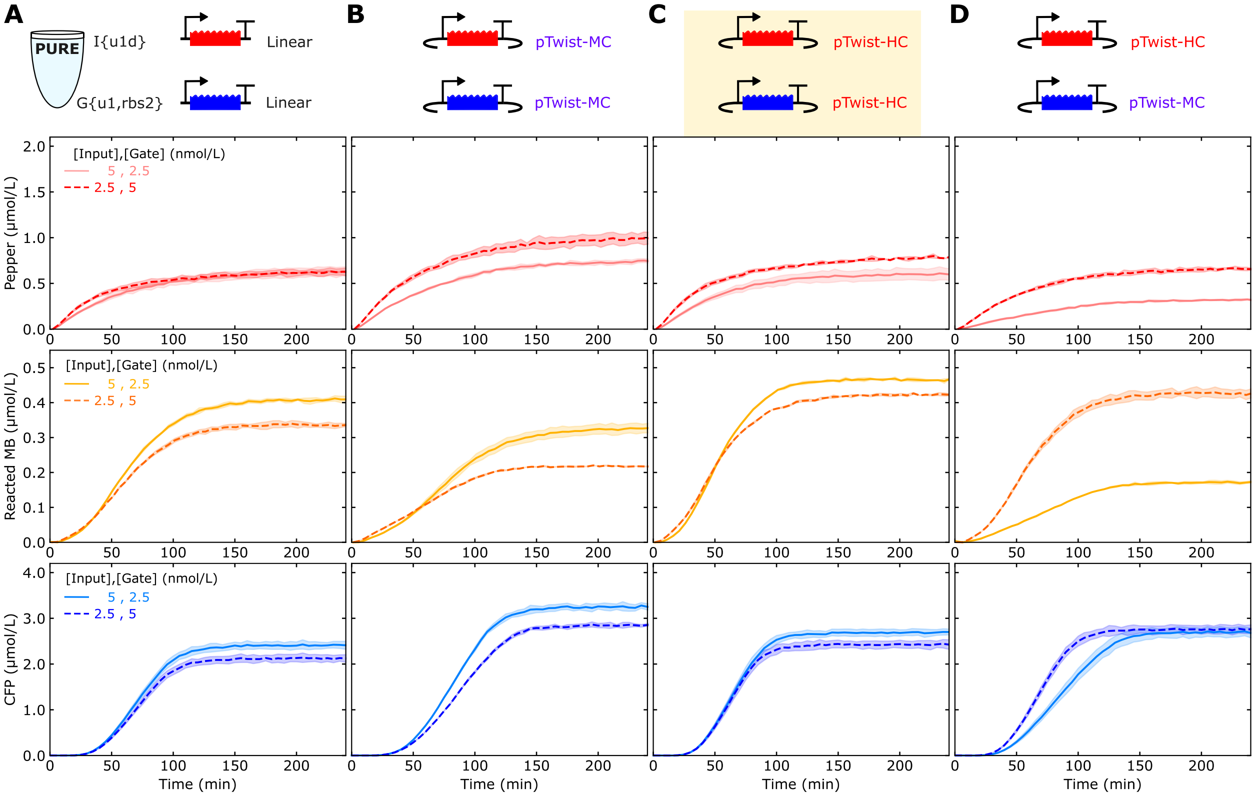

**Supplementary Figure 5**: Extracted plasmids with high-copy backbones produce results closer to linear templates in PURE than plasmids with medium-copy backbones. The input and the THE riboregulator were cloned onto either a high-copy backbone (pTwist-HC) or a medium-copy backbone (pTwist-MC). The plasmid designs shown in the panel highlighted, (C), were selected for use in this study.

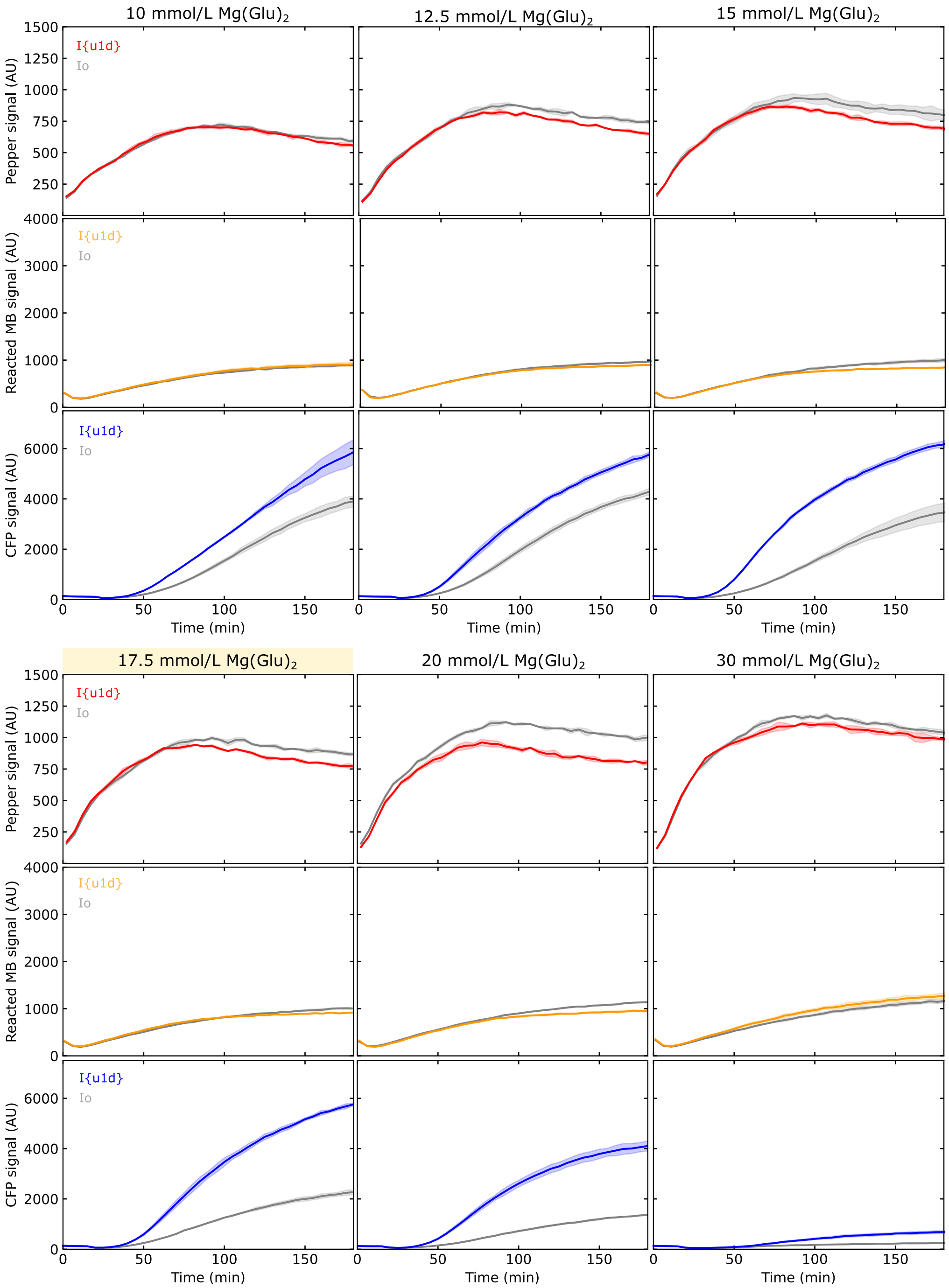

**Supplementary Figure 6**: The concentration of magnesium glutamate in lysate reactions influences CFP ON/OFF ratio for THE riboregulators. Input (IN_1) and THE riboregulator (G_1) DNA templates were both at 5 nmol/L. 17.5 mmol/L of magnesium glutamate was used for the rest of the study (highlighted panel).

### 3 Control measurements across environments

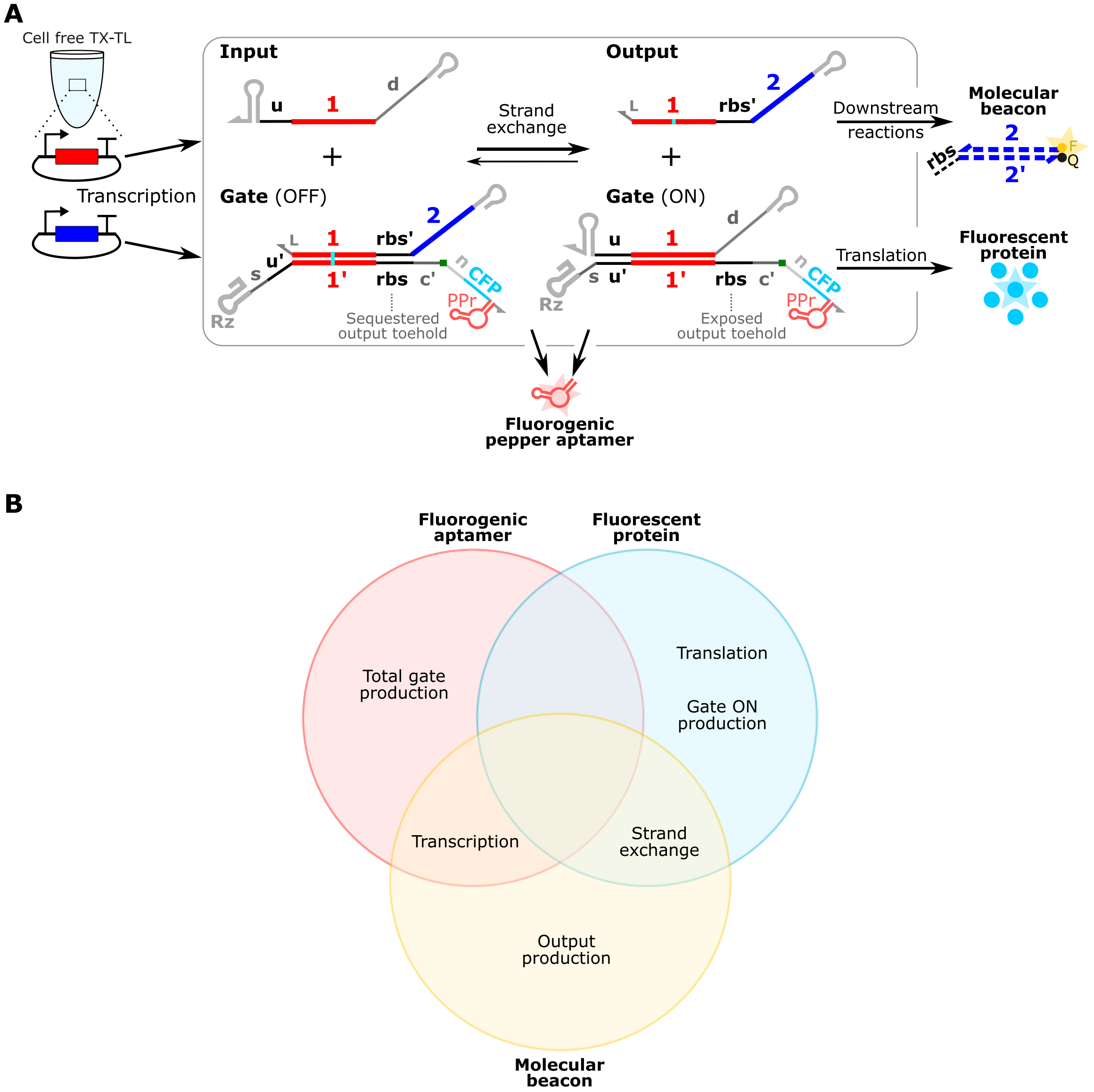

**Supplementary Figure 7:** Overview of THE riboregulator measurements. **(A)** Schematic of THE riboregulator production, operation, and measurement, reproduced from Figure 1A. This scheme allows fluorescence measurements from a fluorogenic aptamer, a molecular beacon, and a fluorescent protein. **(B)** Processes captured by these three measurements. No process is captured by all three measurements, demonstrating the need for multiple measurements. Some processes can be captured by more than one measurement, providing additional insight into the process based on how the two measurements correlate.

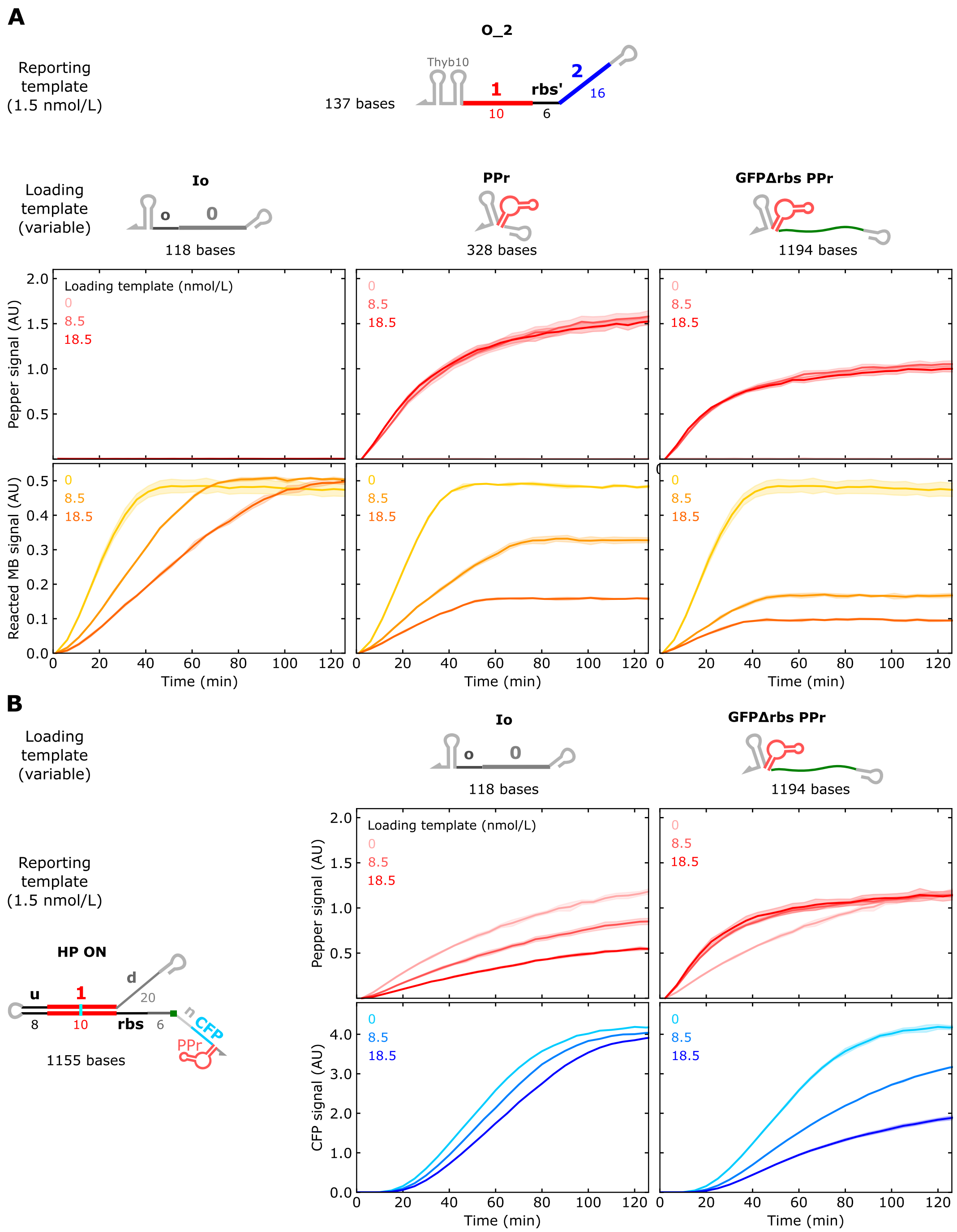

**Supplementary Figure 8**: Increasing the total template concentration decreases the transcription rate of individual components. For a fixed concentration (1.5 nmol/L) of the reporting template, different DNA loading templates were added at varying concentrations to reach different total DNA concentrations (0 nmol/L, 10 nmol/L, and 20 nmol/L). Reporting templates encoded O_2 or HP ON in (A) and (B), respectively.

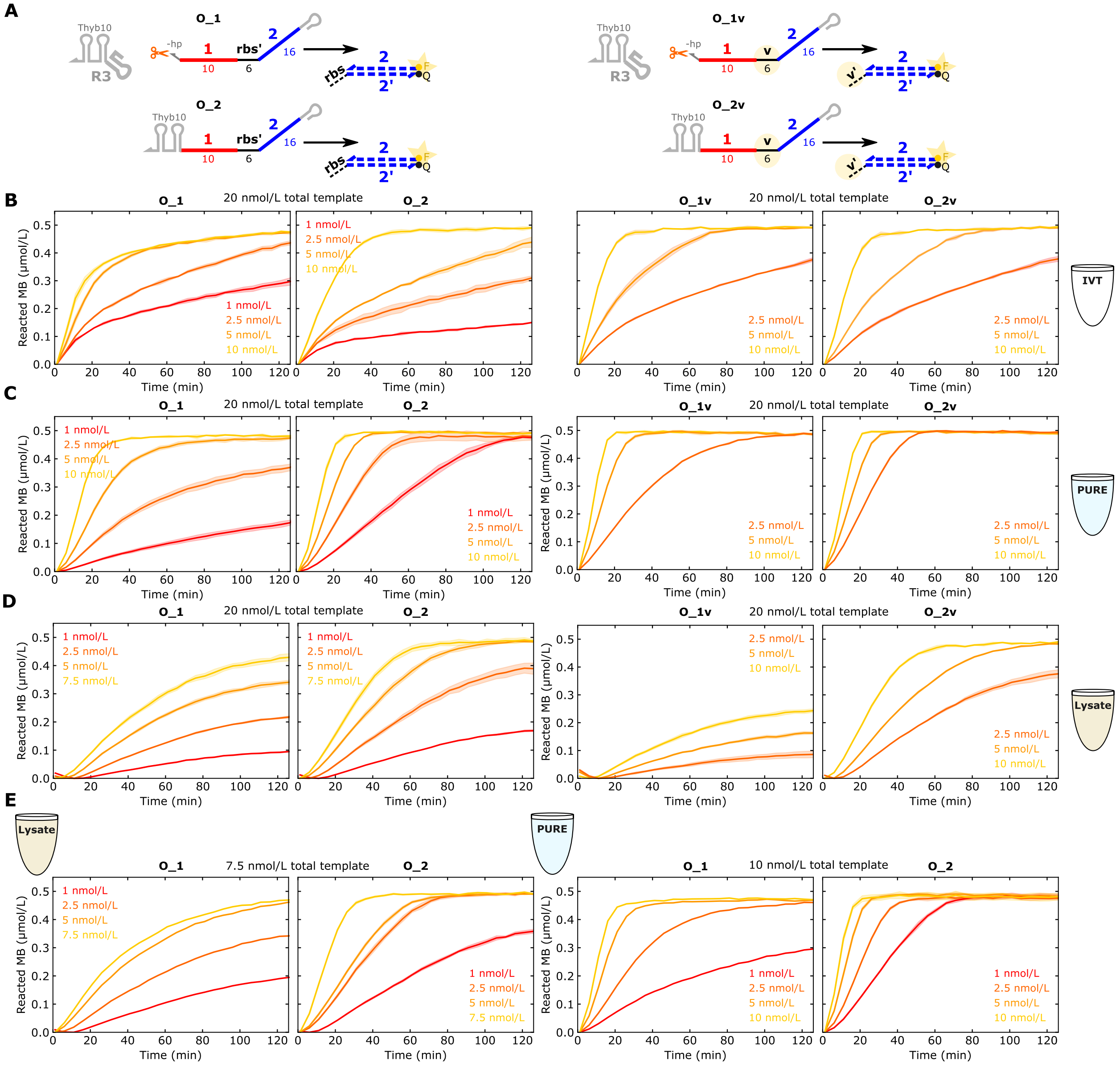

**Supplementary Figure 9**: Measuring transcription with molecular beacons. **(A)** Schematic of templates mimicking the output of a THE riboregulator. O_1 has a self-cleaving ribozyme at the 3′ end to match the exact structure of the output of a THE gate, whereas O_2 ends in the 3′ terminator structure. O_1v and O_2v mimic O_1 and O_2, respectively, but encode a *v* output toehold sequence instead of the *rbs* sequence. **(B-E)** Molecular beacon measurements for the templates described in (A) at different concentrations and in different environments. Io was added to the reaction to reach the total DNA template concentration indicated at the top of each panel. The O_2 measurements in (C) and (D) are shown in Figure 2D of the main text. In IVT experiments in (B), 5 U/µL T7 RNAP was used.

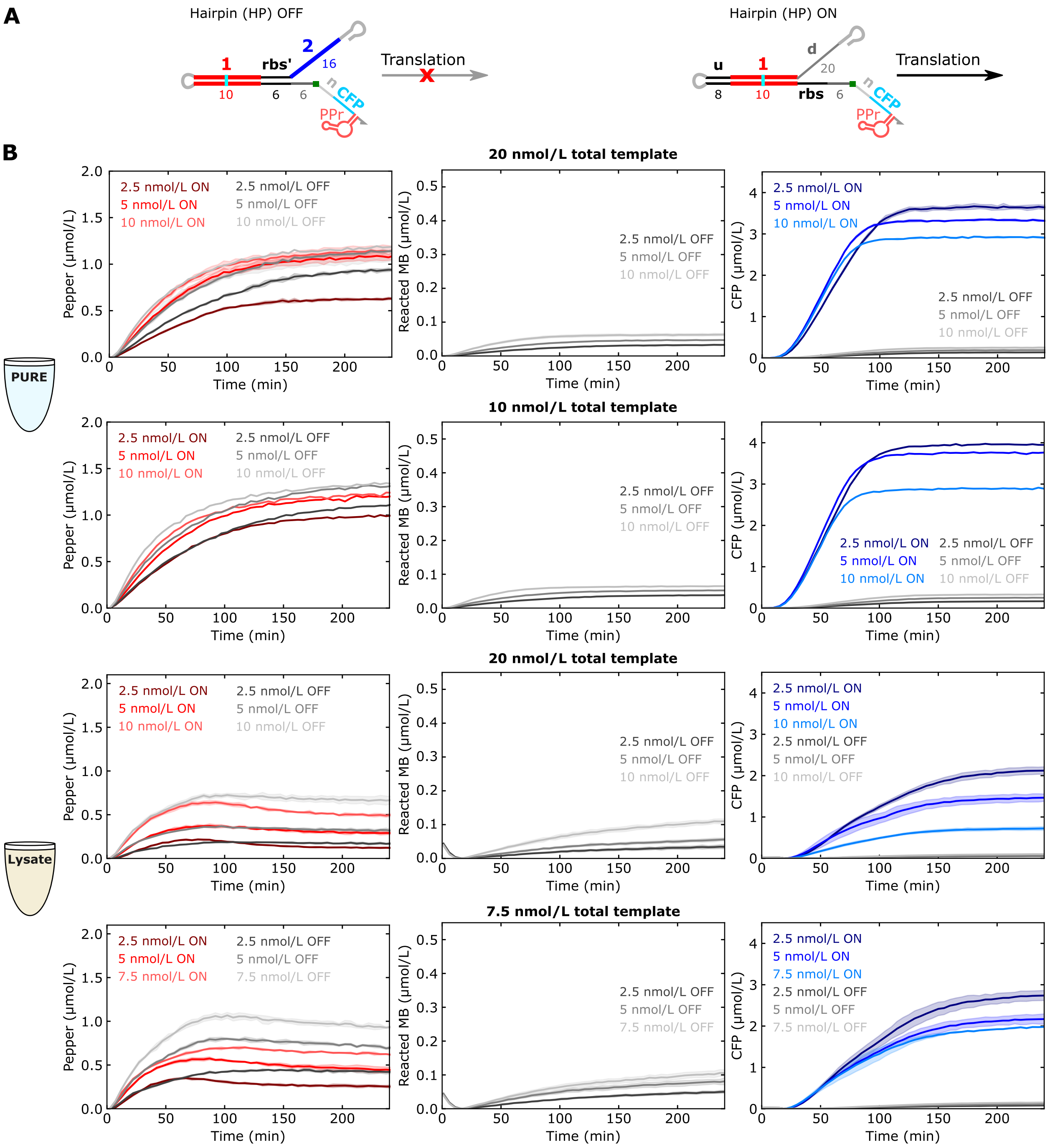

**Supplementary Figure 10**: Translation controls. **(A)** Schematic of constructs mimicking the OFF and ON THE gates (HP OFF and HP ON, respectively) without the added complexity of the ribozyme or the need for strand exchange. **(B,C)** Measurements of Pepper, molecular beacon, and CFP signals for different concentrations of HP OFF and HP ON in **(B)** PURE and **(C)** lysate. Io was added to each reaction to reach the total DNA template concentration indicated at the top of each panel. The 20 nmol/L total DNA template concentrations measurements are also shown in Figure 2D of the main text.

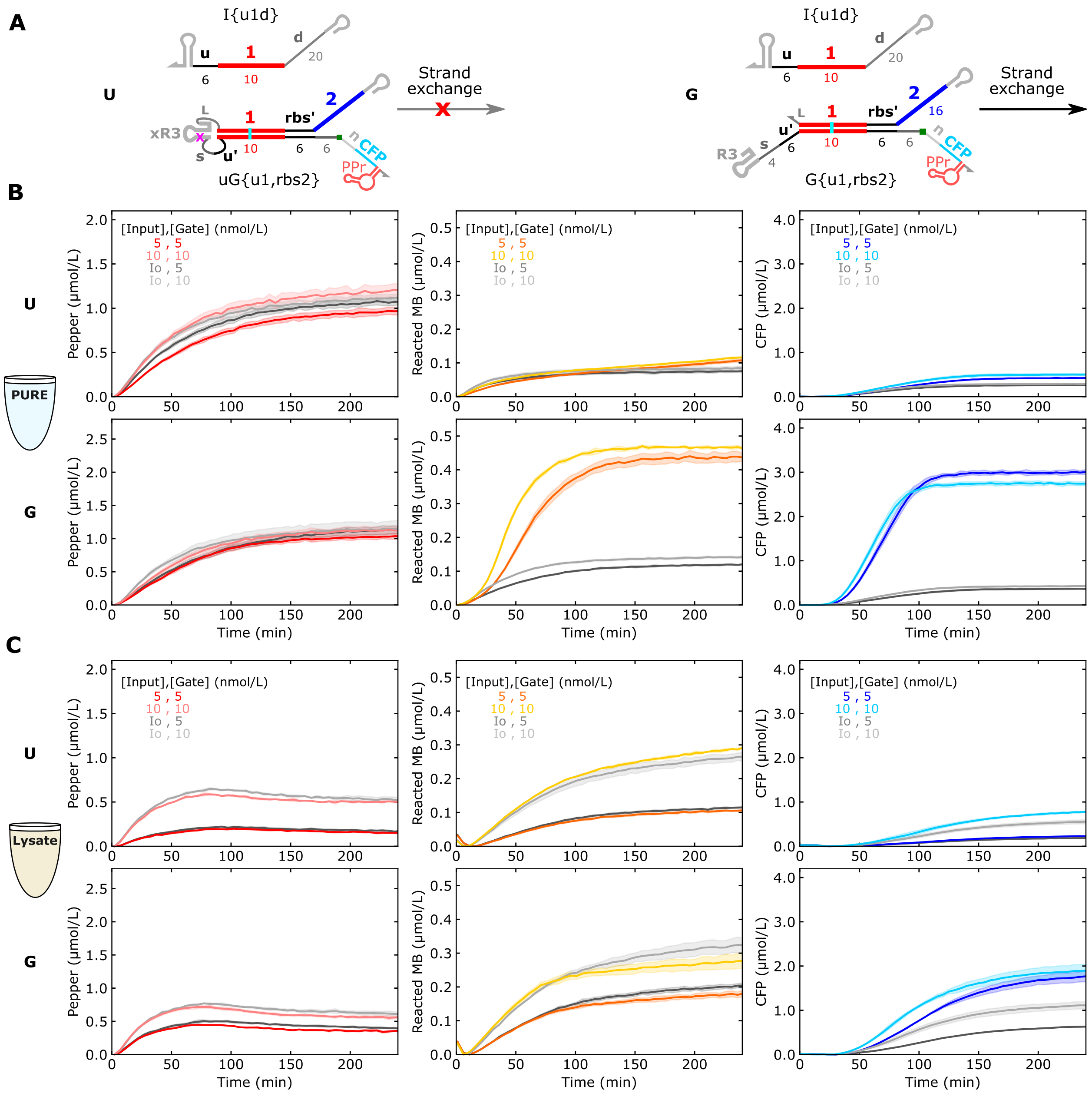

**Supplementary Figure 11**: THE riboregulators with a mutated ribozyme that cannot cleave do not produce appreciable molecular beacon and fluorescent protein signals. **(A)** Schematic of THE riboregulators without (left) and with (right) a functional ribozyme. Strand exchange is not expected to occur when the ribozyme cannot cleave the gate. **(B,C)** Measurements of Pepper, molecular beacon, and CFP signals in **(B)** PURE and **(C)** lysate. In both CFES, THE riboregulators with a non-functional ribozyme have molecular beacon and CFP signals close to background. As expected, the Pepper signal is not affected by ribozyme function. A total template concentration of 20 nmol/L was used in all experiments.

### 4 Additional experiments supporting main text figures

#### 4.1 Additional permutations of THE riboregulator designs and concentrations

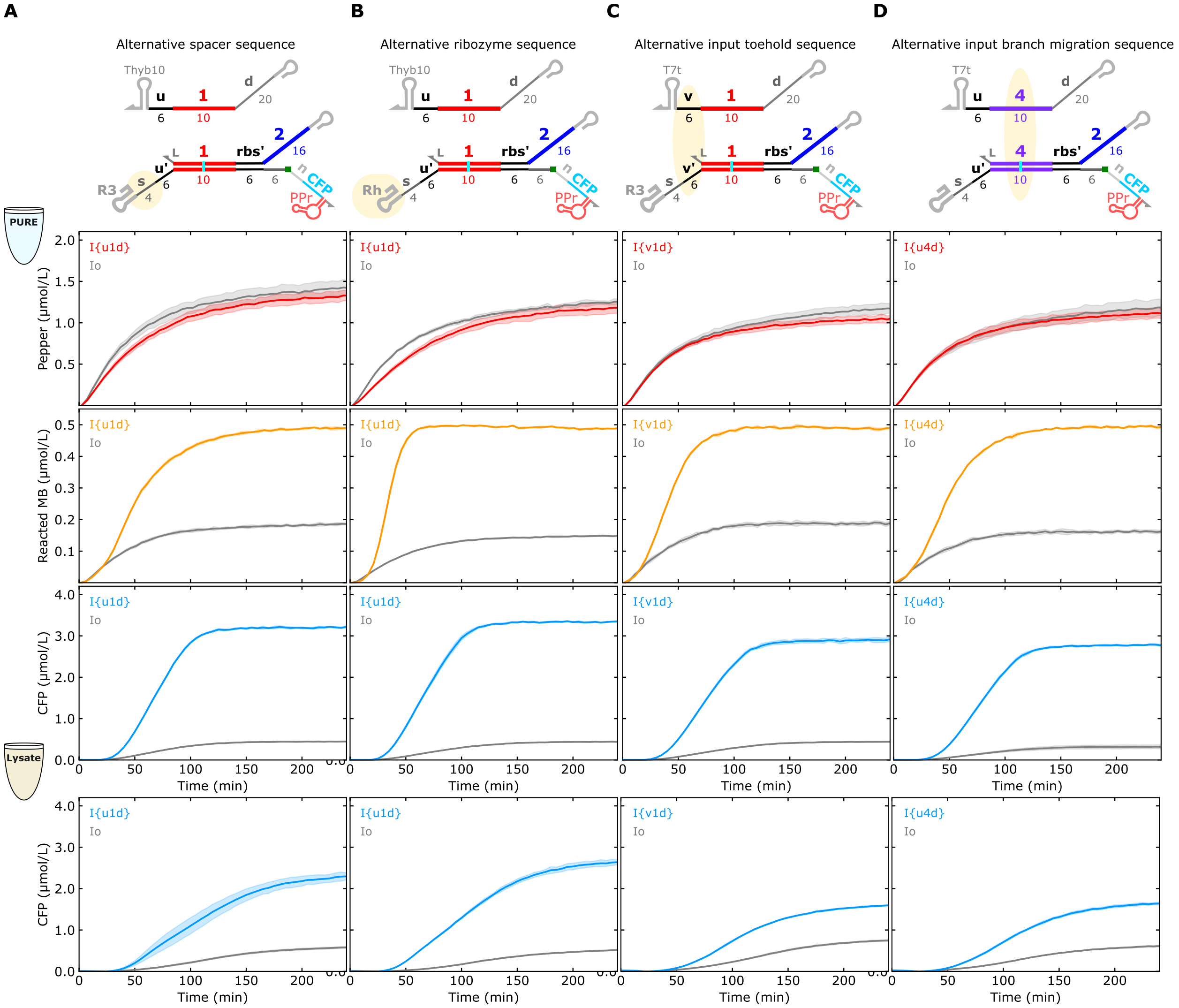

**Supplementary Figure 12:** Sequence variants of gates show similar performance. The altered domains, relative to the design used in Figure 2 of the main text, are highlighted in the schematics above the plots. In PURE, the input and gate DNA templates were both 5 nmol/L. In lysate, the input DNA template was 5 nmol/L, and the gate DNA template was 2.5 nmol/L.

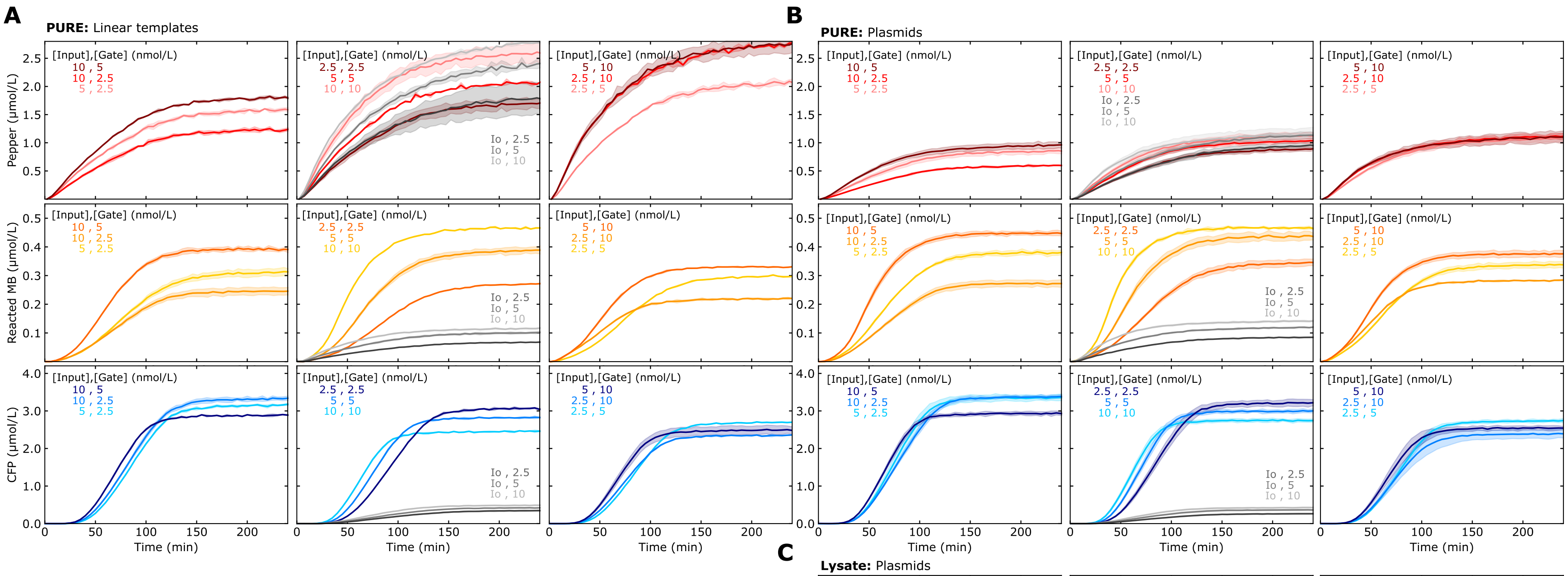

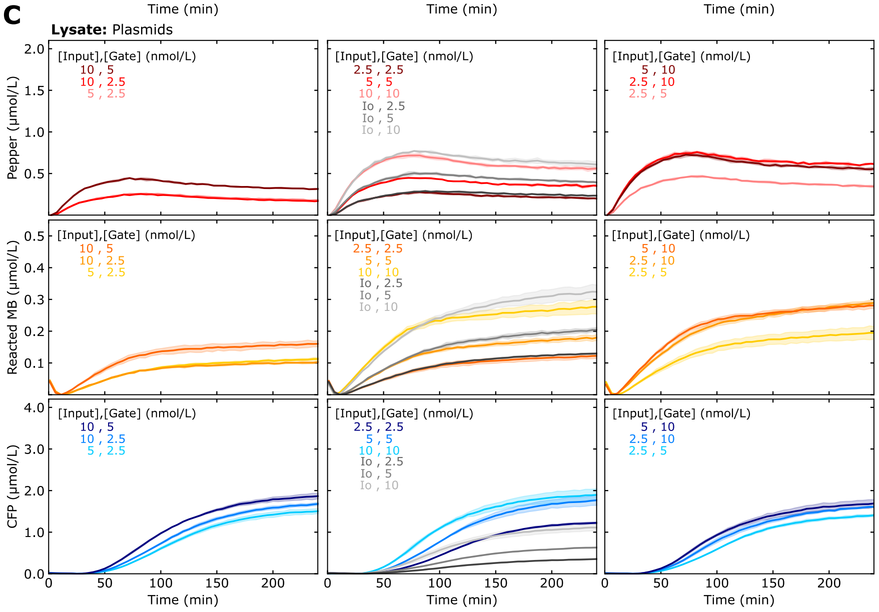

**Supplementary Figure 13:** Tuning protein expression by changing input and gate template concentrations for **A)** linear DNA templates in PURE, **(B)** plasmids in PURE, **(C)** plasmids in lysate. Input and gate DNA template concentrations are labeled in each plot with Io added to keep a constant 20 nmol/L total DNA template across samples. Measurements for reactions with equal concentrations of input (I{u1d} [], [d$_{20}$]) and gate (G{u1,rbs2} [s$_4$u], [ ], [R3]) templates are shown in Figure 2 of the main text.

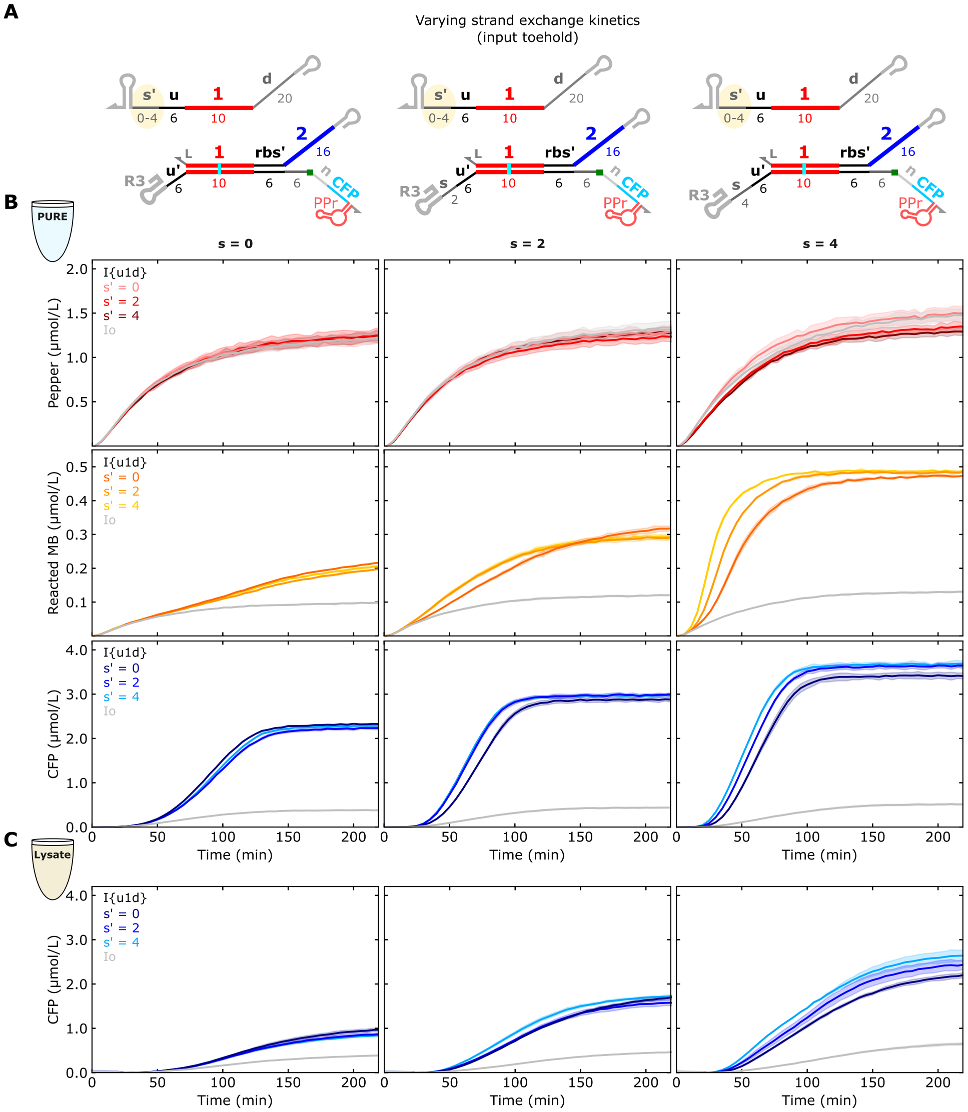

**Supplementary Figure 14:** Tuning protein expression by modulating RNA strand exchange kinetics, specifically via the input toehold length. **(A)** Schematics of inputs with varying input toehold ($s'$) lengths and THE riboregulators with different input toehold ($s$) lengths. **(B,C)** Measurements of Pepper, molecular beacon, and CFP signals in (B) PURE and (C) lysate for the inputs described in (A) combined with each THE riboregulator. In PURE, the input and gate DNA templates were both 5 nmol/L. In lysate, the input DNA template was 5 nmol/L, and the gate DNA template was 2.5 nmol/L. Measurements for reactions where $s$ = 4 are also shown in Figure 4C,D of the main text.

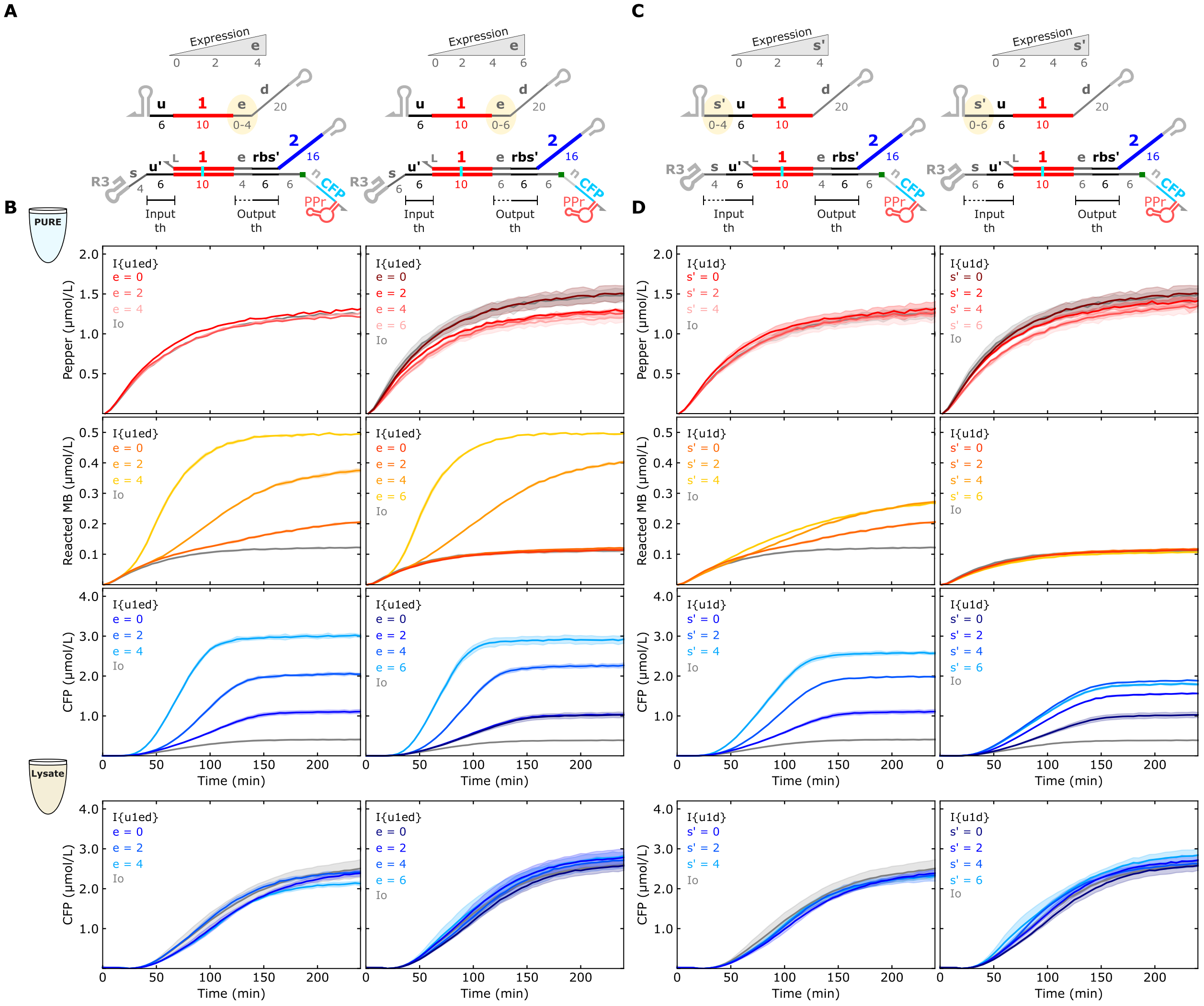

**Supplementary Figure 15:** Tuning protein expression by modulating RNA strand exchange kinetics, specifically via the output toehold length. **(A)** Schematics of inputs with varying *e* domain lengths, inputs with varying input toehold (*s′*) lengths, and a THE riboregulator with a 6-base *e* domain and a 6-base *s* domain. **(B,C)** Measurements of Pepper, molecular beacon, and CFP signals for the systems described in (A) in (B) PURE and (C) lysate. In PURE, the input and gate DNA templates were both 5 nmol/L. In lysate, the input DNA template was 5 nmol/L, and the gate DNA template was 2.5 nmol/L. A 6-base *e* domain shows similar trends to a 4-base *e* domain. The right panel of (B) is shown in Figure 4F,G of the main text.

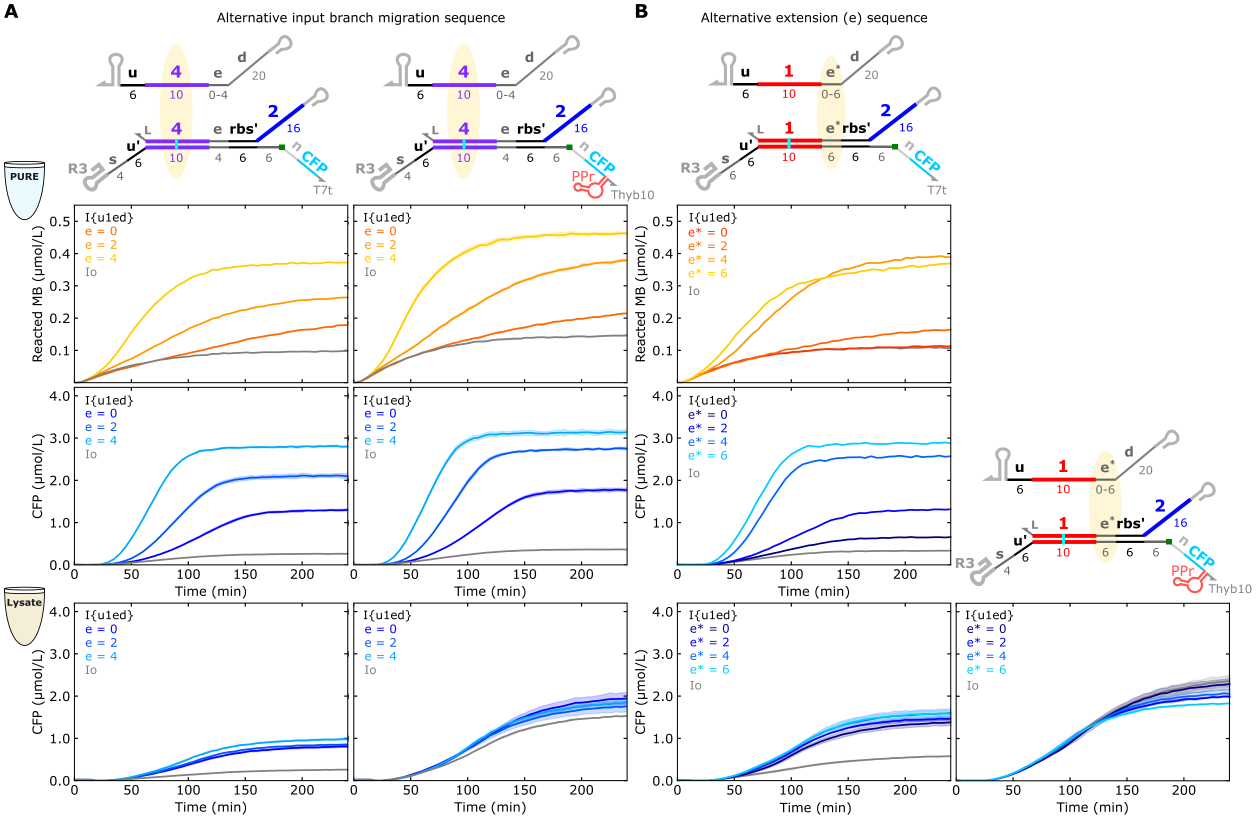

**Supplementary Figure 16:** Sequence variants of gates with extended input branch migration domains. (**A**) A different branch migration sequence (*4* instead of *1*) with two different 3′ UTRs on the gates. (**B**) A different e domain (*e**) with two different 3′ UTRs on the gates. PURE experiments in (B) were a single replicate. For $e^* = 4$, the *d* domain unintentionally adds a complementary base ($e^*+d = 5$), hence the results are closer to $e^* = 6$ than intended. In PURE, the input and gate DNA templates were both 5 nmol/L. In lysate, the input DNA template was 5 nmol/L, and the gate DNA template was 2.5 nmol/L. All input RNAs used the Thyb10 terminator.

#### 4.2 Multi-layer cascades

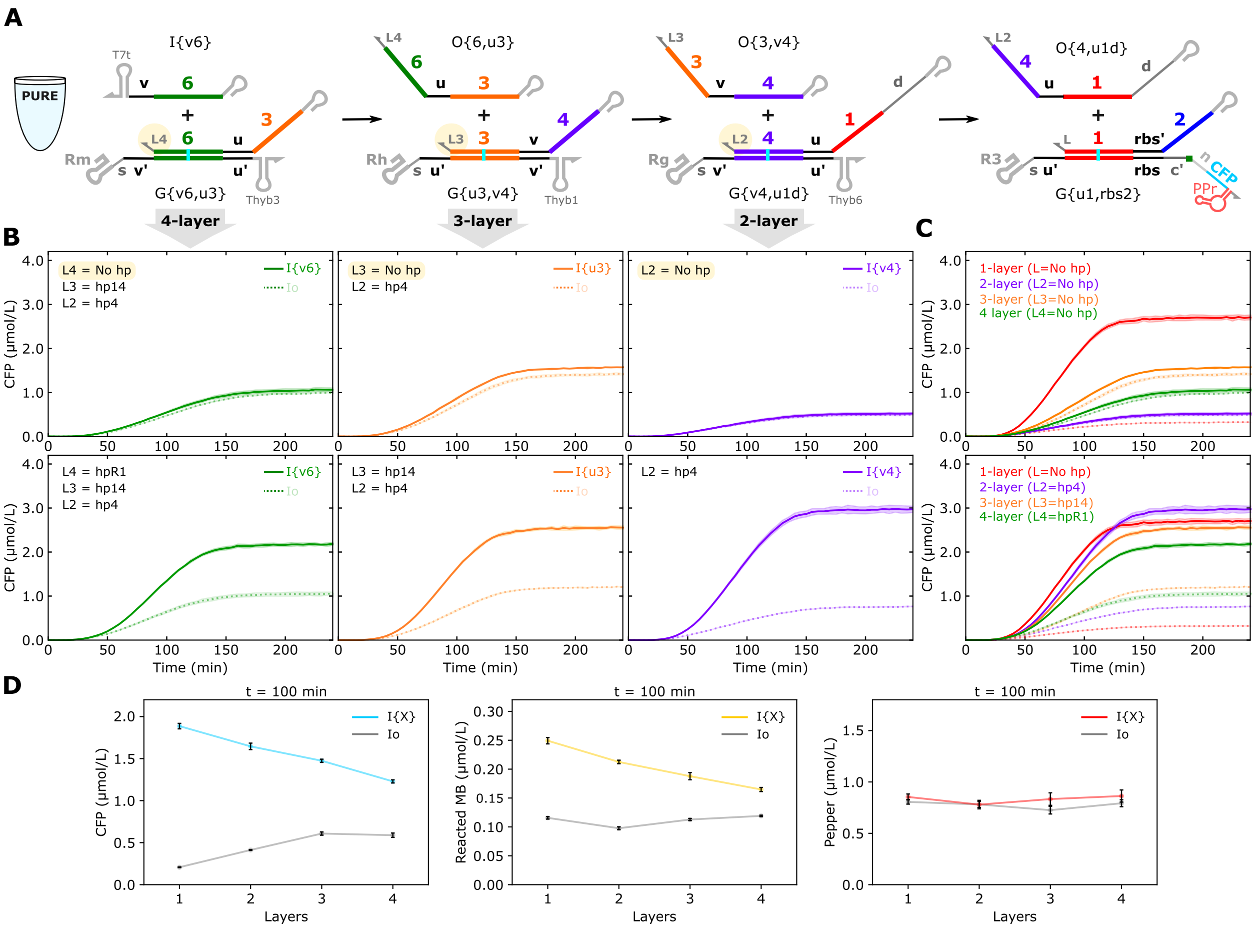

**Supplementary Figure 17**: **Characterizing multi-layer RNA strand exchange reactions in PURExpress.** (**A**) Schematic of a 4-layer RNA strand exchange cascade. Linker domains *L2*, *L3*, and *L4* are designed either to be a short 3-base linker (no hp) or a hairpin-forming linker (hpX) as specified in the plots. (**B**) CFP kinetics for multi-layer cascades without (top) and with (bottom) hairpins at the 3′ end of the output strands. Dashed lines indicate cascades expressed with a non-complementary input (Io). (**C**) Overlay of multi-layer cascade results from (B) with 1-layer results shown for comparison. (**D**) CFP, MB, and Pepper signals at 100 min as a function of cascade layers for cascades of gates with hairpins at the 3′ end of the output strands. ON/OFF ratios decrease for CFP and MB while the Pepper signal remains constant, indicating similar transcription levels for each cascade. Linear DNA templates were used for upstream gates. Io and the THE riboregulator were on plasmid templates. Input and gate DNA template concentrations were each 4 nmol/L with Io added to keep a constant 20 nmol/L total DNA template across samples. In (B) and (C), the shaded regions indicate the standard deviation of three technical replicates. In (D), the error bars indicate the standard deviation of three technical replicates.

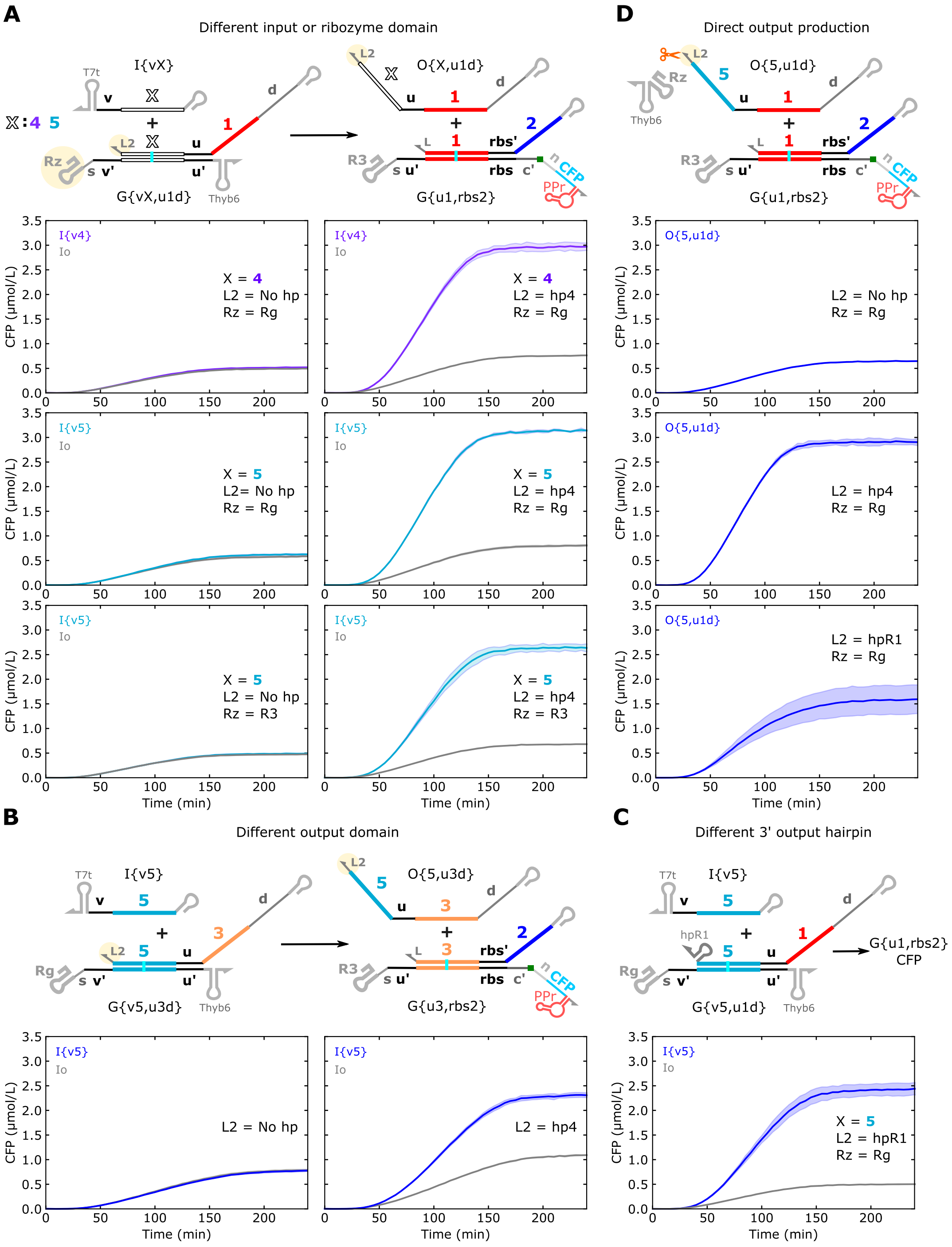

**Supplementary Figure 18:** Different 2-layer cascade gate designs, all of which need a 3′ hairpin on the output strand to produce an ON signal substantially higher than the OFF signal. CFP kinetics of 2-layer cascades with **(A)** different combinations of input domains (*4* or *5*) and intermediate gate ribozymes (Rg or R3); **(B)** a different output domain (*3* instead of *1*); **(C)** a different 3′ hairpin on the output (hpR1 instead of hp4); and **(D)** direct production of the output strand. Linear DNA templates were used for upstream gates. Io and the THE riboregulator were encoded on plasmid templates. Input and gate DNA template concentrations were each 4 nmol/L with Io added to keep a constant 20 nmol/L total DNA template across samples.

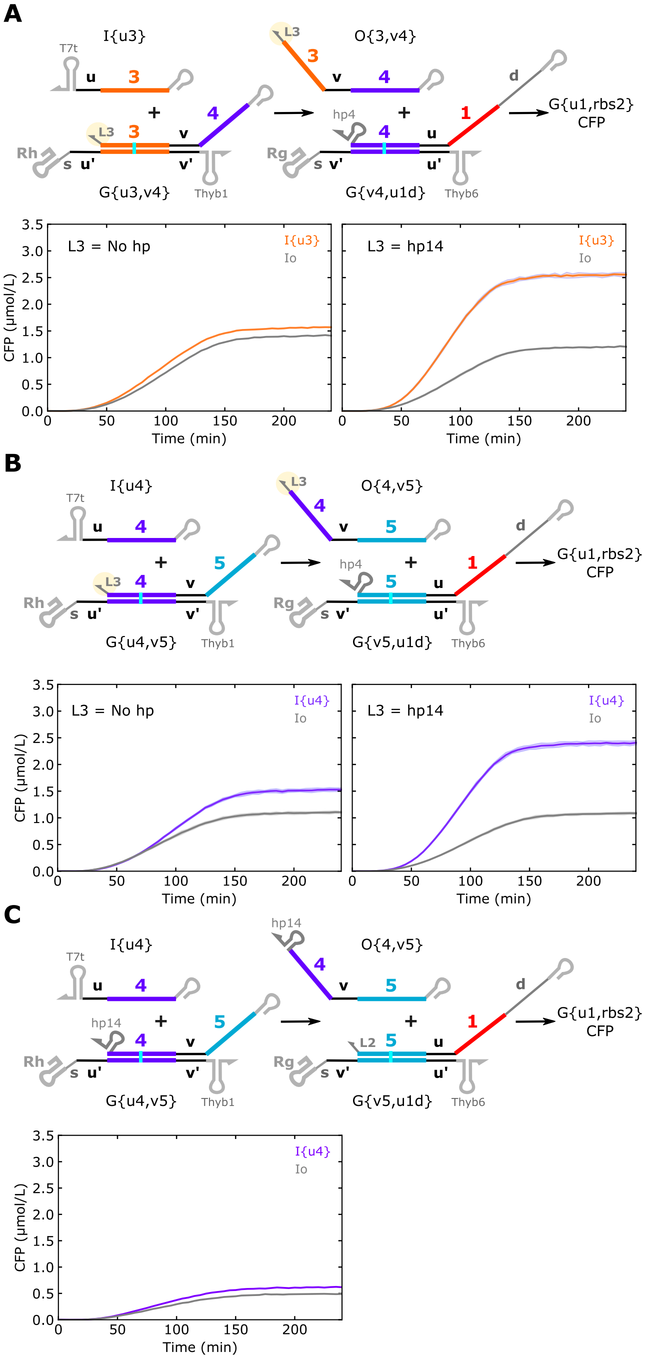

**Supplementary Figure 19:** Different 3-layer cascade gate designs, all of which need a 3′ hairpin on the output strand to produce an ON signal substantially higher than the OFF signal. CFP kinetics of 3-layer cascades with (**A**) domains *3*, *4*, and *1*; (**B**) domains *4*, *5*, and *1*; and (**C**) a gate in the middle of the cascade without a hairpin (G{5,u1d}). Linear DNA templates were used for upstream gates. Io and the THE riboregulator were encoded on plasmid templates. Input and gate DNA template concentrations were each 4 nmol/L with Io added to keep a constant 20 nmol/L total DNA template across samples.

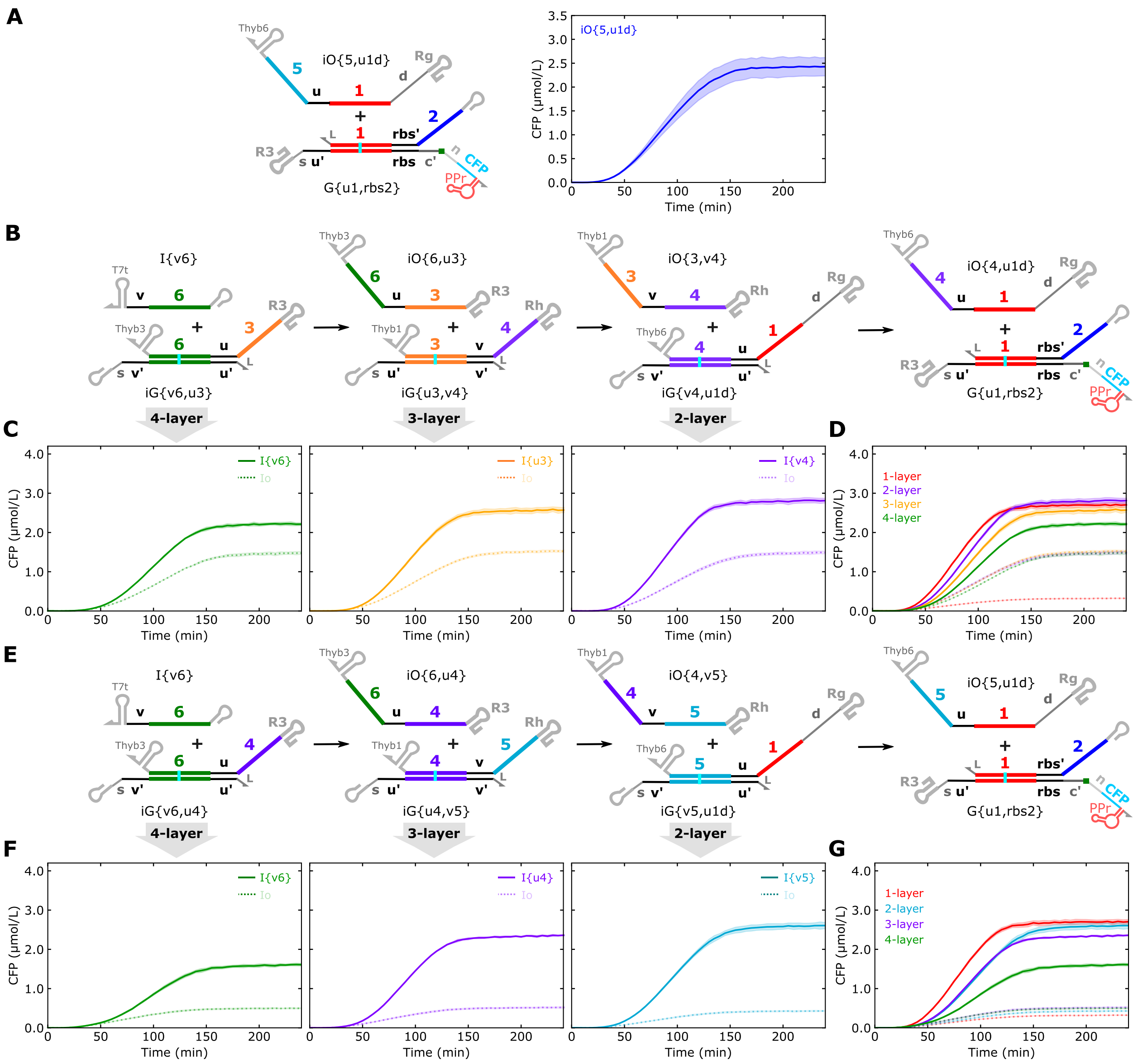

**Supplementary Figure 20:** Multi-layer RNA strand exchange reactions using gates with an inverted transcription order[4]. (**A**) Schematic and CFP kinetics of a scheme mimicking a 2-layer cascade, but with direct production of the output. In contrast with previously tested cascades, the output (iO{5,u1d}) contains a Thyb6 terminator hairpin at the 3′ end and a ribozyme at the 5′ end. The CFP yield of this scheme is similar to that with an output lacking a 5′ ribozyme (Supplementary Figure 17D). (**B**) Schematic of a 4-layer cascade with an inverted transcription order. (**C**) CFP kinetics for multi-layer cascades with components described in (B). Outputs contain a 5′ ribozyme and 3′ terminator hairpin. (**D**) Overlay of multi-layer cascade results from (C) with 1-layer results shown for comparison. (**E**) Schematic of a 4-layer cascade similar to (B), but with a different domain in the second layer (*5* instead of *4*). (**F**) CFP kinetics for multi-layer cascades with components described in (E). Linear DNA templates were used for upstream gates. Io and the THE riboregulator were on plasmid templates. Input and gate DNA template concentrations were each 4 nmol/L with Io added to keep a constant 20 nmol/L total DNA template across samples.

#### 4.3 Additional results with altered ribosome conditions

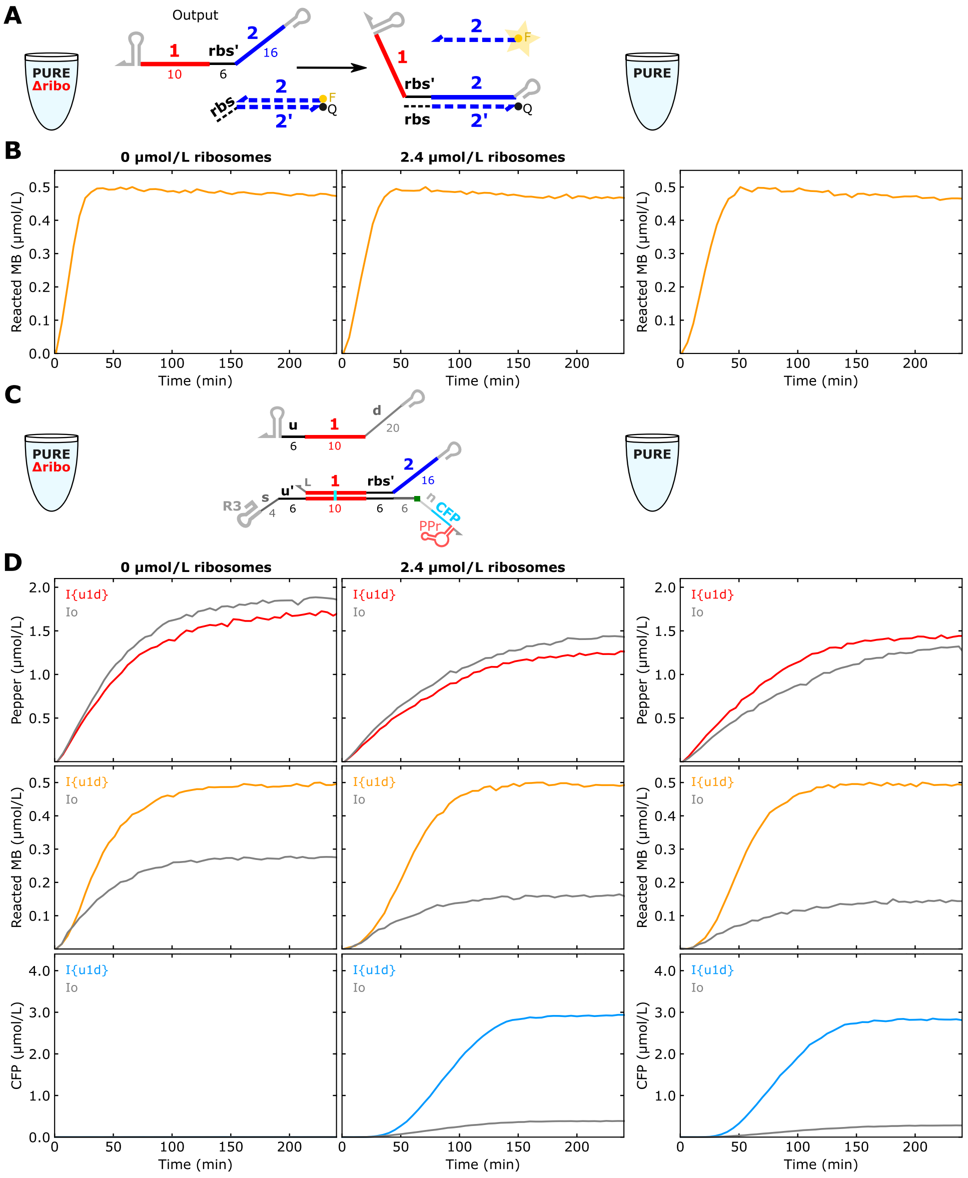

**Supplementary Figure 21:** Control templates for comparing PURExpress with and without ribosomes. (**A,B**) Testing transcription and molecular beacon measurements in PURE without ribosomes (PUREΔribo, left and center) and regular PURE (right). (**C,D**) Testing a single-layer THE riboregulator reaction. Based on molecular beacon and Pepper measurements, transcription seems to be faster in PUREΔribo than PURE. Note that results in PUREΔribo supplemented with 2.4 μmol/L produces similar results to regular PURE. DNA templates were 5 nmol/L in all experiments. All reactions had a single replicate.

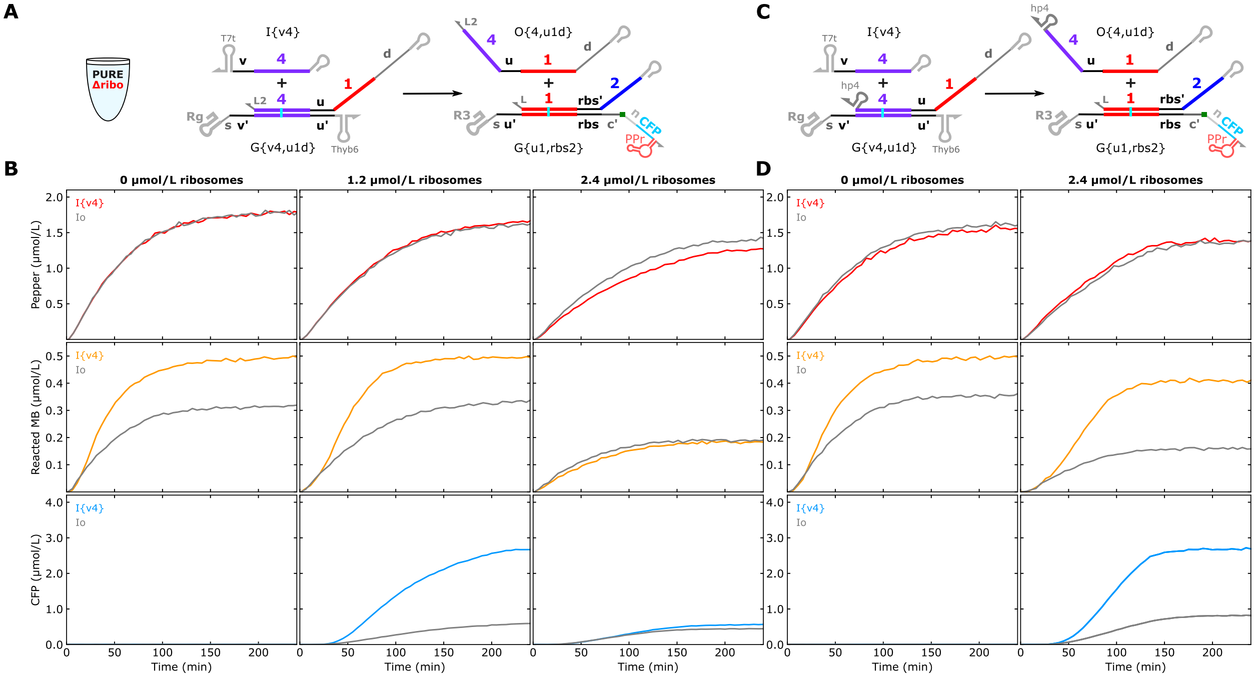

**Supplementary Figure 22:** A 2-layer RNA strand exchange cascade in the PURExpress kit lacking ribosomes (PUREΔribo) with different concentrations of ribosomes added. (**A**,**C**) 2-layer cascades using upstream gates without a 3′ hairpin (A) and with a 3′ hairpin (C) on the output strand were tested. (**B,D**) Kinetic measurements with different concentrations of added ribosomes. Input and both gate DNA template concentrations were each 4 nmol/L with Io added to keep a constant 20 nmol/L total DNA template across samples. RNA strand exchange for an upstream gate without a 3′ hairpin on the output strand is similar in the absence of ribosomes and at half the concentration of ribosomes in PURE. All reactions had a single replicate.

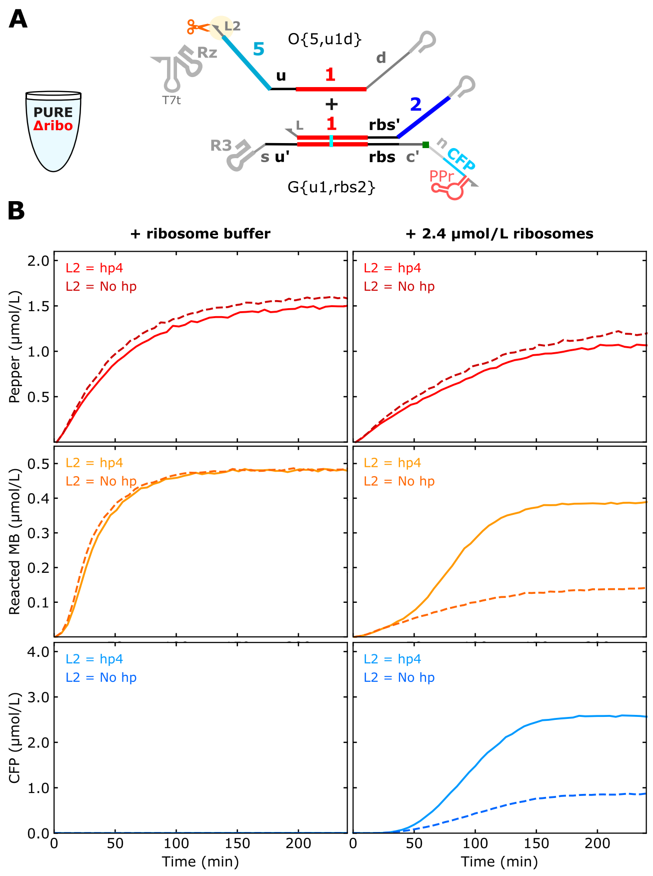

**Supplementary Figure 23:** Ribosome storage buffer does not influence results when added to PURE without ribosomes (PUREΔribo). **(A)** Schematic of components tested. Either a final concentration of 2.4 µmol/L ribosomes or the same volume of ribosome storage buffer lacking ribosomes was added to PUREΔribo. (**B**) Kinetic measurements for samples with O{5,u1d} with and without a 3′ hairpin. Output and gate DNA template concentrations were each 4 nmol/L with Io added to keep a constant 20 nmol/L total DNA template across samples. Ribosome buffer was prepared in house and composed of 20 mmol/L Hepes-KOH (pH 7.6), 10 mmol/L $Mg(OAc)_2$, 30 mmol/L KCl, and 1 mmol/L DTT. All reactions had a single replicate.

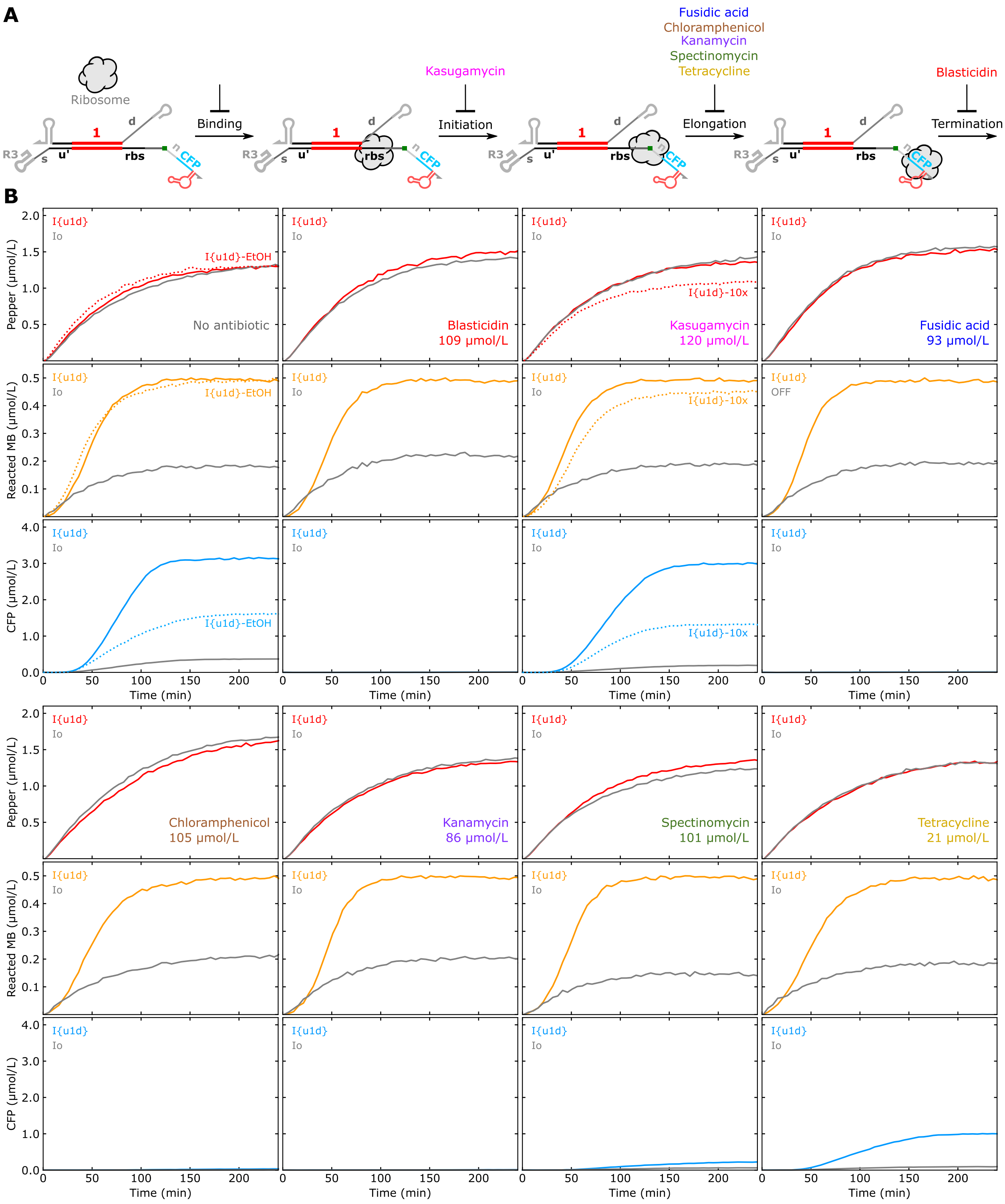

**Supplementary Figure 24:** Testing THE riboregulators in PURE with antibiotics that target different ribosome functions. (**A**) Overview of the antibiotics tested and the steps in translation they inhibit[5]. (**B**) Kinetic measurements with antibiotic concentrations shown in plots. Input (I{u1d} [], [$d_{20}$]) and gate (G{u1,rbs2} [$s_4$u], [ ], [R3]) DNA templates were both 5 nmol/L. Antibiotic stocks were prepared in water with the exception of chloramphenicol, which was prepared in 70 % by volume ethanol (EtOH). Adding EtOH to the same volume fraction as chloramphenicol was detrimental to translation (upper left plots).

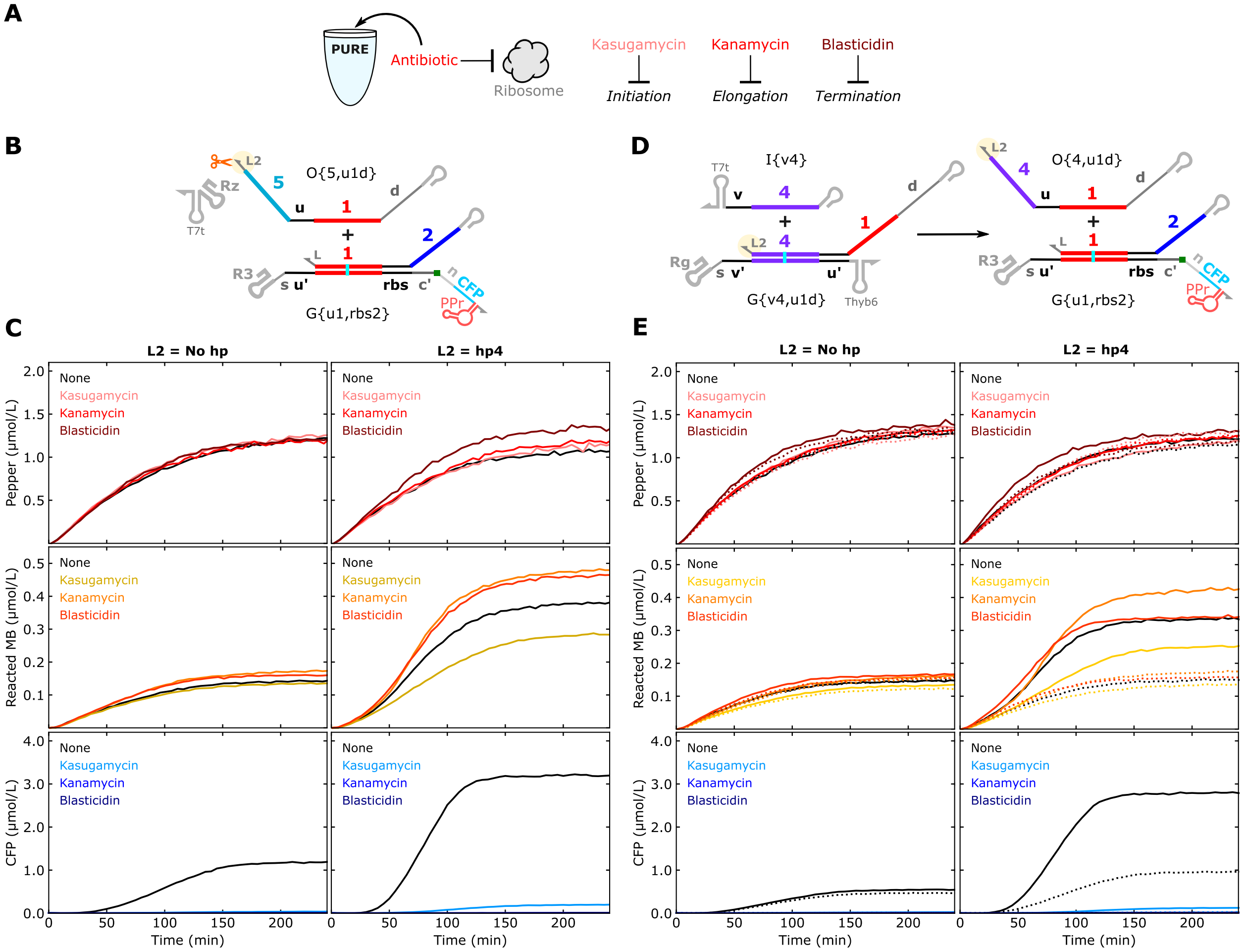

**Supplementary Figure 25**: Characterizing potential interactions of ribosomes and RNA strand exchange components. (**A**) Overview of the different antibiotics used and the mechanisms of translation that they inhibit. **(B)** Schematic of a 1-layer cascade using O{5,u1d} as the input in a PURE reaction. This system was supplemented with different antibiotics that inhibit translation. **(C)** Kinetic measurements of the system in (A) in PURE with different antibiotics. Results for O{5,u1d} without or with a 3′ hairpin are shown in the left and right columns, respectively. (**D**) A 2-layer RNA strand exchange cascade in PURE with different antibiotics that target ribosomes. (**E**) Kinetic measurements of the system in (D) in PURE with different antibiotics. Results for G{v4,u1d} without or with a 3′ hairpin for domain L2 are shown in the left and right columns, respectively. Kasugamycin, kanamycin, and blasticidin were added to a final concentration of 12.0 mmol/L, 85.8 µmol/L, and 109 µmol/L, respectively. All circuit component templates were at 4 nmol/L with Io added to maintain a total template concentration of 20 nmol/L across samples.

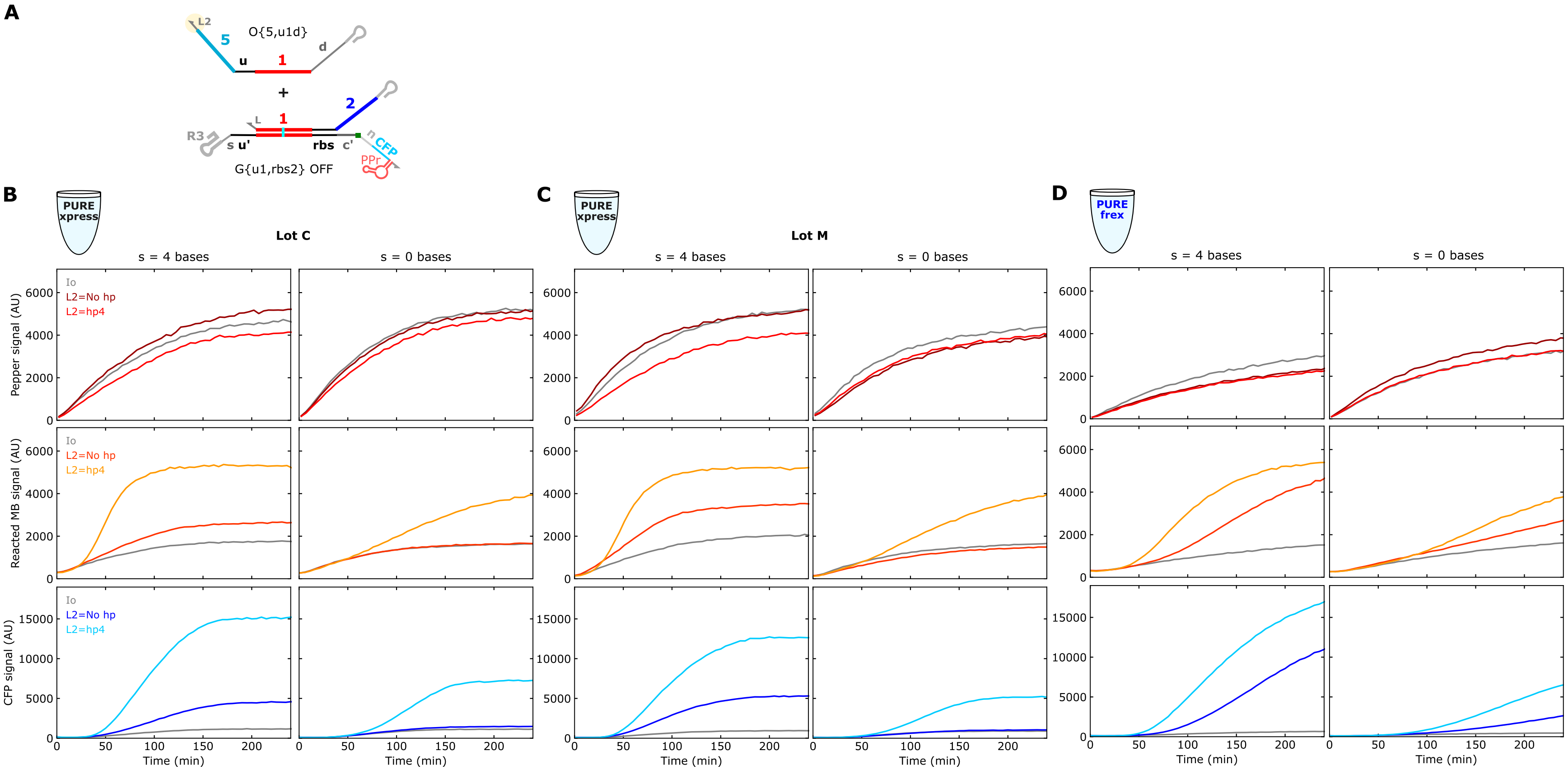

**Supplementary Figure 26**: Investigating the need for a 3′ hairpin (hp) on the triggering strand for a THE riboregulator in two lots (Lot C and Lot M) and manufacturers of PURE (PURExpress and PUREfrex). Slowing down the kinetics of the strand exchange reaction by removing the 4-base spacer (*s*) domain exacerbates the 3′ hairpin dependence across lots. Input and gate concentrations were each 4 nmol/L with Io added to keep a constant 20 nmol/L total DNA template across samples. All reactions had a single replicate.

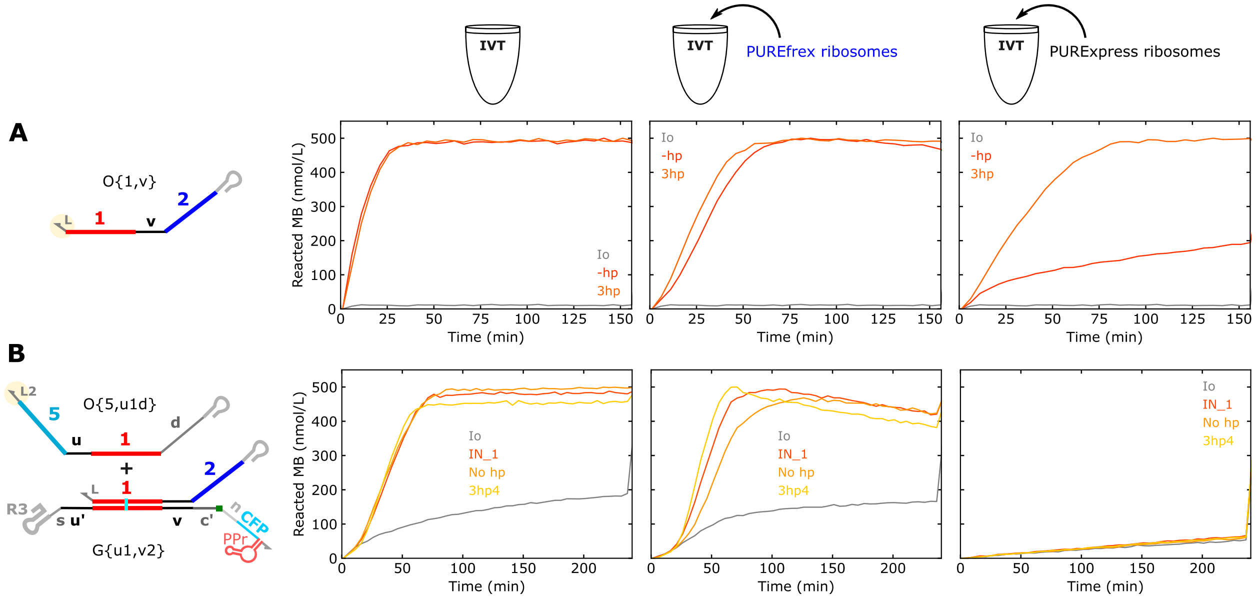

**Supplementary Figure 27**: Characterizing the effect of different ribosome preparations on IVT reactions. PUREfrex ribosomes and PURExpress ribosomes were added to IVT reactions expressing **(A)** O{1,v} or **(B)** G{u1,v2} with either O{5,u1d} or IN_1. All inputs and outputs were tested with and without a 3′ hairpin (3hp or 3hp4). In (A), The O{1,v} DNA template concentration was 5 nmol/L. In (B), O{5,u1d}, IN_1, and G{u1,v2} DNA template concentrations were each 4 nmol/L with Io added to keep a constant 20 nmol/L total DNA template across samples. All reactions had a single replicate.

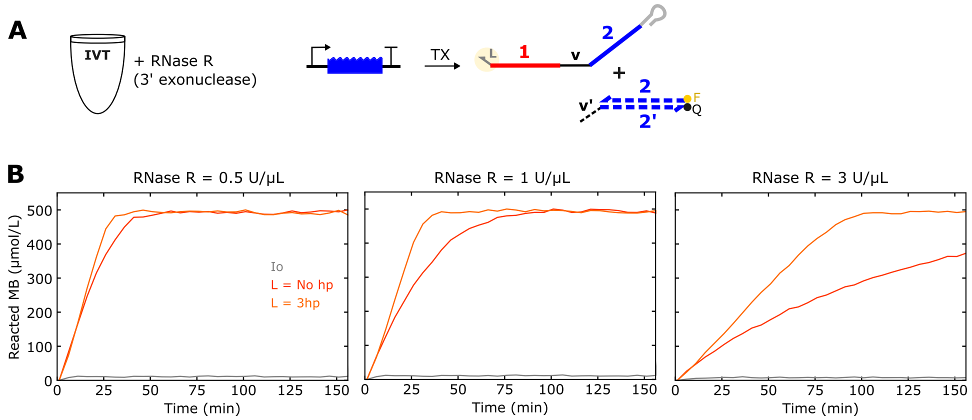

**Supplementary Figure 28**: Characterizing the effect of RNase R (New England Biolabs, cat. no. M0100S) on IVT reactions expressing output strands (O{1,v}) with or without a 3′ hairpin. The ouput DNA template concentration was 2.5 nmol/L. All reactions had a single replicate.

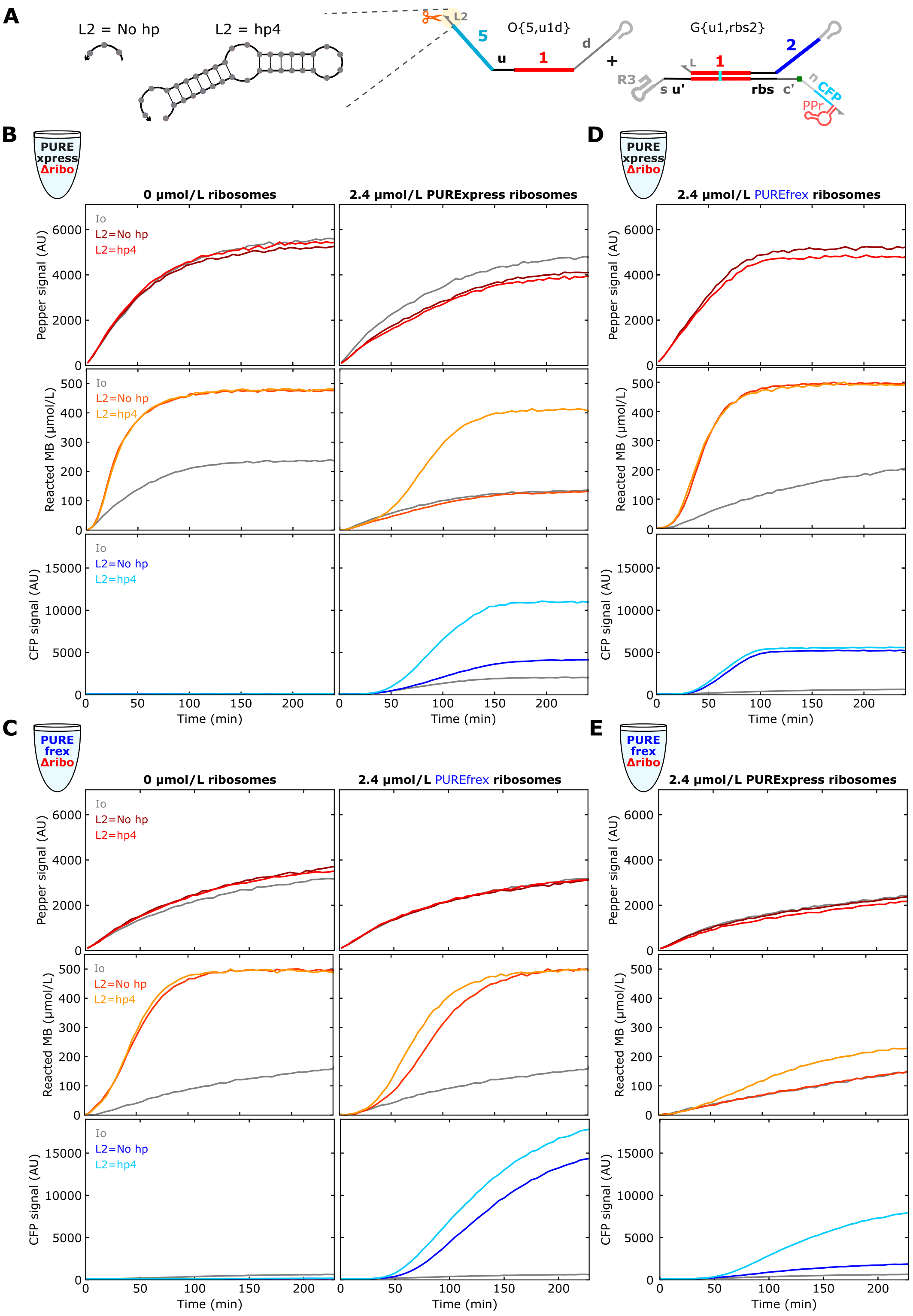

**Supplementary Figure 29**: All kinetic measurements from Figure 5 of the main text. The Pepper aptamer measurements were omitted from the main text figure for clarity.

#### 4.4 Additional results for THE riboregulator applications

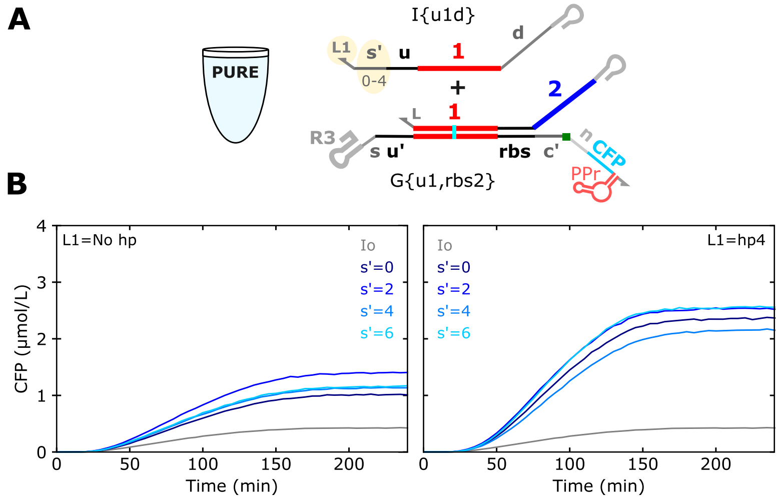

**Supplementary Figure 30**: Inputs without (left) or with (right) a 3′ hairpin (3hp4) and varying input toehold length. The length of the *s′* domain was varied to modulate strand exchange kinetics. Input and gate concentrations were each 4 nmol/L with Io added to keep a constant 20 nmol/L total DNA template across samples. All reactions had a single replicate.

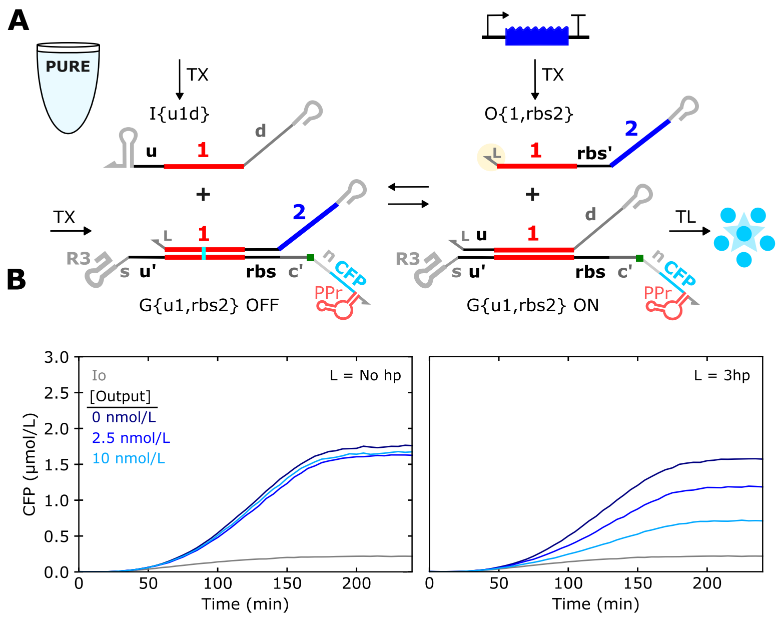

**Supplementary Figure 31**: The reversibility of THE riboregulators enables repression of protein expression. The concentration of the output, without (left) or with (right) a 3′ hairpin (3hp), was varied to modulate the extent of repression. Input and gate concentrations were each 2.5 nmol/L with Io added to keep a constant 20 nmol/L total DNA template across samples. This experiment used G_3, a gate with no spacer (*s* = 0). All reactions had a single replicate.

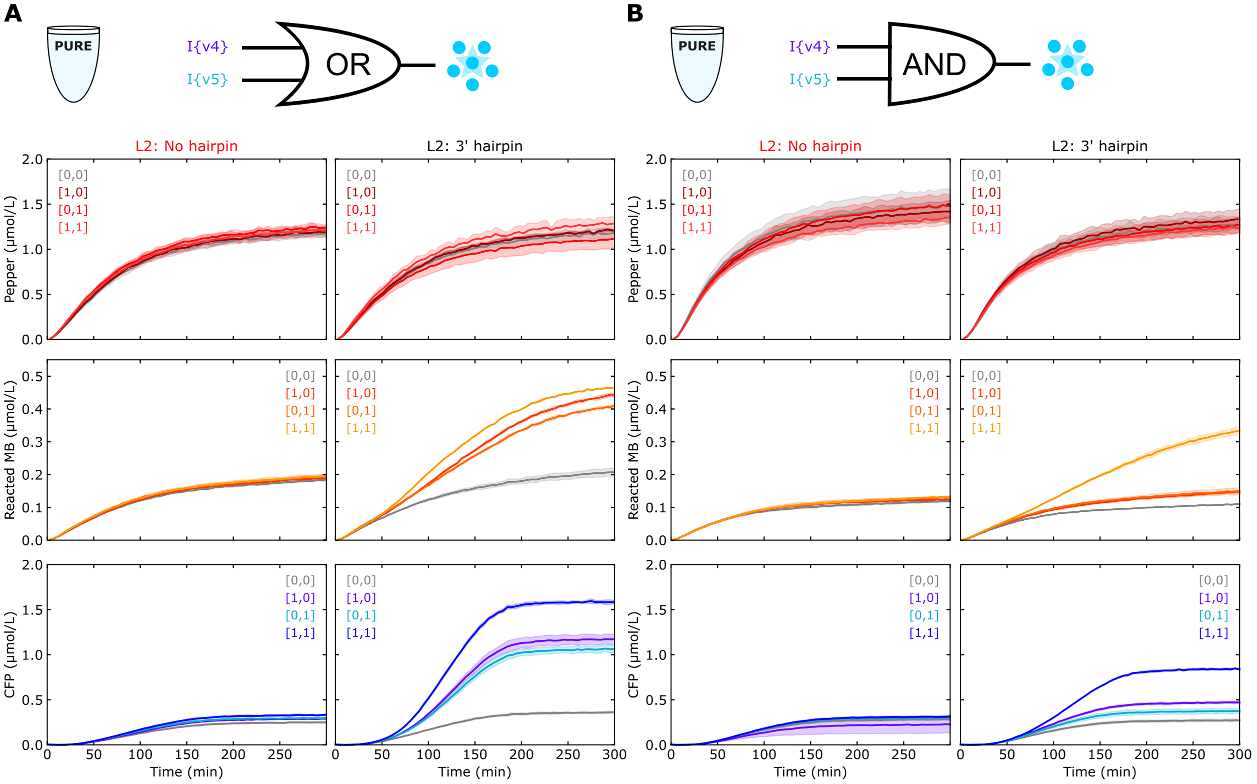

**Supplementary Figure 32:** Full kinetic measurements for the logic circuits shown in Figure 6 of the main text. The [I{v4},I{v5}] legend indicates whether I{v4} or I{v5} templates were present (1 if present in the reaction, 0 if not present).

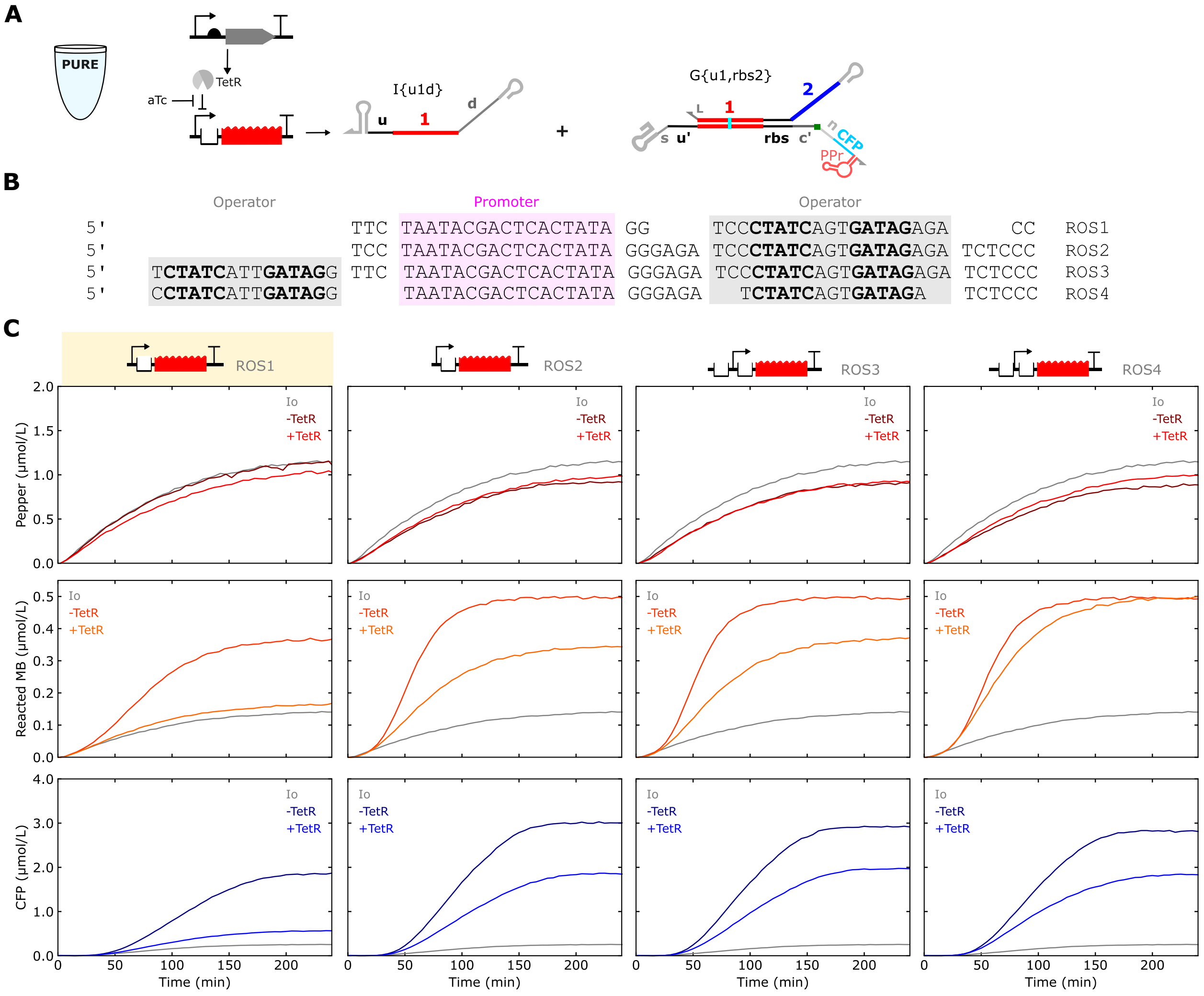

**Supplementary Figure 33:** Testing different TetR operator sequences for regulating RNA input expression. (**A**) Schematic of the system for testing operator sequences. In the absence of anhydrotetracycline (aTc), TetR should repress input transcription. (**B**) Sequence schematics of TetR operators (gray) relative to the T7 promoter sequence (pink). (**C**) TetR-mediated repression of RNA input expression with different operator sequences. In these experiments, the input DNA template concentration was 8 nmol/L, the gate DNA template concentration was 2 nmol/L, and the TetR-encoding template concentration was 4 nmol/L. Io was added to keep a constant 20 nmol/L of total template. All reactions had a single replicate.

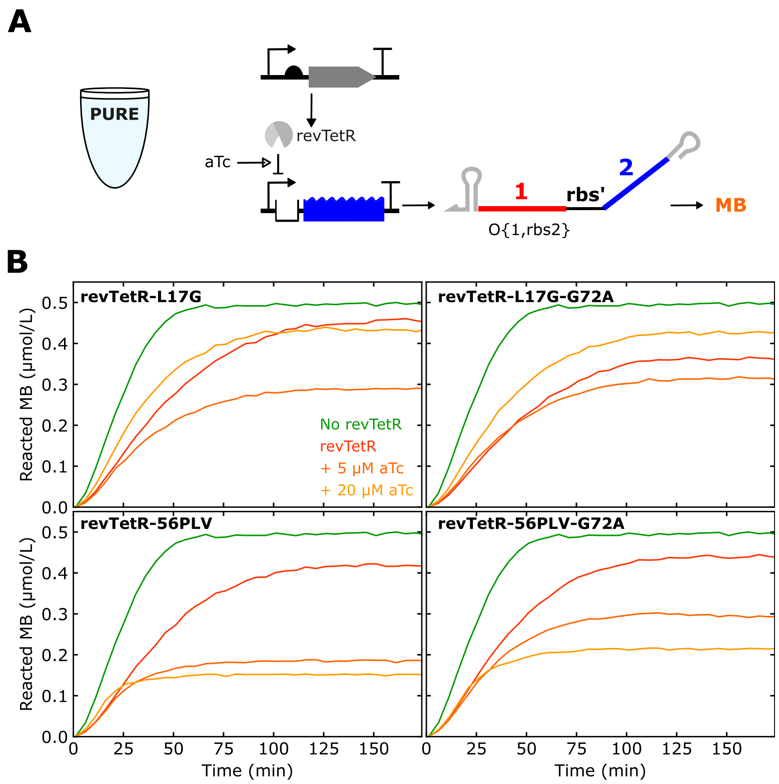

**Supplementary Figure 34:** Screening in PURE of TetR mutants (revTetR) with reverse function relative to wild-type TetR. These mutants were reported in previous studies[6] to repress transcription in the presence of anhydrotetracycline (aTc). **(A)** Schematic of the system for screening TetR mutants. O{1,rbs2} includes a TetR operator. **(B)** TetR-mediated transcription repression at two aTc concentrations. DNA concentrations were 4 nmol/L for the template encoding a TetR mutant and 2 nmol/L for the template encoding the output strand. All reactions had a single replicate.

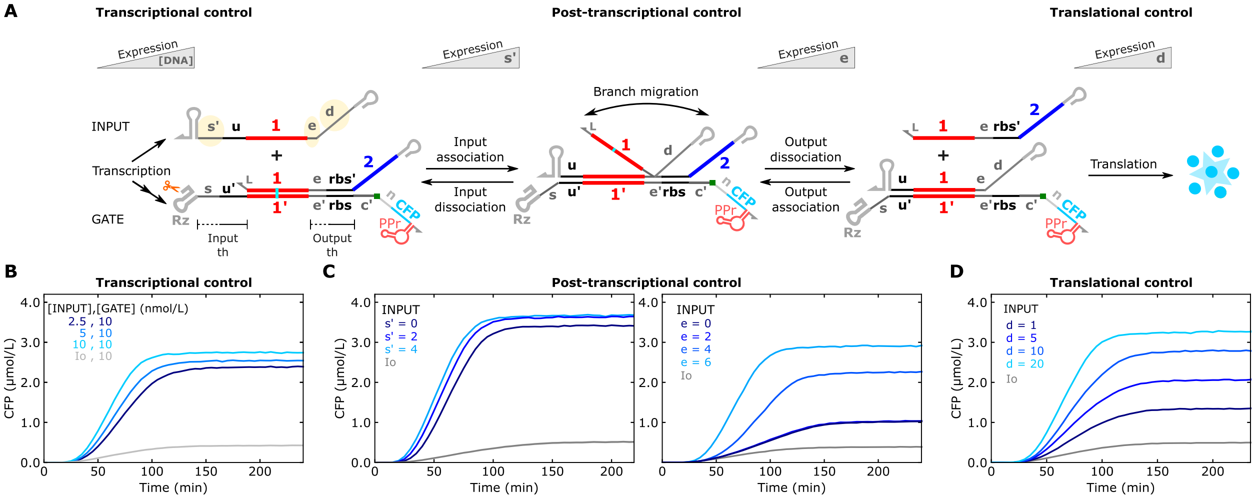

**Supplementary Figure 35: Mult-level protein expression regulation in PURExpress.** (**A**) Schematic illustrating multi-level regulation of protein expression with THE riboregulators. The yellow highlighted domains on the input RNA specify domains that can be modified to control protein expression strength at the transcriptional level (**B**), post-transcriptional level (**C**), or translational level (**D**). Results in (B), (C), and (D) are from Supplementary Figure 13, Figure 4C and D, and Figure 3, respectively.

### 5 Characterization of RNA strand exchange components with different 3′ UTRs

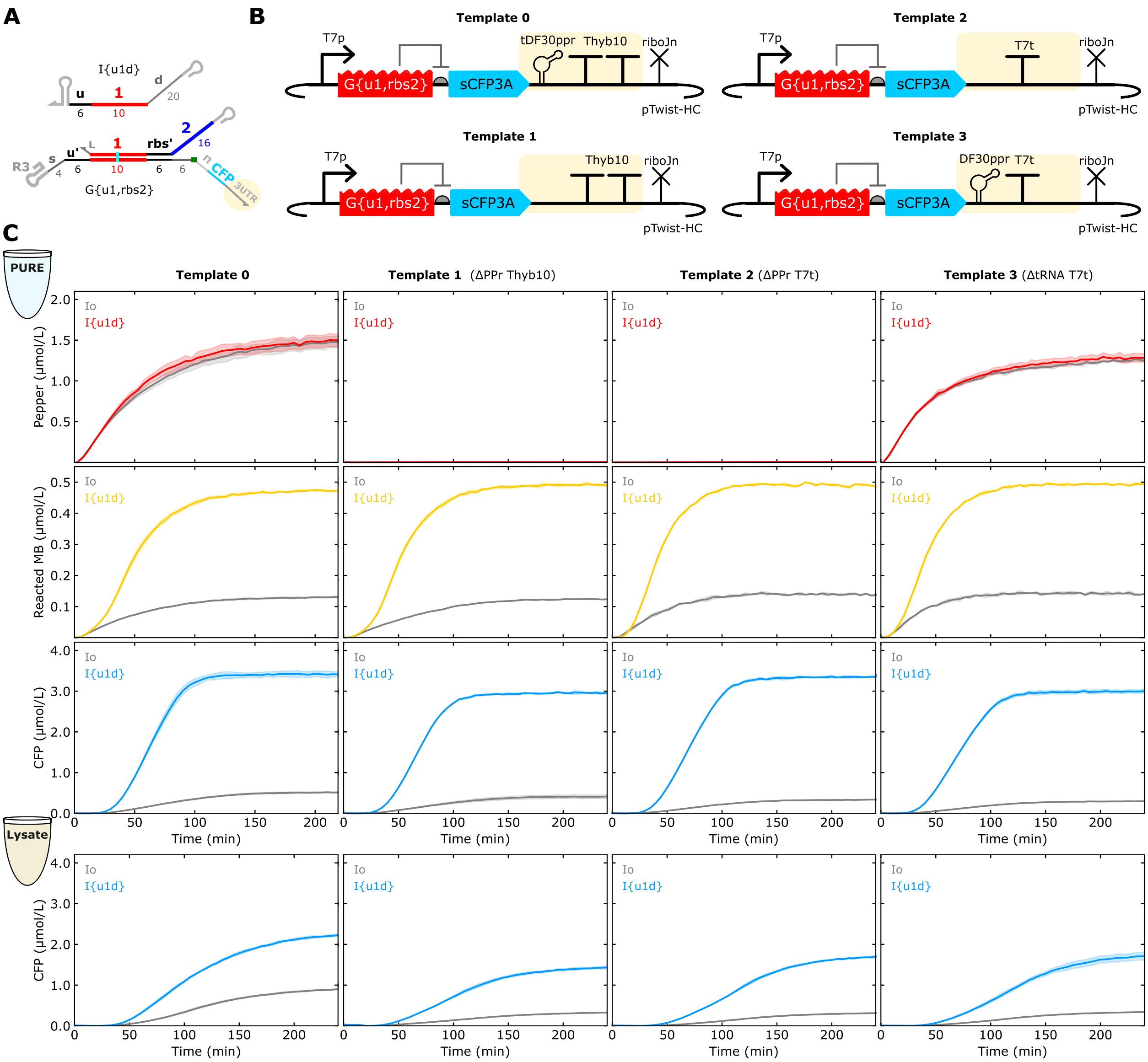

**Supporting Figure 36:** Gate 3′ UTR sequences influence results in CFES. (**A**) Schematic of RNA components tested. All inputs used the Thyb10 terminator. (**B**) Schematics of the DNA template designs for the gates with different 3′ UTRs. Template 0 is the primary design used in the study. Template 1 lacks the Pepper aptamer construct. Template 2 lacks the Pepper construct and uses the wild-type T7 terminator (T7t). Template 3 uses T7t and a Pepper aptamer construct that lacks the tRNA scaffold (DF30ppr). (**C**) Kinetic measurements in PURE and lysate for the different input and gate combinations. In PURE, the input and gate DNA templates were both 5 nmol/L. In lysate, the input DNA template was 5 nmol/L, and the gate DNA template was 2.5 nmol/L. Template 0 results are also shown in Figure 4C,D of the main text.

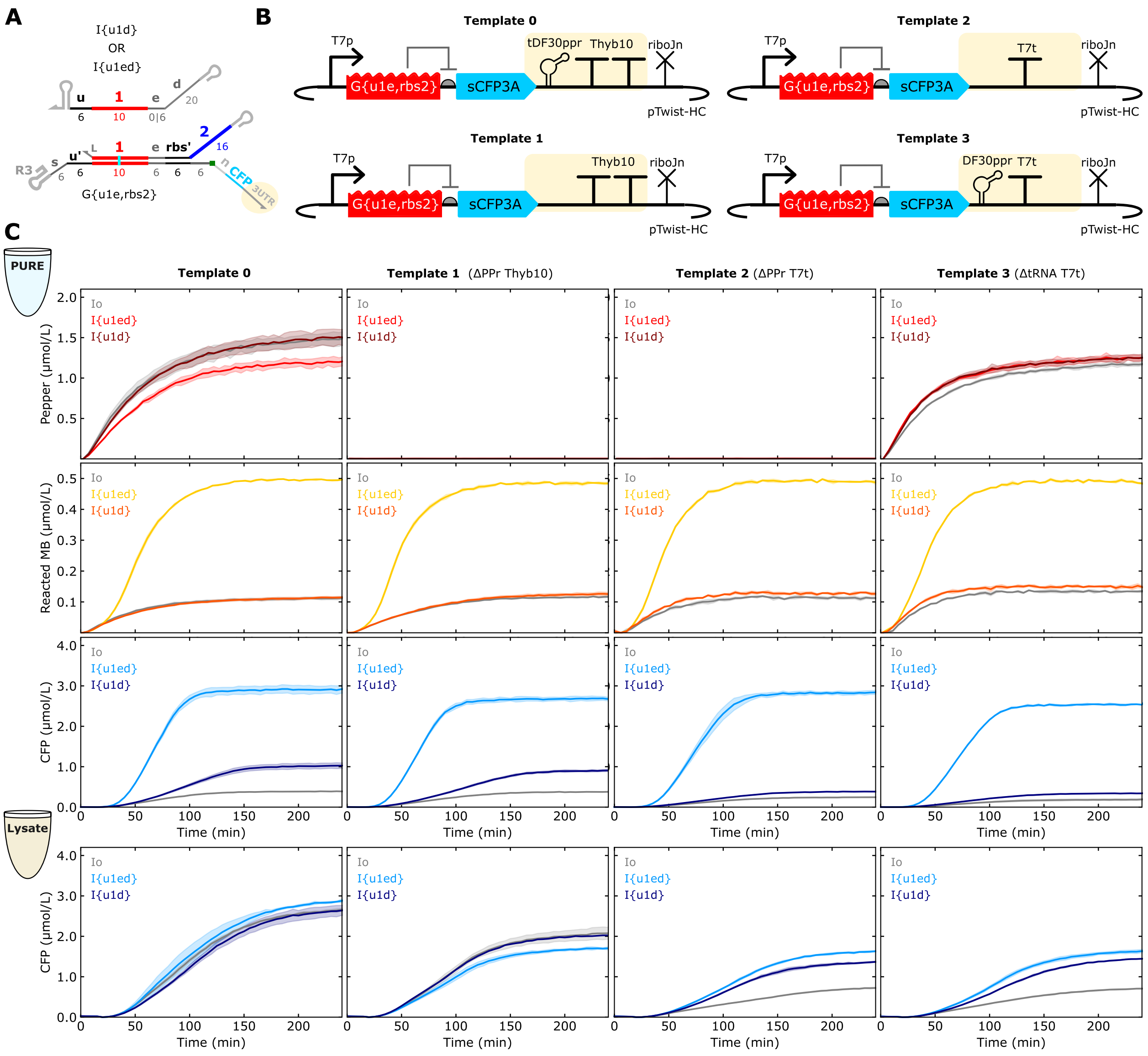

**Supporting Figure 37:** 3′ UTR sequences influence results for gates with extended branch migration domains in CFES. (**A**) Schematic of RNA components tested. All inputs used the Thyb10 terminator. (**B**) Schematics of the DNA template designs for the gates with different 3′ UTRs. See Supplementary Figure 36 for details. (**C**) Kinetic measurements in PURE and lysate for the different input and gate combinations. These results indicate the terminator sequence is the main source of the difference in leak observed in lysate for gates with extended branch migration domains. In PURE, the input and gate DNA templates were both 5 nmol/L. In lysate, the input DNA template was 5 nmol/L, and the gate DNA template was 2.5 nmol/L. Template 0 and template 2 results are also shown in Figure 4F,G of the main text.

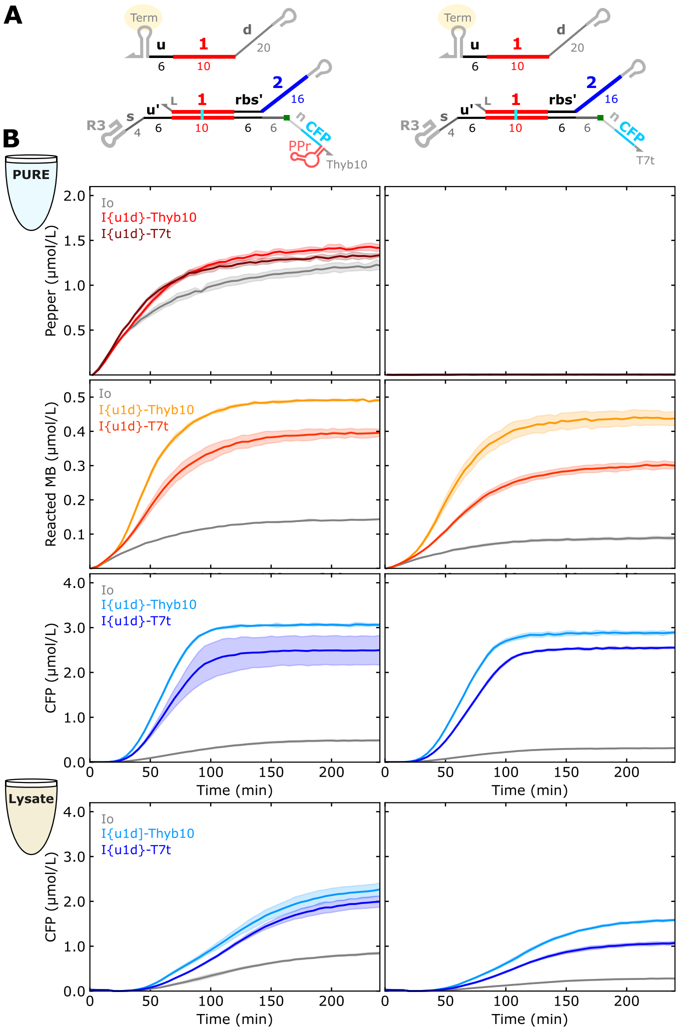

**Supporting Figure 38:** Input 3′ UTR sequences influence results in CFES. (**A**) Inputs with either the Thyb10 terminator or the wild-type T7 terminator (T7t) were tested against two different gate designs. Io used the Thyb10 terminator. (**B**) Kinetic measurements in PURE and lysate for the different input and gate combinations. In PURE, the input and gate DNA templates were both 5 nmol/L. In lysate, the input DNA template was 5 nmol/L, and the gate DNA template was 2.5 nmol/L. In PURE, differences in expression levels may be due to differences in termination efficiency, with Thyb10 likely being more efficient[3]. In lysate, differences in expression could also be related to differences in stability as Thyb10 has two hairpins.

#### 6 Characterization of toehold exchange riboregulators with different output toehold

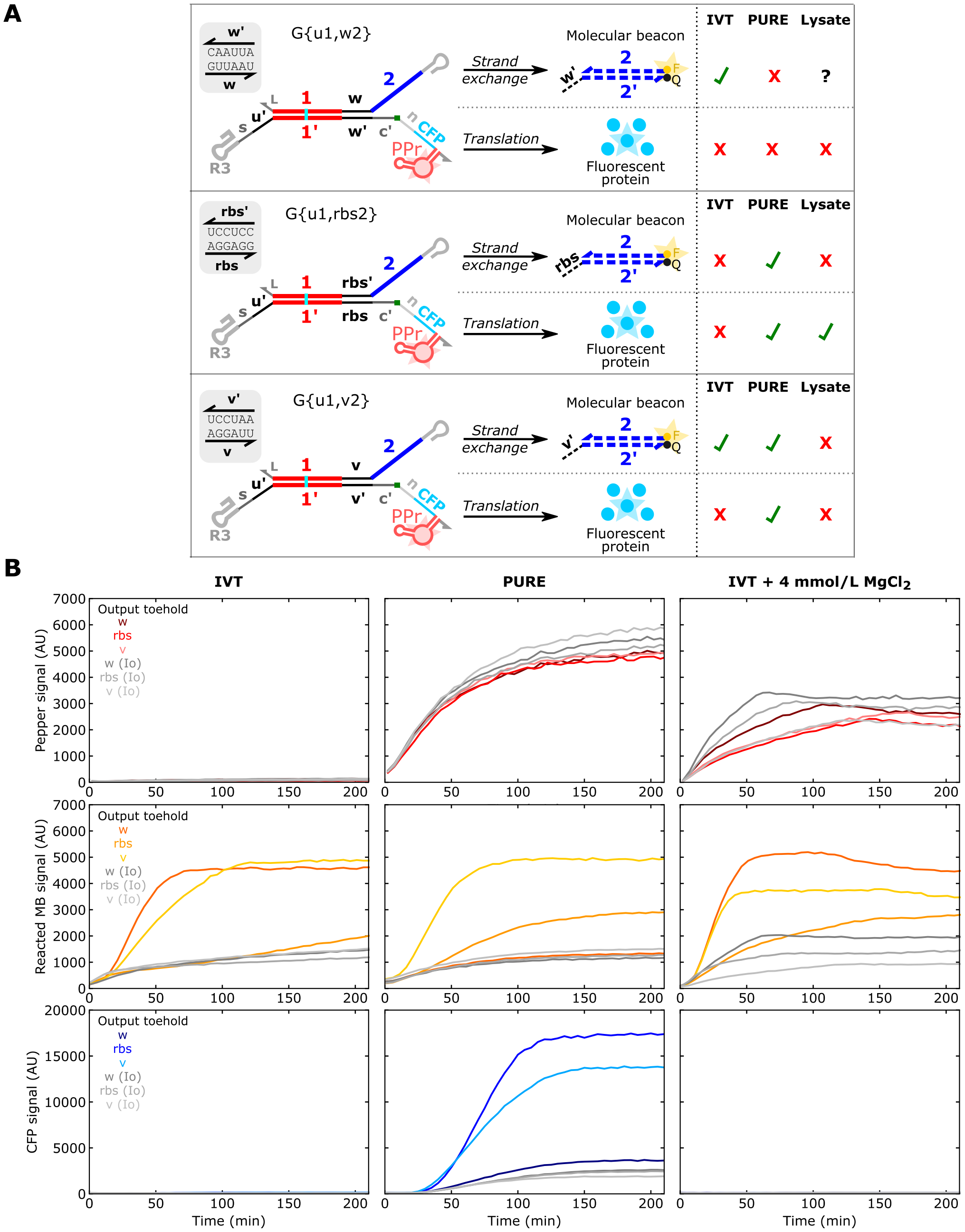

**Supplementary Figure 39**: THE riboregulators with different output toeholds. (**A**) Schematics of gates with different output toeholds and a summary of whether the two strand exchange measurements—molecular beacon and CFP—work in IVT and the two CFES, with 'X' indicating low signals and '✓' indicating high signals. In addition to the *rbs'* output toehold, two sequences *w* and *v* were tested. *v'* could serve as a weaker ribosome binding site in the ON gate and *w'* should not serve as a ribosome binding site. In these experiments, gates were expressed from linear templates, each gate had the $s_6u$ spacer (*s*) sequence, and the Pepper construct lacked the tRNA scaffold (DF30ppr). (**B**) Kinetic measurements of gates with different output toeholds in IVT and PURE. Base IVT conditions are not conducive for measuring Pepper signal (left). Supplementing IVT with an additional 4 mmol/L of $Mg^{2+}$ (2 mmol/L excess of NTPs) enables measurement of Pepper signal in IVT (right). Colored lines indicate gates cotranscribed with their complementary inputs (I{u1d}), gray lines indicate gates cotranscribed with non-complementary inputs (Io). Gate and input template concentrations were 5 nmol/L each. For IVT reactions, 2 U/μL of T7 RNAP was used.

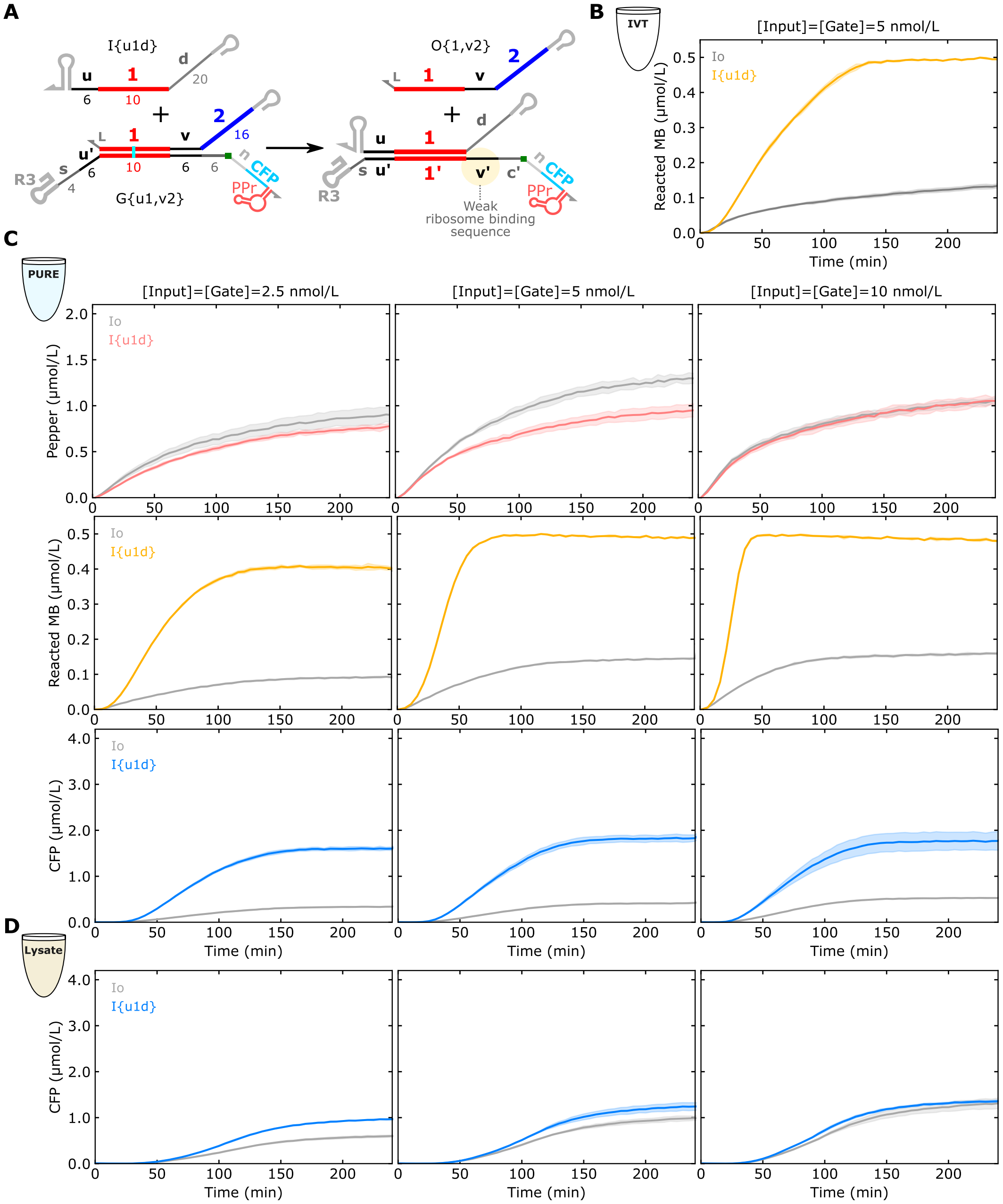

**Supplementary Figure 40**: Measurements of a THE riboregulator with a weaker ribosome binding site sequence for an output toehold (*v′*). (**A**) Schematics of a gate encoding the *v:v′* output toehold (G{u1,v2}), and its complementary input (I{u1d}). (**B-D**) Kinetic measurements in (**B**) IVT with 5 U/µL of T7 RNAP, (**C**) PURE, and (**D**) lysate at multiple input and gate concentrations (concentrations shown above plots). The *v* toehold works well in IVT and PURE, but has low expression in lysate.

### 7 Calibrating measurements to concentrations

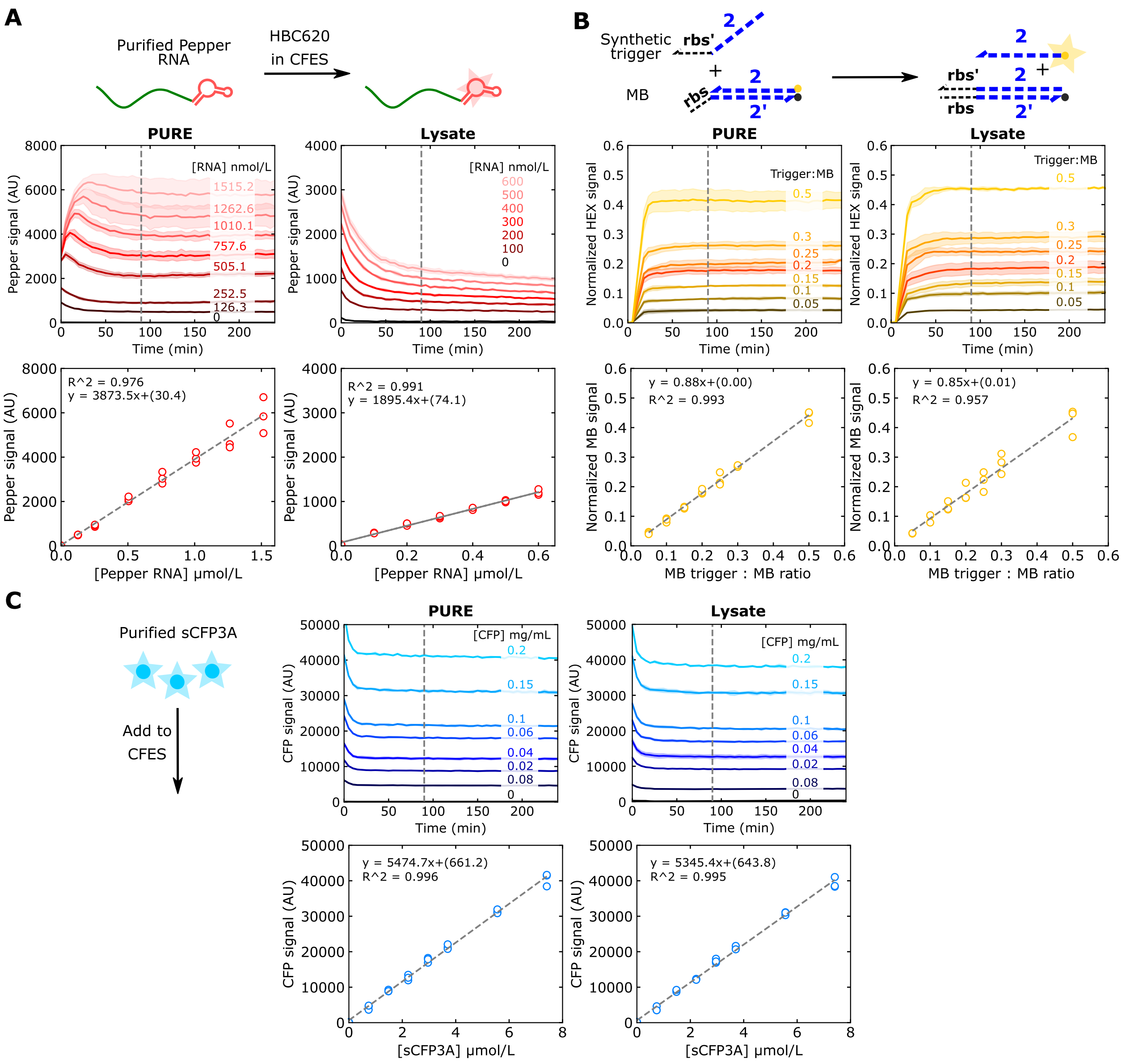

**Supplementary Figure 41**: Calibration curves for Pepper, molecular beacon, and CFP measurements to convert fluorescence intensity values in arbitrary units to concentrations of the relevant molecule. For each measurement, the relevant molecule was added to cell-free reactions in PURE and lysate at multiple concentrations capturing fluorescence intensity values above and below those measured by our experiments. Calibration curves for each CFES were generated based on the signals at 90 min and linear regression was implemented. The slope of these lines was used to correlate fluorescence signal to concentration. The y-intercept of the linear regression was not used because it is highly sensitive to experimental noise and can often results in physically unrealistic negative concentrations. Instead, the fluorescence measurement corresponding to 0 nmol/L concentrations was determined on an experiment-to-experiment basis (see Methods).

**Supplementary Table 1**: Variation in calibration curve slopes at different time points. The calibration curve slopes at 90 min were used for converting to concentration in this study and the '% changed' columns refer to the change in slope relative to the slope determined at 90 min. The CFP calibration curve slope changes less than 4 % from 1 h to 4 h. For PURE, the Pepper calibration curve slope changes less than 2.5 % after 90 min but changes more dramatically if early time points are used, presumably because RNA:dye formation is equilibrating. RNA degradation in lysate presumably causes the calibration curve slope to steadily decrease with time, making exact concentration calibration in this environment challenging. For consistency we used the slope at 90 min for RNA concentration calibration in lysate; the resulting values should not be considered 'true' concentrations but allow comparison of results across experiments on the same scale.

| **sCFP3A protein curve** | | | | |
|---|---|---|---|---|
| **Time (min)** | **PURE slope** | % changed | **Lysate slope** | % changed |
| 60 | 5556.6 | 1.50 | 5277.7 | -1.27 |
| 90 | 5474.7 | 0.00 | 5345.4 | 0.00 |
| 120 | 5520.6 | 0.84 | 5231.8 | -2.13 |
| 150 | 5536.1 | 1.12 | 5228.5 | -2.19 |
| 180 | 5514.9 | 0.73 | 5192.4 | -2.86 |
| 210 | 5457.3 | -0.32 | 5206.4 | -2.60 |
| 240 | 5458.5 | -0.30 | 5157.1 | -3.52 |
| **Pepper RNA curve** | | | | |
| **Time (min)** | **PURE slope** | % changed | **Lysate slope** | % changed |
| 60 | 4036.8 | 4.22 | 2183 | 15.17 |
| 90 | 3873.5 | 0.00 | 1895.4 | 0.00 |
| 120 | 3831.3 | -1.09 | 1791.9 | -5.46 |
| 150 | 3815.6 | -1.49 | 1715.6 | -9.49 |
| 180 | 3783.5 | -2.32 | 1654.9 | -12.69 |
| 210 | 3803.4 | -1.81 | 1584.5 | -16.40 |
| 240 | 3791.7 | -2.11 | 1536 | -18.96 |
