## Supplementary File S2 for "Cell-free systems enable mechanistic characterization of genetically encoded RNA strand exchange circuits for programmable protein expression, logic, and biosensing": README.txt

This folder contains the GenBank files of annotated plasmids used in this study.

The file 'THE CFES GenBank - all DNA RNA.gb' contains all the sequences in one file and can be imported plasmid editor software to batch import all sequences.
Note conversion to GenBank files caused some of the full names of plasmids to be truncated so batch import will include the short names specified in Supplementary File S1 and only part of the full names.

The other '*.gb' files are GenBank files of individual plasmids with the short names in parentheses. Conversion to GenBank files removed {} and [] and commas from the full names that come after the short names.
